# *Glycine max* polygalacturonase inhibiting protein 11 (*Gm*PGIP11) functions in the root to suppress *Heterodera glycines* parasitism

**DOI:** 10.1101/2024.04.27.591452

**Authors:** Sudha Acharya, Hallie A. Troell, Rebecca L. Billingsley, Kathy S. Lawrence, Daniel S. McKirgan, Nadim W. Alkharouf, Vincent P. Klink

## Abstract

Pathogen-secreted polygalacturonases (PGs) alter plant cell wall structure by cleaving the α- (1→4) linkages between D-galacturonic acid residues in homogalacturonan (HG), macerating the cell wall, facilitating infection. Plant PG inhibiting proteins (PGIPs) disengage pathogen PGs, impairing infection. The soybean cyst nematode, *Heterodera glycines,* obligate root parasite produces secretions, generating a multinucleate nurse cell called a syncytium, a byproduct of the merged cytoplasm of 200-250 root cells, occurring through cell wall maceration. The common cytoplasmic pool, surrounded by an intact plasma membrane, provides a source from which *H. glycines* derives nourishment but without killing the parasitized cell during a susceptible reaction. The syncytium is also the site of a naturally-occurring defense response that happens in specific *G. max* genotypes. Transcriptomic analyses of RNA isolated from the syncytium undergoing the process of defense have identified that one of the 11 *G. max PGIP*s, *GmPGIP11*, is expressed during defense. Functional transgenic analyses show roots undergoing *GmPGIP11* overexpression (OE) experience an increase in its relative transcript abundance (RTA) as compared to the *ribosomal protein 21* (*GmRPS21*) control, leading to a decrease in *H. glycines* parasitism as compared to the overexpression control. The *GmPGIP11* undergoing RNAi experiences a decrease in its RTA as compared to the *GmRPS21* control with transgenic roots experiencing an increase in *H. glycines* parasitism as compared to the RNAi control. Pathogen associated molecular pattern (PAMP) triggered immunity (PTI) and effector triggered immunity (ETI) components are shown to influence *GmPGIP11* expression while numerous agricultural crops are shown to have homologs.

## INTRODUCTION

Plant parasitic nematodes (PPNs) are agriculturally ubiquitous, devastating pathogens. Globally, PPNs cause 0.1-0.2 trillion dollars in economic losses, annually (Sasser and Freckman 1987; Chitwood, 2003). This problem makes studying PPNs valuable from both a basic biological, and agricultural standpoint.

Among the most devastating PPNs is *Heterodera glycines*, the soybean cyst nematode (SCN), found globally where soybeans are grown in more than 50 countries (Arjoune et al. 2022). *H. glycines* arrived in the U.S. in 1954, and now causes 36% of all losses by pathogens in the U.S., estimated at 1.5 billion U.S. dollars (Wrather et al. 2001; Arjoune et al. 2022). Consequently, practices have been sought to mitigate these losses through genetic, genomic, biotechnological, and various forms of agricultural management practices targeting both *G. max* and *H. glycines* (Peng et al. 2021).

Initial genetic efforts to identify resistance to *H. glycines* were performed by Ross (1958). Subsequent genetic mapping of the identified *G. max* resistance loci demonstrated the process of resistance is multigenic, with both dominant and recessive genes performing roles (reviewed in Concibido et al. <u>2004</u>; Neupane et al. 2019b). Early studies identified the recessive loci designated as *resistance to heterodera glycines* (*rhg*) *rhg1*, *rhg2*, and *rhg3* while the two dominant resistance loci were designated *Rhg4*, and *Rhg5* (Caldwell et al. <u>1960</u>; Matson and Williams <u>1965</u>; Rao-Arelli <u>1994</u>). Since these studies, some of the actual resistance genes have been identified (Matsye et al. 2012; Cooke et al. 2012; Liu et al. 2012; Hua et al. 2018). Other studies show not all *G. max* genotypes that exhibit resistance have the same genes or copy numbers of the same genes (reviewed in Concibido et al. 2004; Cook et al. 2012; Yu et al. 2016; Patil et al. 2019). For example, the two major sources of *G. max* resistance used in breeding programs for many years originated from *G. max* _[Peking/PI 548402]_ and *G. max* _[PI 88788]_. *G. max* _[PI 88788]_ resistance functions through *rhg1*, *rhg2*, *Rhg4*, and *Rhg5*; *G. max* _[Peking]_ has *rhg3*; while both *G. max* _[Peking/PI 548402]_ and *G. max* _[PI 88788]_ harbor *rhg1*, *rhg2*, and *Rhg5* (Glover et al. <u>2004</u>; reviewed in Concibido et al. <u>2004</u>). Some of these genes, for example found at the *rhg2* locus, remain to be cloned (Suzuki et al. 2020). Other sources of genetic resistance to *H. glycines* have since been identified as it is an ongoing process (Wen et al. 2017; Hua et al. 2018; Jiang et al. 2023; Almeida-Silva and Venancio, 2023).

Localized gene expression is a strategy that allows organisms to limit biological activities to specific cells during processes including development, differentiation, and defense to pathogen attack (Bell et al. 1984; Chrispeels et al. 1984; Broglie et al. 1984). The same processes are in play by parasitic nematodes that limit the expression of their parasitism genes to their various gland cells at specific times of their life cycle (Peng et al. 2012).

In combating pathogen attack, plants have a 2-tiered defense platform involving the recognition of epitopes that are produced directly or indirectly as a consequence of the interaction occurring between the plant and pathogen (Jones and Dangl, 2006). Collectively, the epitopes are called pathogen activated molecular patterns (PAMPs) (Janeway, 1989; Medzhitov and Janeway, 1997). The two defense tiers are referred to as PAMP (pattern) triggered immunity (PTI) and effector triggered immunity (ETI) (Jones and Dangl, 2006). PTI occurs through PAMP perception, performed by pattern recognition receptors (PRRs) which provides a basal level of resistance (Jones and Dangl, 2006). ETI involves the perception of a pathogen effector that, if the plant’s downstream signaling is intense enough, can lead to the sacrifice of its cell(s) through a hypersensitive response (HR) that leads to cell death, producing a boundary beyond which the pathogen cannot grow (Flor, 1971; Jones and Dangl, 2006). The PTI and ETI signaling processes exhibit cross communication that influences each other’s activity and defense function, so they are not mutually exclusive (Yi et al. 2015; Chen et al. 2017; McNeece et al. 2019; Liu et al. 2020; Yuan et al. 2021; Dongus and Parker, 2021; Lang et al. 2021; Ngou et al. 2021). This defense platform, whose many components were identified in the plant genetic model *Arabidopsis thaliana*, relates to *G. max* resistance to *H. glycines* (Pant et al. 2014; Klink et al. 2021).

In *A. thaliana*, PTI PRRs share the LRR-RLK co-receptor BRI1-associated kinase 1 (BAK1) and associate with a receptor-like cytoplasmic kinase (RLCK), BOTRYTIS INDUCED KINASE1 (BIK1), acting through mitogen activated protein kinases (MAPKs), transducing defense signals into an output response (Li and Chory, 1997; Jonak et al. 2002; MAPK Group, 2002; Hazzalin and Mahadevan, 2002; Veronese et al. 2006; Zipfel et al. 2006; Chinchilla et al. 2007; Liu et al. 2013; Ma et al. 2020; Lang et al. 2021; McNeece et al. 2019; Klink et al. 2021). However, BIK1 activity can lead to the expression of defense genes in the absence of signaling through MAPKs (Ngou et al. 2021). Furthermore, in *A. thaliana*, ETI can function through a plasma membrane (PM)-localized late embryogenesis abundant (LEA) coiled-coil nucleotide binding leucine rich repeat (CC-NB-LRR) protein known as NONRACE-SPECIFIC DISEASE RESISANCE 1 (NDR1), structurally similar to animal integrin (Century et al. 1995, 1997; Shapiro and Zhang, 2001; Coppinger et al. 2004). Integrins function as a receptor during cell surface adhesion that transduces signals (Tamkun et al. 1986; Knepper et al. 2011). *NDR1* expression, consistent with its role in ETI, can be induced by the heat stable, glycine rich protein effector harpin produced by the bacterium *Erwinia amylovora,* leading to a defense response (Wei et al. 1992). NDR1 associates with RPM1-INTERACTING PROTEIN 4 (RIN4), RESISTANCE TO PSEUDOMONAS SYRINGAE PV MACULICOLA1 (RPM1), and RESISTANCE TO PSEUDOMONAS SYRINGAE2 (RPS2) to activate defense signaling that can be impaired by pathogen effectors (Kunkel et al. 1993; Grant et al. 1995; Mackey et al. 2002; Belkhadir et al. 2004; Kim et al. 2005; Day et al. 2006; Sun et al. 2014). The pathogen effectors deactivate RIN4 through hyperphosphorylation, however, leading RPM1 and RPS2 to perceive the affected RIN4 which drives processes favoring defense responses including MAPK activation (Century et al. 1995, 1997; Desikan et al. 1998; Mackey et al. 2002, 2003; Axtell and Staskawicz, 2003; Lee et al. 2007; McNeece et al. 2019). Nematode associated molecular patterns (NAMPs), including the ascaroside Ascr#18 also activate MAPK signaling, driving a defense response (Manosalva et al. 2015). A functional transgenic analysis of the entire *Glycine max MAPK* gene family identified the scope of *MAPK*s having a defense role (McNeece et al. 2019). Furthermore, systemic signals can act through PTI and ETI, involving NONEXPRESSOR OF PR1 (NPR1) which functions as a co-transcriptional activator protein (Cao et al. 1994; Liu et al. 2020). Functioning upstream of NPR1 is the lipase ENHANCED DISEASE SUSCEPTIBILITY 1 (EDS1) (Feys et al. 2001). EDS1 obtains signaling input from PRRs including the LRR-RP Cf-4 and LRR-RP Ve1 that act through the BAK1 co-receptor, implicating PTI as the source of the defense signal (Feys et al. 2001; Dongus and Parker, 2021). NPR1 functions with its co-transcriptional activator TGA2, driving defense gene transcription, consistent with observations made for its role in combating parasitic nematode infection (Kawata et al. 1992; Zhou et al. 2000; Pant et al. 2014; Kaloshian and Teixeira, 2019; Sato et al. 2019). A hypothetical NAMP can function in *A. thaliana* through the <u>N</u>EMATODE-INDUCED <u>L</u>RR-<u>R</u>LK 1 (NILR1) protein with genetic mutants increasing *H. schachtii* parasitism by about 33% (Mendy et al. 2017). This observation is consistent with earlier experiments showing the *G. max* BIK1-6 reduces *H. glycines* parasitism by ∼90% and the identification of a NAMP that functions through MAPK, and likely through PTI (Pant et al. 2014; Manosalva et al. 2015). NILR1 was later shown to directly bind Ascr#18 (Huang et al. 2023).

An important class of pathogen-generated proteins that facilitate infection are polygalacturonases (PGs) (Phaff 1947). Pathogen-produced PGs function to break down cell wall polymers, permitting infection. PGs facilitate infection by cleaving the α-(1→4) linkages occurring between D-galacturonic acid residues in homogalacturonan (HG), causing cell separation, leading to the maceration of host tissue. PPNs have active PGs (Muse et al. 1970; Chitwood and Krusberg, 1977; Jaubert et al. 2002; Bird et al. 2009; Danchin et al. 2010). Plants secrete polygalacturonase inhibiting proteins (PGIPs) to interfere with this process, demonstrating the importance of an intact plant secretion system to the defense process (Collins et al. 2003; Matsye et al. 2012). PGIPs directly interfere with the activity of the PGs leading to the accumulation of oligogalacturonides (OGs) that then elicit defense responses (Cervone et al. 1997; Ridley et al. 2001). Therefore, the OGs serve as damage associated molecular patterns (DAMPs) (Matzinger 1994; Seong and Matzinger, 2004). In this manner, PGIPs deactivate the offending effector while also producing and amplifying a signaling cascade to further impair the pathogen (Matzinger 1994; Seong and Matzinger, 2004). PGIPs and PGs have long been known to relate to PPN interactions and plant defense in general (Mahalingam et al. 1996, 1999; Hammouch et al. 2012; Benedetti et al. 2015). Therefore, DAMPs relate to pectin modification (Nothnagle et al. 1983). Importantly, the necrotrophic fungus *Sclerotinia sclerotiorum* that causes stem rot diseases on more than 600 plant species produces an elicitor designated as PGIP-INactivating Effector 1 (SsPINE1) that directly interacts with and functionally inactivates PGIP as it macerates tissue (Wei et al. 2022).

MAPKs function downstream effectively through PTI and ETI in plant defense to many pathogens, including parasitic nematodes (Desikan et al. 1999, 2001; Trujillo et al. 2008; Pant et al. 2014; Aljaafri et al. 2017; McNeece et al. 2017, 2019; Cao et al. 2018; Alsherhi et al. 2018; Klink et al. 2021). Klink et al. (2022) presented a gene ontology (GO) analysis examining RNA seq data generated from RNA isolated from *GmMAPK3-1*-OE, *GmMAPK3-1*-RNAi, *GmMAPK3-2*-OE, and *GmMAPK3-2*-RNAi expressing roots in comparison to their respective controls. The analyses identified hundreds of genes with increased or decreased relative transcript abundances (RTAs) and provided their understood biological role(s). In those analyses, *GmMAPK3-1* and *GmMAPK3-2* induce the expression of a common set of *GmPGIPs* including *Glyma.05G123700*, *Glyma.05G123900*, and *Glyma.08G078900* (Klink et al. 2022). The transcriptomic experiments also show that *GmMAPK3-1* and *GmMAPK3-2* induce the expression of an *AtPEPR1* PRR co-receptor homolog *Glyma.10G195700* (*GmBAK1-1*), which functions in the defense response to *H. glycines* (Liu et al. 2013; Jing et al. 2020; Klink et al. 2021, 2022). This discovery is noteworthy because in *A. thaliana* PEPR1 recognizes short peptides which then activates BAK1 and BIK1 to promote defense responses through MAPK signaling (Liu et al. 2013). Furthermore, the *Pennisetum glaucum* (pearl millet) *MAPK4* (*PgMPK4*) gene product induces the expression of a *PGIP* defense gene (Melvin et al. 2015). More broadly, the top 10 GO terms enriched in the *GmMAPK3-1* and *GmMAPK3-2* OE induced gene set included GO:0005515 (protein binding), GO:0005524 (ATP binding), GO:0006355 (regulation of transcription, DNA-templated), GO:0003677 (DNA binding), GO:0008270 (zinc ion binding), GO:0016772 (transferase activity, transferring phosphorus-containing groups), GO:0006468 (protein phosphorylation), GO:0004672 (protein kinase activity), GO:0004674 (protein serine/threonine kinase activity), GO:0003700 (transcription factor activity, sequence-specific DNA binding) (Niraula et al. 2022a). In contrast, The top 10 GO terms enriched in the *GmMAPK3-1* and *GmMAPK3-2* OE suppressed set included GO:0005515 (protein binding), GO:0005524 (ATP binding), GO:0016772 (transferase activity, transferring phosphorus-containing groups), GO:0004672 (protein kinase activity), GO:0006468 (protein phosphorylation), GO:0004674 (protein serine/threonine kinase activity), GO:0006355 (regulation of transcription, DNA-templated), GO:0003824 (catalytic activity), GO:0003677 (DNA binding), GO:0004713 (protein tyrosine kinase activity) (Niraula et al. 2022a). The top 10 GO terms enriched in the *GmMAPK3-1* and *GmMAPK3-2* RNAi induced set included GO:0005524 (ATP binding), GO:0005515 (protein binding), GO:0004672 (protein kinase activity), GO:0006468 (protein phosphorylation), GO:0016772 (transferase activity, transferring phosphorus-containing groups), GO:0004674 (protein serine/threonine kinase activity), GO:0006355 (regulation of transcription, DNA-templated), GO:0043531 (ADP binding), GO:0006952 (defense response), GO:0004713 (protein tyrosine kinase activity) (Niraula et al. 2022a). The top 10 GO terms enriched in the *GmMAPK3-1* and *GmMAPK3-2* RNAi suppressed set included GO:0005515 (protein binding), GO:0003824 (catalytic activity), GO:0006355 (regulation of transcription, DNA-templated), GO:0016491 (oxidoreductase activity), GO:0003677 (DNA binding), GO:0020037 (heme binding), GO:0005506 (iron ion binding), GO:0055085 (transmembrane transport), GO:0009055 (electron carrier activity), and GO:0005524 (ATP binding) (Niraula et al. 2022a). Genes exhibiting various types of differential expression in only the *GmMAPK3-1* OE, *GmMAPK3-1* RNAi, *GmMAPK3-2* OE, and *GmMAPK3-2* RNAi were also identified in the analyses (Niraula et al. 2022a).

Efforts to understand *H. glycines* pathogenicity have relied heavily on genomics-based approaches since genetically-based approaches are currently intractable. Early efforts categorized the available *H. glycines* expressed sequence tags (ESTs) (Alkharouf et al. 2007; Ithal et al. 2007; Elling et al. 2007, 2009). The goal of these efforts was to employ targeted silencing of the *H. glycines* genes, much in the same manner observed in the model genetic free-living nematode *Caenorhabditis elegans* after injection with double stranded (ds) RNA in a process described as RNA interference (RNAi) (Fire et al. 1998; Alkharouf et al. 2007). Importantly, subsequent experiments on *C. elegans* demonstrated that feeding them with *Escherichia coli* transformed with genetic constructs that expressed the target gene as an inverted tandem repeat accomplished the same gene silencing outcome (Timmons and Fire, 1998). Consequently, experiments on *H. glycines* were designed under the hypothesis that when SCN was exposed *in vitro* to dsRNA (soaking) or fed from the *G. max* root cells expressing the target gene as a tandem inverted repeat they would also ingest the dsRNA, leading to RNAi silencing of the cognate SCN RNA much in the same way it occurs in the *C. elegans* (Fire et al. 1998; Timmons and Fire, 1998; Alkharouf et al. 2007; Klink et al. 2009). The identification, and annotation of the *H. glycines* EST sequences, based on similarity to conserved genes in the *C. elegans* genome that were lethal when mutated or silenced, led to the identification of gene sequences that when silenced resulted in significant SCN mortality rates of greater than 90% (Alkharouf et al. 2007). Related experiments designed to express the ESTs in transgenic *G. max* roots as tandem inverted repeats also led to high SCN mortality rates of greater than 90% (Klink et al. 2009). Subsequent efforts resulting in the generation of an *H. glycines* genome led to the identification of hundreds of putative parasitism genes (effectors) that when silenced impaired *H. glycines* development (Masonbrink et al. 2019; Lian et al. 2019). Among the functionally tested effectors were *H. glycines* pectate lyases shown to be expressed in the subventral esophageal glands (*Hg-pel-1*, *Hg-pel-2*). Other *H. glycines* pectate lyases included *Hg-pel-5*, also shown to accumulate in the subventral esophageal gland cells at the pre-second stage juvenile (pre J2), and early parasitic J2 stages (Peng et al. 2012). Four additional pectate lyases (*Hg-pel-3*, *Hg-pel-4*, *Hg-pel-6* and *Hg-pel-7*) functioning during the early *H. glycines* migratory phase accumulated in the subventral esophageal gland cells, strongly expressed at the egg, pre J2, and early parasitic J2 stages with RNAi of *Hg-pel-6* suppressing SCN development to significant levels (Peng et al. 2016). Two expansin-like genes (*Hg-exp-1* and *Hg-exp-2*), containing signal peptides at the N-terminal end were expressed in the subventral esophageal gland cells with RNAi silencing of *Hg-exp-1*, decreasing the numbers of J2s and females in *G. max* significantly (Zhang et al. <u>2018</u>). Two *H. glycines* lysozyme genes (*Hg-lys1*, and *Hg-lys2*) whose transcripts accumulate in their intestinal cells, were up-regulated when the J2s were exposed to the Gram positive *Bacillus thuringiensis, Bacillus subtilis* or *Staphylococcus aureus* with RNAi, causing a significant decrease in SCN survival and therefore are important for the or viability (Wang et al. 2019). The virulence effector Hg16B09 functions to enhance *G. max* susceptibility to *H. glycines* (Hu et al. 2019). An expression atlas of most *H. glycines* developmental stages except adult males identified 633 parasitism-associated genes encoding secretory proteins, and 156 genes whose encoded proteins share their highest similarities to plant or microbial proteins, some of which also contained a signal peptide (Elling et al. 2009). Related genomics approaches, including single nematode genome sequencing identified PPN effectors specific to certain individuals (Wang et al. 2014; Gardner et al. 2018; Maier et al. 2021; Masonbrink et al. 2019; 2021; Ste-Croix et al. 2023a). Other *H. glycines* effectors included the annexin Hg4F01, and the CLE effector proteins HgCLE1, and HgCLE2 (Patel et al. 2010; Wang et al. 2010). The *H. glycines* Hg19C07 effector, expressed specifically in the dorsal gland cell during parasitism, targets the <u>L</u>ike <u>A</u>U<u>X</u>1 3 (LAX3) auxin influx transporter (Lee et al. 2011). Importantly, LAX3 and the LAX3-induced PG are expressed in the developing syncytium as well as the cells that will be incorporated into the syncytium (Klink et al. 2009; Lee et al. 2011). This result meant that host PG-inhibiting proteins could be important in impairing pathogen success. The *H. glycines* Hg30C02 protein binds to an *A. thaliana* host plant β-1,3-endoglucanase to facilitate parasitism (Hammouch et al. 2012). The HgGLAND18 effector exhibits homology to an immunosuppressive effector domain of the malaria parasite *Plasmodium* spp. (Noon et al. 2016). Furthermore, the Hg-GLAND4 effector, which is transported to the *G. max* cell nucleus to bind to two different lipid transfer protein (LTP) gene sequences, resulted in suppressing their expression and facilitated *H. glycines* parasitism (Barnes et al. 2018). The Hg16B09 effector was exclusively expressed in the dorsal esophageal cells, during the parasitic-stage juveniles with its constitutive transgenic expression in *G. max* roots causing enhanced susceptibility while RNAi impaired parasitism as compared to the empty vector control (Hu et al. 2019). Effectors that impair ETI included GLAND1, GLAND9, and annexin, while PTI-targeting effectors included GLAND5, GLAND6, GLAND8, 2A05 (HgVAP2), Hg-5D06, Hg-13A06, Hg-33A09 (Pogorelko et al. 2020). In contrast, the Ascr#18 effector activated MAPK signaling, and plant defense (Manosalva et al. 2015). Other studies demonstrated the ability of *H. glycines* effectors Hg-VAP1, Hg-VAP2 (2A05), 3H07, 4D06, 4D09, 4G12, 5D06, 7E05, 10A06, 13A06, 25G01, 27D09, 28B03, 32E03, 34B08, and 33A09 to overcome the cell death function of the *G. max* Bcl-2 associated anthanogene 6 (BAG6) (Wang et al. 2020). Several potential *H. glycines* microRNA effectors have been identified as well but require functional study (Ste-Croix et al. 2023b). A number of these effectors are conserved between different plant parasitic nematode species, indicating that the strategies required to subvert the plant’s defense processes are conserved through plant evolution to some extent (Manosalva et al. 2015). However, plants with an effective defense barrier appear to possess additional components that are required to overcome the activities of the PPN effectors indicating an arms race is in place between the plant defense apparatus and the PPN parasitism array (Klink et al. 2009; Hewezi et al. 2010; Wang et al. 2014; Pokhare et al. 2020; Maier et al. 2021). While most of these studies proposed a very specific manner in which *H. glycines* effectors function to subvert the *G. max* defense processes, more general, evolutionarily conserved, and overarching processes may be targeted. For example, recent experiments demonstrated the involvement of the *G. max* circadian rhythm gene *CIRCADIAN CLOCK ASSOCIATED1* (*GmCCA1*), *GIGANTEA* (*GmGI*), *CONSTANS* (*GmCO*), and *TIMING OF CAB1* (*TOC1*) being involved and altering the normal circadian rhythm to hasten the defense response (Niraula et al. 2022b). These results indicated that *H. glycines* could broadly disarm the defense response by targeting the circadian rhythms, consistent with the observations made for other PPNs (Mishra and DiGennaro, 2020). However, the plant defense apparatus can detect the pathogen and rapidly respond to defend itself (Niraula et al. 2022b).

It is clear that PGIPs are important to plant defense processes, and a number of reports provide information on *PGIP* regulation and function. In *Phaseolus vulgaris* (common bean), *PGIP* expression is localized to specific cell types (embryo suspensor cells) (Frediani et al. 1993). *PGIP* promoter analyses in *P. vulgaris* demonstrated reporter expression under specific conditions including wounding but not elicitor treatment or pathogen infection (Devoto et al. 1998). Single amino acid (aa) substitution of *P. vulgaris* PGIPs led to new recognition abilities (Leckie et al. 1999). The crystal structure of a *P. vulgaris* PGIP provided new insights into its defense function (Di Matteo et al. 2003). *Brassica napus* (rapeseed) *PGIP*s are expressed to different levels in response to biotic, and abiotic stresses (Li et al. 2003). *P. vulgaris PGIP*s exist in complex loci, and undergo sub-functionalization, leading to specialized defense roles against different types of pathogens (fungi and insects) (D’Ovidio et al. 2004). Heterologous expression of a *Malus domesticus* (apple-“Granny Smith”) *PGIP* in *Nicotiana tabacum* (tobacco) inhibited disease development of the ascomycetes *Botryosphaeria obtuse,* and *Diaporthe ambigua* (Oelofse et al. 2006). An experimental examination of each of the *G. max* PGIP protein family members revealed that the observed protein activity detected in soybean tissues originated specifically from the *Gm*PGIP3 (D’Ovidio et al. 2006). Codon substitution models provided insight into the amino acids acting in the interplay between PGIPs and PGs by allowing for the narrowing down of amino acids that may function as the recognition sites present between the PG and PGIP (Casasoli et al. 2009). Mutating the identified PGIP binding sites to alanine led to the disruption of PG binding, thus identifying the recognition sites (Casasoli et al. 2009). *PGIP* gene families in *G. max*, and other legumes showed a birth and death phenomenon, possibly explaining some of the findings made in the codon substitution models (Kalunke et al. 2014). The transgenic expression of *GmPGIP* enhanced resistance to different diseases when expressed in *Triticum aestivum* (wheat) (Wang et al. 2015). A *Beta vulgaris* (sugar beet) *PGIP*, when expressed in *Nicotiana benthamiana,* limited the pathogenicity of *Rhizoctonia solani*, *Fusarium solani,* and *Botrytis cinerea* whose pathogenicities are normally driven by their PGs (Li and Smigocki, 2018).

Previous work on *G. max PGIP*s (*GmPGIP*s) functionally examined them through transgenesis; including the study of *GmPGIP1*, *GmPGIP2*, *GmPGIP3*, *GmPGIP4*, and *GmPGIP7* (D’Ovidio et al. 2006; Frati et al. 2006; Kalunke et al. 2014). The examination of *GmPGIP1*, *GmPGIP2*, *GmPGIP3*, and *GmPGIP4*, performed prior to the release of the *G. max* genome, included functional pathogenic studies on the ascomycete fungi *Sclerotinia sclerotiorum* PGb, *S. sclerotiorum* PGa, *Fusarium moniliforme, F. graminearum, Botrytis aclada, B. cinerea, Aspergillus niger*, and *Colletotrichum acutatum*, (D’Ovidio et al. 2006; Frati et al. 2006). D’Ovidio et al. (2006) demonstrated *GmPGIP1*, *GmPGIP3*, and *GmPGIP4* experienced an increase in their RTAs by 8 hours after mechanical wounding. Unlike *GmPGIP1* and *GmPGIP4*, *GmPGIP3* experienced a decrease in its RTAs between 24 and 48 hours (D’Ovidio et al., 2006). In contrast to *GmPGIP1*, *GmPGIP3*, and *GmPGIP4*; *GmPGIP2* never increased its RTAs even up to 48 hours after wounding (D’Ovidio et al., 2006). D’Ovidio et al. (2006) identified an increase in RTAs for *GmPGIP1*, *GmPGIP3*, and *GmPGIP4* after *S. sclerotiorum* infection. *GmPGIP2* RTAs increased by 48 hours post infection (D’Ovidio et al. 2006). In those studies, total RNA and a ubiquitin gene were used as loading and expression controls, respectively (D’Ovidio et al. 2006). Kalunke et al. (2014) demonstrated an increase in RTAs for *GmPGIP3*, *GmPGIP5*, and *GmPGIP7* by 8 hours after infection with *S. sclerotiorum,* normalized with *GmELF1A*. Kalunke et al. (2014) demonstrated greater than a 1,000-fold increase in RTA by 48 hours after *S. sclerotiorum* infection. Kalunke et al. (2014) also demonstrated a large increase in *GmPGIP7* RTA by 48 hours after *S. sclerotiorum* infection. The reader is directed to the Kalunke et al. (2014) reference for even more detail. Regarding *Gm*PGIP activity, D’Ovidio et al. (2006) demonstrated *Gm*PGIP1, *Gm*PGIP2, and *Gm*PGIP4 have no activity on PGs of *S. sclerotiorum* PGb, *S. sclerotiorum* PGa, *F. moniliforme*, *B. aclada, A. niger, B. cinerea, C. acutatum,* and *F. graminearum. Gm*PGIP3 has activity on PGs of each of these pathogens (D’Ovidio et al. 2006). Bulk PGIPs isolated from soybean seedlings exhibited readings similar to those observed for *Gm*PGIP3 (D’Ovidio et al. 2006). A *P. vulgaris* PGIP, *Pv*PGIP2, also exhibited activity on PGs from these examined pathogens (D’Ovidio et al. 2006). Further details of these experiments can be obtained at D’Ovidio et al. (2006). Experiments on *GmPGIP3* and *GmPGIP4* also included the hemipteran insects *Lygus rugulipennis, Adelphocoris lineolatus, Orthops kalmi,* and *Closterotomus norwegicus* (D’Ovidio et al. 2006; Frati et al. 2006). Functional studies on *GmPGIP7* included pathogenicity analyses of *S. sclerotiorum*, *F. graminearum*, *C. acutatum*, and *A. niger* (D’Ovidio et al. 2006; Kalunke et al. 2014). The analyses presented *GmPGIP1*, *GmPGIP*2, and *GmPGIP*5, also showing two remnant *GmPGIP*s on chromosome 5. Furthermore, analyses presented *GmPGIP3*, *GmPGIP4*, and *GmPGIP7* on chromosome 8 also showing two remnant *GmPGIP*s (D’Ovidio et al. 2006; Kalunke et al. 2014). The sequences were not obtained from the released soybean genome *G. max* _[Williams 82/PI 518671]_ (Schmutz et al. 2010). However, the cloning of the genes was performed using *G. max* _[Williams 82/PI 518671]_ DNA used in the original genome sequencing study (D’Ovidio et al. 2006; Schmutz et al. 2010; Kalunke et al. 2014).

The localized, cellular response of *G. max* infection by *H. glycines* has been determined through a series of histological, and electron micrographic studies. *H. glycines* burrow into *G. max* roots of either resistant or susceptible genotypes, migrating at similar rates through the host tissue (Endo, 1964, 1965, 1991).While doing so, *H. glycines* selects a cell adjacent to the vascular tissue, using its tubular, mouth-like apparatus known as a stylet to pierce the root cell to initiate the development of a feeding site from which it nurses (Endo, 1964, 1965). This event occurs approximately 2 days post inoculation (2 dpi) and initiates a process with the end goal of producing a syncytium which happens during both the susceptible and resistant reactions (Endo, 1964, 1965, 1991). During this process, the cells adjacent to the feeding site become metabolically hyperactive (Endo, 1964, 1965, 1991; Endo and Veech, 1970). Subsequently, the cell walls adjacent to the selected parasitized cell begin to dissolve. By 3 dpi, the parasitized root cell incorporates, through fusion with additional neighboring cells, eventually merging to form a syncytium (Endo, 1964, 1965, 1991; Endo and Veech, 1970). The susceptible and resistant reactions are characterized by subsequent diverse mechanisms that are evident anatomically (Endo, 1964, 1965, 1991; Endo and Veech, 1970). The susceptible reaction includes nuclei and nucleoli hypertrophy, cytoplasmic organellar proliferation, vacuolar reduction or dissolution, and cell expansion as it accumulates adjacent cells, ultimately incorporating 200-250 cells into a common cytoplasm defined as a syncytium (Endo, 1964, 1965, 1991; Endo and Veech, 1970; Gipson et al. 1971; Riggs et al. 1973; Jones and Dropkin, 1975a, b). In contrast, the potent and rapid nature of the *G. max*_[PI 548402/Peking]_ defense response was noted because most of the nematodes die early during process of parasitism at the parasitic second stage juvenile (p-J2) stage (Colgrove and Niblack 2008). The *G. max* _[PI 548402/Peking]_-type of defense response is already evident at the cellular level at 4 dpi and involves necrosis of the root cells that surround *H. glycines* head, separating the syncytium from the various other root cells that surround it (Endo 1964, 1965, 1991; Riggs et al. 1973; Kim et al. 1987; Kim and Riggs 1992). The *G. max*_[PI 548402/Peking]_-type of defense response is also defined by the presence of cell wall appositions (CWAs) (Aist,1976; Schmelzer 2002; Hardham et al. 2008). CWAs are structures defined as physical and chemical barriers to cell penetration (Aist 1976; Schmelzer 2002; Hardham et al. 2008). A second type of defense response has been observed in *G. max* _[PI 88788]_. In contrast to *G. max* _[PI 548402/Peking]_, the *G. max* _[PI 88788]_ defense response is also potent, but the response is more prolonged as characterized by the nematodes dying at the J3 or J4 stages (Acedo et al. 1984; Kim et al. 1987; Colgrove and Niblack 2008). In contrast to the *G. max*_[PI 548402/Peking]_-type of defense, the *G. max*_[PI 88788]_-type of response differs substantially by lacking a necrotic layer that, otherwise, develops around the head of the nematode (Kim et al. 1987). The initial stages of the *G. max* _[PI 88788]_-type of defense response occurs by 5 dpi and involves nuclear degeneration within the syncytium and extensive accumulation of cisternae and rough ER (Kim et al. 1987). Molecular studies confirm the existence of vesicular bodies and the expression of genes whose protein products associate with them (Klink et al. 2017; Han et al. 2023). Furthermore, the *G. max*_[PI 88788]_-type of defense response lacks the appositions that are observed in *G. max*_[PI 548402/Peking]_-type of defense. These observations were taken into consideration when selecting the studied time points in the cell-specific laser microdissection (LM) studies of the susceptible, and resistant reactions that *G. max* have to *H. glycines* parasitism attempts (Klink et al. 2007, 2009, 2010a, b, 2011, Matsye et al. 2011).

From these basic observations it can be hypothesized that large changes in *G. max* gene expression can accommodate the processes of susceptibility and resistance to *H. glycines* parasitism. Therefore, differences in RTAs will be identified when comparing gene lists found in whole root gene expression studies and studies that focus in on the syncytium and over time (Klink et al. 2007a, b). Studies of whole root RNA isolates and cell-type specific studies of the syncytium undergoing the processes of defense confirm these hypotheses (Klink et al. 2007a, b, 2009, 2010a, b, 2011; Matsye et al. 2011). Analyses also show *H. glycines* gene expression changes during its course of successful parasitism (Ithal et al. 2007). These experiments have been followed up by comparative gene expression studies of *H. glycines* experiencing a susceptible reaction in *G. max* _[PI 548402/Peking]_ through infection with *H. glycines* _[race 14/TN8/HG-type 1.3.6.7]_ or a resistant reaction in *G. max* _[PI 548402/Peking]_ through infection with *H. glycines*_[race 3/NL1-Rhg/HG-type 7]_ (Klink et al. 2009). Furthermore, RNAi analyses of *H. glycines* genes annotated to have essential functions as well as putative parasitism genes play important roles in subverting *G. max*’s ability to defend itself (Alkharouf et al. 2007; Ithal et al. 2007; Elling et al. 2007, 2009; Klink et al. 2009).

The analysis presented here extracted and updated the PGIPs existing in *G. max*_[Williams 82/PI 518671]_ that were originally sequenced by Schmutz et al. (2010). The *Gm*PGIPs were compared to those present in the genomes of other important crops and other plants. The *GmPGIP11* (*Glyma.19G145200*) is shown to be expressed in the *H. glycines*-induced syncytium, a root cell that normally functions as the nematode’s feeding site but in the *G. max*_[Peking/PI 548402]_ and *G. max*_[PI 88788]_ genotypes is the site of the defense response (Endo 1964, 1965, 1991). The *GmPGIP11* is overexpressed in the *H. glycines* susceptible *G. max*_[Williams 82/PI 518671]_ in transgenic hairy roots of chimeric, genetically-mosaic *G. max* plants as demonstrated by real time quantitative PCR (RT-qPCR). This experiment was done because it is hypothesized that *GmPGIP11* overexpression will decrease *H. glycines* parasitism in a *G. max* genotype (*G. max*_[Williams 82/PI 518671]_) that is otherwise naturally susceptible (Matsye et al. 2012). In contrast, *GmPGIP11* is cloned for RNAi in the *H. glycines*-resistant, *G. max*_[Peking/PI 548402]_, confirmed by RT-qPCR. This experiment was done to see if *GmPGIP11* RNAi increases parasitism in a *G. max* genotype (*G. max* _[Peking/PI 548402]_) that is otherwise naturally resistant to *H. glycines* parasitism (Matsye et al. 2012). The results provided a bioinformatics-base characterization of the *G. max PGIP* gene family, identified the *G. max PGIP*s whose expression was influenced by MAPKs, and showed that the *GmPGIP11* functioned in defense to *H. glycines* where previous experiments were not performed. Results are presented that show PTI and ETI defense branches regulate *GmPGIP11* RTAs. Another *G. max PGIP*, *GmPGIP1* whose expression was induced by PTI and ETI components but was not expressed within the syncytium undergoing a defense response to *H. glycines* parasitism, is shown to not function in the defense response under the study conditions. Bioinformatics analyses compared the 3-dimensional (3-D) structures of *Gm*PGIP11 and *Gm*PGIP1 to identify any potential differences that may be related to the results obtained in the *H. glycines* parasitism studies as revealed by the female indexes (Fis). Further comparisons are made to two *Beta vulgaris* (sugar beet) *PGIP*s that are expressed during defense to another root pathogen, the sugar beet root maggot (SBRM) *Tetanops myopaeformis*, and function in defense responses to different pathogens (Li and Smigocki et al. 2016; Alkharouf et al. 2024). The results provide promising evidence that *GmPGIP*s can function in defense to other root pathogens (Shah et al. 2017). A conglomerate gene expression atlas highlights the expression of the 11 *G. max PGIP*s and 112 PGs extracted from numerous publications that span the resistant and susceptible reactions of different populations of *H. glycines* as well as localized expression occurring in resistant and susceptible syncytia Vaghchhipawala et al. 2001; Alkharouf et al. 2004; Khan et al. 2004; Tucker et al. 2007; Klink et al. 2005, 2007b, 2009, 2010a, b, 2011; Ithal et al. 2007; Puthoff et al. 2007; Afzal et al. 2009; Tucker et al. 2011; Matsye et al. 2011; Wan et al. 2015; Li et al. 2018; Song et al. 2019; Neupane et al. 2019a; Guo et al. 2020; Miraeiz et al. 2020; Shi et al. 2021; Kofsky et al. 2021; Chu et al. 2022; Torabi et al. 2023).

## MATERIALS AND METHODS

### General statement of materials and methods

The materials and methods used in this study employ procedures found in Klink et al. (2021), and the cited references. Other cited references include Niraula et al. (2022a, b); Klink et al. (2022), and Khatri et al. (2022). Otherwise presented here, details of the bioinformatics analyses can be found in Acharya et al. (2023).

### PGIP identification

The PGIP proteins presented for the targeted plant species were identified through a Basic Local Alignment Search Tool program (BLAST) search of their proteomes (BLASTP) that used the default parameters at Phytozome (http://www.phytozome.net/) (Altschul et al. 1990; Goodstein et al. 2012). The querying included: Target type: Proteome; Program: BLASTP-protein query to protein database; Expect (E) threshold: -1; Comparison matrix: BLOcks SUbstitution Matrix (BLOSUM) 62 (BLOSUM62); Word (W) length: default = 3; number of alignments to show: 100 that allowed for gaps and filter query, in the order that they appear on the BLAST program. Therefore, through these analyses it was possible to extract the genomic DNA, transcript, cDNA, protein accessions, their sequences, and gene family members. The presented analyses permitted the extraction of protein homologs and splice variants from the selected agricultural crops of international importance, importance in the U.S., and those that are important biologically according to Klink et al. (2021; Acharya et al. 2023). The analyzed proteome list is provided (Goodstein et al. 2012) (**Supplemental Table 1**).

To obtain the protein sequences used in this analysis, the identified PGIP proteins for the studied crops were compiled using a cutoff of a Bitscore of 140. To identify the PGIP proteins, each of the 11 *G. max* PGIP protein sequences were queried into the proteomes of the studied, agriculturally important crops and other plant species. The individual queries for *Gm*PGIP1 through *Gm*PGIP11 were stored in individual tabs in Excel. Then, the PGIPs with Bitscores of 140 or higher were compiled for all of the queries for the individual *Gm*PGIPs. The duplicate PGIPs were removed in Excel. The analysis resulted in a list of PGIP proteins that included the products of alternate splicing so the protein numbers in some cases were higher than the numbers of genes in some genomes.

### Signal peptide and glycosylation prediction

Signal peptide prediction was done using SignalP - 6.0 in the Eukarya option set to the default settings (Teufel et al. 2022). *N*-linked glycosylation was predicted using NetNGlyc - 1.0 set to the default settings (Gupta and Brunak, 2002). *O*-linked glycosylation was predicted using NetOGlyc - 4.0 set to the default settings (Steentoft et al. 2013).

### Artificial intelligence

Prediction of eukaryotic protein subcellular localization was done using the deep learning program DeepLoc-1.0 (Almagro Armenteros et al. 2017). The DeepLoc-1.0 analysis determined the importance of a particular aa along a protein chain relevant for prediction (attention) of its subcellular location. The analyses were done under default settings. DeepLoc-1.0 then predicted the subcellular localization of eukaryotic proteins, differentiating between 10 different localizations including the nucleus, cytoplasm, extracellular, mitochondrion, cell membrane, endoplasmic reticulum, chloroplast, Golgi apparatus, lysosome/vacuole, and peroxisome. The analyses were done using default settings.

### 3-dimensional modeling of PGIP proteins

SWISS-MODEL (https://swissmodel.expasy.org/) was used for homology modeling of the protein sequences. The SWISS-MODEL Workspace is a personal web-based working environment, where several modelling projects can be carried out in parallel (Waterhouse et al. 2018). In addition to that, protein structure models can be compared by superimposing them over each other. The output was obtained using the default parameters.

### Pairwise protein sequence comparison

The PGIP pairwise comparison was made using NEEDLE of EMBOSS under default settings. The Needleman-Wunsch algorithm was employed (Needleman and Wunsch, 1970). The employed matrix was EBLOSUM62 (Madeira et al. 2019).

### Cell type-specific *PGIP* gene expression studies

A previously performed transcriptomic analysis was used to facilitate the selection of the *G. max PGIP* (*GmPGIP*) gene family members for their use in the transgenic studies (Matsye et al. 2011, 2012). The original transcriptomic analysis involved the isolation of RNA from root cells that underwent LM. To increase the confidence that the observed *GmPGIP* gene expression was consistent in nature, the LM-assisted analyses were done independently in triplicate using two different *H. glycines*-resistant *G. max* genotypes (see below). The analyses presented here were done to identify the *GmPGIP*s that were expressed consistently in the same manner at the same time points in each genotype. The 2 *H. glycines*-resistant genotypes, *G. max*_[Peking/PI 548402]_ and *G. max*_[PI 88788]_, were infected with *H. glycines*_[NL1-Rhg/HG-type 7/race 3]_. This procedure generated a defense response in each genotype (Matsye et al. 2011). These roots were processed for paraffin-embedding and subsequent histology, and also LM on a Leica ASLMD microscope. RNA was isolated from LM-collected cells at 0 days post infection (dpi) mock-infected control cells (pericycle) that were sampled prior to the roots being infected with *H. glycines.* Subsequent to *H. glycines* invasion (infection) of the root, the parasitic nematode produces syncytia that were also sampled as they were undergoing the process of defense at 3, and 6 dpi. The reason for selecting these time points was that the 3 dpi time point occurs prior to the onset of visible (histological) defense response signs. In contrast, by the 6 dpi time point the defense response was clearly differentiated from the susceptible reaction, cytologically. RNA was isolated from the laser microdissected cells using the PicoPure RNA Isolation kit (Molecular Devices®) with a DNAse treatment using DNAfree® (Ambion®) added just before the second column wash, according the manufacturer’s instructions. RNA yield and quality determination, using the RNA 6000 Pico Assay® (Agilent Technologies®) in the Agilent 2100 Bioanalyzer®, was done according to the manufacturer’s instructions. The cDNA probe preparation and its hybridization to the probe sets on the Affymetrix® Soybean GeneChip® were performed according to Affymetrix® guidelines (Affymetrix®). The experiment was run using probe generated from each of the *G. max* _[Peking/PI 548402]_ and *G. max* _[PI 88788]_ genotypes in triplicate for a total of 6 microarrays studied for each time point. The Bioconductor implementation of the standard Affymetrix® DCM method was used in the gene expression analysis. The analysis identified genes that were considered expressed at a particular time point through the detection call methodology (DCM) (Klink et al. 2010b). The determination of gene expression was made if the probe signal was measurable above threshold on all three arrays for that time point (i.e., 0, 3 or 6 dpi) for both *G. max* _[Peking/PI 548402]_ and *G. max* _[PI 88788]_ (6 total arrays), p < 0.05. The standard Affymetrix® microarray DCM analysis, done in Bioconductor, consisted of 4 steps. The 4 steps included saturated probe removal, discrimination score calculation, Wilcoxon’s rank test p-value calculation, and detection call assignment. This procedure is quantitative, and determined whether the gene’s expression was provably different from zero. The analyses determined if the expression was present (P), had uncertain (marginal) measurement (G) or was not provably different (absent) from zero (A). In the analyses presented here, a candidate gene (i.e., *GmPGIP*) met the measured [M] criteria when the probe signal was detectable above threshold (p < 0.05) on all 6 microarrays for a given time point if probe signal was detected at a statistically significant level (p < 0.05) on each of the 6 microarrays using the Mann–Whitney–Wilcoxon (MWW) Rank-Sum Test (Mann and Whitney 1947). In contrast, the *GmPGIP* expression was considered not measured (NM) if probe signal was not detected at a statistically significant level (p ≥ 0.05) on any single one of the 6 microarrays using the MWW Rank-Sum Test. For the purpose of the analysis here, the MWW Rank Sum Test is a nonparametric test of the null hypothesis that does not require the assumption of normal distributions (Mann and Whitney 1947). Not all *GmPGIP* gene expression profiles were determined under the analyses parameters because some genes did not have probe sets fabricated onto the microarray. Consequently, for those *GmPGIP*s gene expression was not determined and was categorized as not applicable (n/a) to this study. Two different *G. max* genome annotations were used in obtaining the gene accession numbers. The Affymetrix microarray annotations were mapped to the original *G. max* genome release (Accession 1) Wm82.a1.v1.1 (2010) out of necessity, used at the time of publication of Matsye et al. (2011), because just that annotation was available. Subsequently, these older annotations were updated with the accessions presented here for a more recent *G. max* Wm82.a2.v1 (2015) genome assembly and annotation (Accession 2).

These studies also included a 9 dpi time point which is beyond the onset of visible signs of the resistant reaction (Endo, 1965, 1991; Klink et al. 2009, 2010a, b, 2011; Matsye et al. 2011). The 9 dpi analyses were performed at the same time and manner that the 0, 3, and 6 dpi time point were done. The data for the *Gm*PGIP gene expression were extracted and provided for the 9 dpi time point.

### *PGIP* gene cloning

The *GmPGIP* (*GmPGIP11* and *GmPGIP1*) DNA primers were designed to facilitate directional Gateway cloning (Curtis and Grossniklaus 2003). The Gateway cloning pipeline was compatible with the features of the pRAP15 (overexpression [OE]) and pRAP17 (RNA interference [RNAi]) destination vectors (Klink et al. 2009, 2021; Matsye et al. 2012). *GmPGIP* gene integration employed the CACC nucleotide sequence that was added to the 5’ end of the forward primer. This DNA sequence first promoted the incorporation of the PCR-generated amplicon into the pENTER/D-TOPO entry vector (Invitrogen) (**Supplemental Table 2**) (Klink et al. 2021). *GmPGIP* amplicons were generated by PCR and then visualized in 1% agarose gel. The *GmPGIP* amplicons were purified by the Wizard® SV Gel and PCR Clean-Up System and protocol (Promega). *GmPGIP* amplicons were ligated directionally into the pENTER/D-TOPO vector and protocol (Invitrogen). The reaction contents were transformed into the chemically competent *E. coli* One Shot TOP10 (TOP10) strain by protocol (Invitrogen). Transformed TOP10 were chemically selected on LB-kanamycin (LB-kan) at 50 μg/ml on agar plates by protocol (Invitrogen). Plasmids were isolated by Wizard® SV minipreps by protocol (Promega). *GmPGIP* gene sequences were confirmed against the original *G. max* Phytozome *GmPGIP* accession. The *GmPGIP* amplicons contained in the pENTER/D-TOPO vector was transferred into the pRAP15 and pRAP17 destination vectors through an LR clonase reaction by protocol (Invitrogen). The reaction produced the recombinant pRAP15 and pRAP17 destination vector plasmids for overexpression and RNAi of *GmPGIP1 and GmPGIP11,* respectively. The experimental controls were the un-engineered pRAP15 or pRAP17 vectors that contain the *ccd*B gene located in place where, otherwise, the targeted *GmPGIP* amplicon was engineered during the LR clonase reaction. The un-engineered pRAP15-*ccd*B (OE control) and pRAP17-*ccd*B (RNAi control) vectors, therefore, were appropriate controls to account for any non-specific effects caused by candidate resistance gene (CRG) OE or RNAi (Pant et al. 2014). The 301-bp figwort mosaic virus (FMV) sub-genomic transcript (Sgt) promoter fragment [sequence −270 to +31 from the transcription start site (TSS)] drives the transcription of the *GmPGIP*, as well as the pRAP15-*ccd*B, and pRAP17*-ccd*B control OE and RNAi cassettes respectfully. Enhanced green fluorescent protein (e*GFP*) gene expression in both pRAP15 and pRAP17 plasmids were driven by the *rol*D promoter. Agar plates containing LB-tetracycline (LB-tet), 5 μg/ml, were used to select for TOP10 bacteria that had the vectors that contained the integrated *GmPGIP* amplicon. The isolated *GmPGIP*-containing destination vectors were transformed to chemically competent *Agrobacterium rhizogenes* strain K599 (K599) that employed the freeze-thaw method with colony selection that occurred on LB-tet agar (5 μg/ml) (Matsye et al. 2012). The transformed K599 culture was incubated for 16 h at 28° C before performing a vacuum infiltration transformation to produce transgenic *G. max* roots.

### Genetic engineering of *G. max*

Genetic transformation began by using six-day old *G. max* plants already grown in the greenhouse in sterilized sand at ambient temperatures. The growth conditions included a 16h day/8h night cycle with supplemental light provided by cool white, fluorescent lights (Sylvania 21781 FO32/841/ECO T8, 32 Watt, 4100 Kelvin, 2950 Lumens 48 inch tube bulbs, color rendering index [CRI] of 85). For K599 transformed with the pRAP15 OE vector (*GmPGIP*-containing or *ccd*B control-containing), the bacteria were used to genetically engineer into *H. glycines*-susceptible genotype *G. max*_[Williams 82/PI 518671]_. For K599 transformed with the pRAP17 RNAi vector (*GmPGIP*-containing or *ccd*B control-containing), the genetic cassettes were genetically engineered into the *H. glycines* -resistant *G. max* _[Peking/PI548402]_. Notably, roots genetically engineered using the K599 containing the applicable pRAP15-*ccd*B (OE) or pRAP17-*ccd*B (RNAi) “empty” vector served as their respective, controls.

A 250-ml culture of K599 genetically transformed with the *GmPGIP*-containing plasmid or respective control was pelleted by centrifugation in a Sorvall RC6 Plus Superspeed Centrifuge at 4 C for 20 min. The K599 pellet was resuspended in Murashige and Skoog medium including vitamins (MS) (Duchefa Biochemie) that contained 3.0% sucrose, pH 5.7 (MS media). The *GmPGIP* genes were genetically engineered in the *H. glycines* -susceptible genotype *G. max*_[Williams 82/PI 518671]_ for overexpression. In contrast, the *GmPGIP* genes were genetically engineered in the *H. glycines* -resistant genotype *G. max* _[Peking/PI 548402]_ for RNAi. The pRAP15-*ccd*B-OE and pRAP17-*ccd*B RNAi controls were generated in the same manner. Transgenic *G. max* plants were made by cutting off the root of a 6 day-old plant at the hypocotyl using a new, sterile razor blade. The rootless plants were then immersed in the K599 solution in MS media in a Petri dish prior to cutting. This step provided the genetically transformed K599 immediate access into the wound site made by the removal of the root. Then, 25 rootless plants were placed in a 140-ml glass beaker containing 25 ml of transformed K599 in MS media solution. Infiltration of the plant tissue, with cut ends in the genetically transformed K599 occurred under vacuum generated using the VP60 Two Stage Vacuum Pump (CPS Products, Inc.) in a Bel-Art “Space Saver” polycarbonate vacuum desiccator with a clear polycarbonate bottom (Bel-Art) for 5 min. The pump was turned off after 5 min. Subsequently, the stopcock was closed, and the plants were left in the vacuum for an additional 10 min. The vacuum then was slowly released. The slow release of the vacuum allowed the transformed K599 further entry into the plant tissue. After the vacuum was completely released, the cut ends of *G. max* were placed individually into fresh coarse Vermiculite Grade A-3 (Palmetto Vermiculite) in 50-cell 725602C Propagation Trays (T.O. Plastics). The propagation trays were held in 710245C Standard Flats that had holes in the bottom (T.O. Plastics). Holes that were 1 cm wide made with a glass cylinder and at a depth of 3–4 cm deep were made for the rootless plants. The plant trays were then placed in a Sterilite, 25 Qt/23 L Modular Latch Box (Sterilite). The container was then covered with its lid. The covered containers were placed under fluorescent lights at a distance of 20 cm from standard fluorescent cool white 4,100 K, 32-watt bulbs. The bulbs, emitting 2,800 lumens (Sylvania), were used to illuminate the plants for 5 days at ambient lab temperatures (22° C) on a 16/8 light/dark cycle. The plants were subsequently transferred to the greenhouse. The plants were removed from the container that allowed their recovery for 1 week. The eGFP fluorescence reporter was used to visually identify successful transgenic events using the Dark Reader Spot Lamp (Clare Chemical Research). Fluorescence did not occur in non-transformed roots. The effect the targeted *GmPGIP* gene had on its relative transcript abundance (RTA) was determined by real time quantitative PCR (RT-qPCR) The result, identified by eGFP reporter expression and confirmation by RT-qPCR was an initial stable transformation event in the root somatic cell that culminated in the generation of a transgenic root system, even though the DNA cassette was not incorporated into the germline. Notably, roots subsequently developed from the transgenic cell over a period of a few weeks through mitotic and developmental processes into a transgenic root system. The resultant genetically mosaic plants had a non-transgenic shoot with a transgenic root system. Consequently, the experiment produced individual transgenic root systems. Each root system was employed as an independent transformant (root system). RT-qPCR is described subsequently.

### Real time-Quantitative PCR (RT-qPCR)

The appropriate control genotype for the OE experiments was the *H. glycines* - susceptible genotype *G. max* _[Williams 82/PI 518671]_ unless otherwise noted. In contrast, the appropriate control genotype for the RNAi experiments was the *H. glycines* -resistant *G. max*_[Peking/PI548402]_ unless otherwise noted. The collected RNA samples served as template for cDNA generation (Invitrogen). RT-qPCR was used to examine *GmPGIP* gene expression. For these RT-qPCR-based studies, the *G. max* ribosomal protein gene *GmRPS21* (*Glyma.15G147700*) was used as the internal control. A Taqman 6 carboxyfluorescein-6-FAM (6-FAM) probe with Black Hole Quencher (BHQ1) was used for the RT-qPCR reactions that included the forward and reverse primers (MWG Operon) (**Supplemental Table 2**). The RT-qPCR reactions, performed on a StepOnePlus Real-Time PCR System (Applied Biosystems), had a pre-incubation of 50° C for 2 min, followed by 95° C for 10 min and alternating steps composing a cycle. There was a 95° C for 15 second (s) step followed by a 60° C for 1 min step with these two steps repeated in succession for 40 cycles. The 2^-ΔΔCt^ method was used to calculate the RTA (Livak and Schmittgen 2001). The statistical significance of RT-qPCR analysis results was determined by a Student’s t-test (Yuan et al. 2006).

### *H. glycines* infection of *G. max* and sample collection

*GmPGIP* function during the *G. max* defense response to *H. glycines* parasitism was studied through their OE and RNAi as compared to their appropriate controls. The *H. glycines*_[race 3/NL1-Rhg/HG-type 7]_ (*H. glycines)* was used for the infection of the transgenic roots to evaluate parasitism. *H. glycines*_[race 3/NL1-Rhg/HG-type 7]_ infection produced a compatible interaction that led to a susceptible reaction in *G. max*_[Williams 82/PI 518671]_. In contrast, *H. glycines* produced an incompatible interaction that led to a resistant reaction in *G. max*_[Peking/PI548402]_. *H. glycines* was used to infect the transgenic roots to evaluate the effect of the *GmPGIP* OE and RNAi constructs on parasitism as compared to their respective controls.

*H. glycines* eggs were collected from cysts extracted from 60-day-old greenhouse-grown stock infected *G. max* plants cultivated in 500-cm^3^ polystyrene pots. The sucrose flotation technique was used for *H. glycines* cyst extraction (Jenkins, 1964; Klink et al. 2021). The *H. glycines*-infected roots were washed in nested 850-µm-pore and 250-µm-pore sieves. The collection of cysts occurred from the 250-µm-pore sieve. The eggs were liberated from the cysts by grinding the cysts in a mortar and pestle. Gravitational sieving and subsequent sucrose centrifugation were used to collect the eggs. Eggs were recovered through nested 75-µm-pore over 25-µm-pore sieves. A modified Baermann funnel placed on a Slide Warmer (Model 77) (Marshall Scientific) at 28^1^ C was used to collect the J2s from hatched eggs. Egg hatching led to J2 liberation and occurred after 4–7 days. The J2 collection occurred on a 25-µm-pore sieve that was directed into 1.5-ml tubes. The centrifugation of the tube and its contents occurred at 10,000 rpm for 1 min, followed by content washing in sterile distilled water. A subsequent centrifugation occurred, concentrating the J2s in an IEC clinical centrifuge for 30 seconds (s) at 1,720 rpm. The final optimized concentration of J2s was 2,000 pre-infective (pi) J2/ml. Each targeted genetically chimeric, composite plant having the transgenic root, was inoculated with 1 ml (2,000 pi-J2) of inoculum. The J2s were placed in 1 cm holes made in the soil near the base of the root system. The holes were covered with soil once the inoculum had soaked into the soil to prevent expulsion of the J2s during watering. *H. glycines* infection was confirmed by the acid-fuchsin staining procedure of Byrd et al. (1983) on the 0, 3, and 6 dpi time point samples on the representative root system. The same procedure was used on the 9 dpi time point. Infection proceeded for 30 days. The *H. glycines* cysts were collected over nested 20- and 100-mesh sieves at the experiment’s conclusion. The soil was washed several times with the rinse water being sieved to ensure collection of all cysts for FI enumeration (Golden et al. 1970; Klink et al. 2021). Root samples were placed immediately in liquid nitrogen following their collection. The root samples were then stored in a -80°C freezer until processed for downstream analysis.

### Female index (FI) calculation and data analysis

The FI was calculated for each transgenic root system for each genetically mosaic composite plant that had a transgenic root and a non-transgenic shoot (Golden et al. 1970). The FI equation is FI = (Nx/Ns)*100 (Golden et al. 1970). Nx was the average number of females isolated from the pRAP15 or pRAP17 transgenic roots that contained the CRG under study (i.e. *GmPGIP*-OE or *GmPGIP*-RNAi) (Golden et al. 1970). Ns was the average number of females isolated from the respective, appropriate transgenic pRAP15-*ccd*B or pRAP17-*ccd*B empty vector control root (Golden et al. 1970). The FI was calculated as a function of the whole transgenic root system (wr) and also the mass (per gram [pg]) of the whole transgenic root system, tested statistically using the MWW Rank-Sum Test, p < 0.05. The wr analysis procedure is the historically-performed method used to enumerate cysts and does not account for any effect the plant genotype, transgenic event or even *H. glycines,* itself, has on parasitism in relation to root system growth. In contrast, the pg analysis procedure accounts for the effect the plant genotype, transgenic event, or *H. glycines* itself, has on parasitism in relation to root system growth (Pant et al. 2014). This relationship happens because the calculation of the FI adjusts for root system mass. Each transgenic root system from the genetically chimeric composite plant was considered an experimental replicate. The analyses all had three biological replicates, with at least 10 individual experimental replicates in each biological replicate, that were used to calculate the FI (Klink et al. 2021). The numbers of transgenic root system, spanning the 3 replicates, for each gene was *GmPGIP11*-OE: n = 36, pRAP15-*ccd*B: n = 36, *GmPGIP11*-RNAi: n = 36, pRAP17-*ccd*B: n = 36; *GmPGIP1*-OE: n = 36, pRAP15-*ccd*B: n = 36, *GmPGIP1*-RNAi: n = 36, pRAP17-*ccd*B: n = 36, consistent with other studies (Klink et al. 2021).

### Root system mass analysis

The effect that the altered *GmPGIP* expression had on root system mass was determined statistically from 3 biological replicates. Each biological replicate contained at least 10 individual experimental replicates. For the difference in root system mass, in each analysis the results were considered statistically significant if p < 0.05, determined using the MWW (Mann and Whitney, 1947).

### RNA-seq gene expression analyses of transgenic root RNA

Functional transgenic analyses of the 32-member *G. max MAPK* (*GmMAPK*) gene family demonstrated that *GmMAPK3-1* and *GmMAPK3-2* had a defense role during *H. glycines* parasitism (McNeece et al. 2019). The RNA sequencing (RNA seq) data under study was obtained from BioProject ID PRJNA664992, Submission ID: SUB8182387 (Alshehri et al. 2018). The template RNA used in the single replicate generation of RNA seq analyses was isolated from the respective *GmMAPK* overexpression (*GmMAPK3-1*-OE [*Glyma.U021800*] and *GmMAPK3-2*-OE [*Glyma.12G073000*]), *GmMAPK*-RNAi (*GmMAPK3-1*-RNAi and *GmMAPK3-2*-RNAi), respective OE (pRAP15-*ccd*B plasmid), and RNAi (pRAP17-*ccd*B plasmid) control expressing *G. max.* The RNA-seq results were confirmed by RT-qPCR of the *GmMAPK*s and numerous other targeted genes including 16 conserved oligomeric Golgi (COG) complex genes, 5 exocyst genes, and 8 putatively secreted genes (Sharma et al. 2020; Lawaju et al. 2020; Niraula et al. 2020). For the RNA-seq analyses, RNA was isolated from the collected root samples as already described. The collected samples were validated, sequenced, and analyzed, producing Illumina® RNA-seq data for use in examining gene expression of the 55,022 genes in the *G. max* genome (Alshehri et al. 2018). The RNA-seq fold change (FC) data, that represented the relative transcript abundance (RTA) was mined here specifically for *GmPGIP* gene paralog expression using the methods of Wang and Wang (2021). The FC for the OE experiments was determined in comparisons of the transgenic *GmMAPK-*OE RNA-seq data as compared to the RNA-seq data obtained from the transgenic pRAP15-*ccdB* (overexpression) control (Wang and Wang 2021). The FC for the RNAi experiments was determined in comparisons of the transgenic *GmMAPK-*RNAi RNA-seq data as compared to the RNA-seq data obtained from the transgenic pRAP17*-ccdB* (RNAi) control (Wang and Wang 2021). When presented, confirmation of the RNA-seq RTAs, given as FC, was performed by RT-qPCR using 2^-ΔΔCT^ as described (Livak and Schmittgen 2001; Klink et al. 2021).

#### Conglomerate gene expression analysis

Transcriptomic data for the conglomerate gene expression analysis for *G. max* PGIPs was obtained from the references cited here. *G. max* PGs were also studied due to their interactions with PGIPs. The *G. max* PGs of Wang et al. (2016) were employed. Gene expression pertaining to the whole root susceptible reaction was extracted (Vaghchhipawala et al. 2001; Khan et al. 2004; Tucker et al. 2007; Klink et al. 2007a; Ithal et al. 2007; Puthoff et al. 2007; Afzal et al. 2009; Tucker et al. 2011; Wan et al. 2015; Li et al. 2018; Song et al. 2019; Neupane et al. 2019b; Guo et al. 2020; Miraeiz et al. 2020; Shi et al. 2021; Kofsky et al. 2021; Torabi et al. 2023). Gene expression pertaining to the whole root resistant reaction was extracted (Alkharouf et al. 2004;(Klink et al. 2005, 2007b, 2009, 2010a, b, 2011; Ithal et al. 2007; Matsye et al. 2011); Wan et al. 2015; Li et al. 2018; Song et al. 2019; Neupane 2019; Miraeiz et al. 2020; Guo et al. 2020; Shi et al. 2021; Kofsky et al. 2021; Chu et al. 2022; Torabi et al. 2023). Analysis of the resistant syncytium include span 3, 6, 8, and 9 dpi (Klink et al. 2007b, 2009, 2010a, b, 2011; Matsye et al. 2011). Details of the analyses are available in those references. Analysis of the susceptible syncytium include span 3, 6, 8, and 9 dpi (Klink et al 2007, al. 2010b Matsye et al. 2011). The specific details regarding the materials and methods are presented in those referenced analyses.

## RESULTS

### The *G. max PGIP* (*GmPGIP*) gene family

A BLASTP search using the previously-identified *G. max* PGIP (*Gm*PGIP) Glyma.05G123700 led to the identification of 11 *Gm*PGIP-like sequences (**Table 1**) (Acharya et al. 2023). Two of these *Gm*PGIPs, Glyma.05G123800 (237 aa), and Glyma.08G078800 (142 aa) were much shorter in length than the other *Gm*PGIPs (334-373 aa). The much shorter length of these proteins indicated they could be the result of genetic truncations. However, genetically truncated *PGIP*s encoding proteins that are even one-third the size of the full length polypeptide can have the same activity against some pathogen PGs (Chiu et al. 2021). Thus, the shorter *GmPGIP*s were included in subsequent bioinformatics analyses.

**Table 1.** *G. max PGIP* genes.

| Protein | Accession 1 | Length | (aa) | Signal | N | O | Subcellular |
| --- | --- | --- | --- | --- | --- | --- | --- |
| *GmPGIP1* | Glyma.05G123700.1 |  | 336 | y | y | y | extracellular |
| *GmPGIP2* | Glyma.05G123800.1 |  | 237 | n | n | y | cytoplasm |
| *GmPGIP3* | Glyma.05G123900.1 |  | 337 | y | y | y | extracellular |
| *GmPGIP4* | Glyma.05G124000.1 |  | 335 | y | y | y | extracellular |
| *GmPGIP5* | Glyma.08G078800.1 |  | 142 | n | n | y | nucleus |
| *GmPGIP6* | Glyma.08G078900.1 |  | 337 | y | y | y | extracellular |
| *GmPGIP7* | Glyma.08G079100.1 |  | 333 | y | y | y | extracellular |
| *GmPGIP8* | Glyma.08G079200.1 |  | 334 | y | y | y | extracellular |
| *GmPGIP9* | Glyma.15G209200.1 |  | 373 | y | y | y | cell |
| membrane |  |  |  |  |  |  |  |
| *GmPGIP10* | Glyma.15G209300.1 |  | 340 | n | y | y | extracellular |
| *GmPGIP11* | Glyma.19G145200.1 |  | 351 | y | y | n | extracellular |

Genome analyses were performed to better understand the chromosomal regions where the identified, conceptually translated *GmPGIP* sequences exist as duplicated genes (Kalunke et al. 2014). The region of *G. max*_[Williams 82/PI 518671]_ chromosome 5 was confirmed here to have 4 tandemly duplicated *GmPGIP* copies, that included *Glyma.05G123700*, *Glyma.05G123800*, *Glyma.05G123900*, and *Glyma.05G124000*. The region of chromosome 8 was confirmed to have 4 *GmPGIP* copies that included *Glyma.08G078800*, *Glyma.08G078900*, *Glyma.08G079100*, and *Glyma.08G079200*. The arrangement of the genes had begun with the tandem copies of the *Glyma.08G078800*, *Glyma.08G078900*, followed by an annotated N-acetyl-gamma-glutamyl-phosphate reductase/NAGSA dehydrogenase (*Glyma.08G079000*) that was then followed by *Glyma.08G079100* and *Glyma.08G079200*. The region of chromosome 15 was confirmed to have 2 tandemly duplicated copies of *PGIP* that included *Glyma.15G209200*, and *Glyma.15G209300*. The region of *G. max*_[Williams 82/PI 518671]_ chromosome 19 was confirmed to have the *GmPGIP Glyma.19G145200*. An examination of other *G. max* genotypes was beyond the scope of the analysis. Comparative analyses of identified *Gm*PGIP proteins show the *G. max* accession numbers for the previously identified Gmpgip, naming convention by D’Ovidio et al. (2006) Gmpgip1 AJ972660, was Glyma.05G124000, Gmpgip2 AJ972661 was Glyma.05G123900, Gmpgip3 AJ972662 was Glyma.08G079100.1, and Gmpgip4, AJ972663 was Glyma.08G079200.1 (D’Ovidio et al. 2006; Kalunke et al. 2014). The remaining Gmpgip5 and Gmpgip7 protein accessions could not be determined since their sequences were not available (D’Ovidio et al. 2006; Kalunke et al. 2014). The *Gm*PGIP protein sequence (AF130974.1 [AAD45503]) found in an *H. glycines* -resistant *G. max* _[PI 437654]_ was 100% identical to Glyma.08G079100 but was missing the N-terminal 19 aa sequence MSKLSILFLLVLSFSSVLS (Mahalingam et al 1999). The next most closely related *Gm*PGIP sequence to AF130974.1 was Glyma.05G123700.1, having 72% identity so AF130974.1 was most likely Glyma.08G079100. BLASTP analyses demonstrated the Gmpgip1 (AJ972660) was most likely Glyma.05G124000. The Gmpgip2 (AJ972661) was most likely Glyma.05G123900. The Gmpgip3 (AJ972662) was most likely Glyma.08G079100. The Gmpgip4 (AJ972663) was most likely Glyma.08G079200. This conclusion was made because each PGIP had 100% identity along their full length protein sequences with any other related sequence having much lower identity.

### *G. max* PGIPs have features consistent with being secreted proteins

PGIPs act outside of the cell, requiring a functional secretion system and processing for activity (Powell et al. 2000; Prabhu et al. 2015). The 11 *Gm*PGIPs were examined for the presence of a signal peptide and accompanying signal peptidase cleavage site. The results of those analyses using SignalP 6.0 demonstrated *Gm*PGIP1, *Gm*PGIP3, *Gm*PGIP4, *Gm*PGIP6, *Gm*PGIP7, *Gm*PGIP8, *Gm*PGIP9, and *Gm*PGIP11 were predicted to have signal peptides, and a predicted cleavage site (**Figure 1**, **Table 1)**. This result was mostly consistent with those obtained by Deeploc 1.0 which obtained extracellular locations for *Gm*PGIP1, *Gm*PGIP3, *Gm*PGIP4, *Gm*PGIP6, *Gm*PGIP7, *Gm*PGIP8, *Gm*PGIP10, *Gm*PGIP11 (long [L] and short [S] predictions). Thus, there was a discrepancy between Deeploc 1.0 which predicted a signal peptide for *Gm*PGIP10 and SignalP 6.0 which did not. *Gm*PGIP11 had a long signal sequence that was not supported as strongly through Signal P 6.0 analyses (**Figure 1**, **Table 1)**. A BLASTP search using the 351 aa *G. max Gm*PGIP11 sequence led to the identification of a predicted 333 aa *Gm*PGIP11 homolog (KAG4913035) from cultivar Huaxia 3 (*G. max* _[Huaxia 3]_) that matched *Gm*PGIP11 100% over the 333 aa sequence and had a better supported signal peptide (likelihood *Gm*PGIP11^[KAG4913035]^ = 0.9997 vs. likelihood *Gm*PGIP1^[Glyma.19G145200/GmPGIP11]^ = 0.5249) (**Supplemental Figure 1)** (Chiu et al. 2021). Furthermore, SignalP 6.0 calculated a cleavage site that occurred between aa positions 22 and 23 with a probability of 0.977916 for *Gm*PGIP11^[KAG4913035]^ while a predicted cleavage site occurred between aa positions 40 and 41 with a probability of 0.502115 for *Gm*PGIP1^[Glyma.19G145200/GmPGIP11]^. *Gm*PGIP10 was not predicted to have a signal sequence (**Figure 1**, **Table 1)**. The shorter *Gm*PGIP2, and *Gm*PGIP5 lack predicted signal peptides (**Supplemental Figure 2**, **Table 1**).

**Figure 1.**
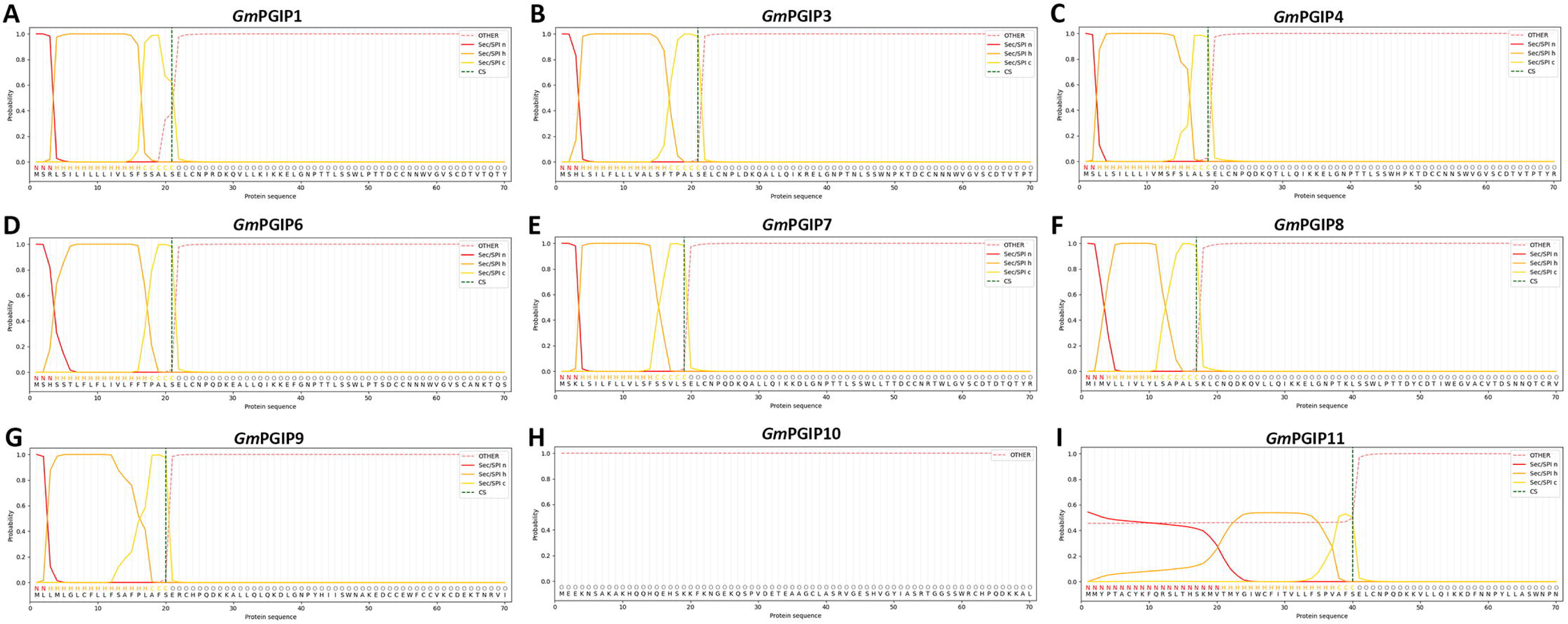
PGIP signal peptide prediction. **A**. *Gm*PGIP1. **B**. *Gm*PGIP3. **C**. *Gm*PGIP4. **D**. *Gm*PGIP6. **E**. *Gm*PGIP7. **F**. *Gm*PGIP8. **G**. *Gm*PGIP9. **H**. *Gm*PGIP10. **I**. *Gm*PGIP11. Signal peptides were not predicted for *Gm*PGIP2, *Gm*PGIP5 under the analysis procedures. Signal peptide-the N-terminal n-region (Sec/SPI n); signal peptide-the N-terminal hydrophobic-region (Sec/SPI h); signal peptide-the C-terminal c-region (Sec/SPI c); cleavage sites, CS; other, O. The region of the protein that is relevant to signal peptide prediction is shown (Teufel et al. 2022).

The Golgi apparatus acts in protein glycosylation, important to the function of secreted proteins (Kingsley et al. 1986). The results from the *N*-linked glycosylation analysis are shown (**Table 1, Supplemental Figure 3)**. The results from the *O*-linked glycosylation analysis are shown (**Table 1**, **Supplemental Figure 4**). Prediction of the cellular localization for the predicted PGIP proteins was done using the Deeplock 1.0 artificial intelligence software. In this analysis, firstly, the importance of each aa of each *Gm*PGIP protein, including two potential forms of *Gm*PGIP11 (the 351 aa PGIP11-L, and 333 aa PGIP11-S) were done (Almagro Armenteros et al. 2017). In the first analysis, the importance of a particular aa along the *Gm*PGIP protein chain that was relevant for prediction (attention) of its subcellular location was determined under default settings (**Table 1, Supplemental Figure 5)** (Almagro Armenteros et al. 2017). The removal of the 18 aa sequence that was found in the predicted *Gm*PGIP11-L changed its attention values at the N-terminus of the protein (**Table 1, Supplemental Figure 5)**. Subsequently, the subcellular localization was predicted (**Table 1, Supplemental Figure 6)**. Furthermore, the *Gm*PGIP11 with the N-terminal 18 aa removed was characterized for its protein localization, improving the probability from the longer, 351 aa predicted protein (p = 0.0914) to the 333 aa shorter version (p = 0.994) that it is secreted as expected (**Supplemental Figure 6**).

### *GmPGIP*s are expressed in *H. glycines*-parasitized root cells undergoing a defense response

Gene expression analyses were done using RNA isolated from *G. max*_[Peking/PI 548402]_ and *G. max*_[PI 88788]_ control cells at 0 dpi, and syncytia that underwent a defense response at 3, and 6 dpi to *H. glycines*_[race 3/NL1-Rhg/HG-type 7]_ to generate probe for microarray analyses (**Figure 2)**. An additional 9 dpi time point, performed at the same time as the 0, 3, and 6 dpi time point studies, was also analyzed with the results presented here. The analyses identified *GmPGIP11* as being expressed within the syncytia of *G. max*_[Peking/PI 548402]_ and *G. max*_[PI 88788]_, each of which was undergoing a defense response at 3, and 6 dpi as compared to the control which lacked measurable expression (**Table 2; Supplemental Table 3)**. The remaining *GmPGIP*s lack measurable expression in the syncytium and control cells, which indicated they may not perform a role in the defense response (**Table 2**; **Supplemental Table 3)**. The 9 dpi time point revealed that only *G. max* _[Peking/PI 548402]_ has measurable expression in each of the 3 replicates while *G. max* _[PI 88788]_ lacked expression in any one of the replicates (**Table 2; Supplemental Table 3)**. The results indicated *GmPGIP11* was a reasonable defense gene candidate that exhibited features of secreted proteins, to test to determine if its overexpression impairs *H. glycines* parasitism and whether RNAi facilitates parasitism (Pant et al. 2014; Niraula et al. 2020).

**Figure 2.**
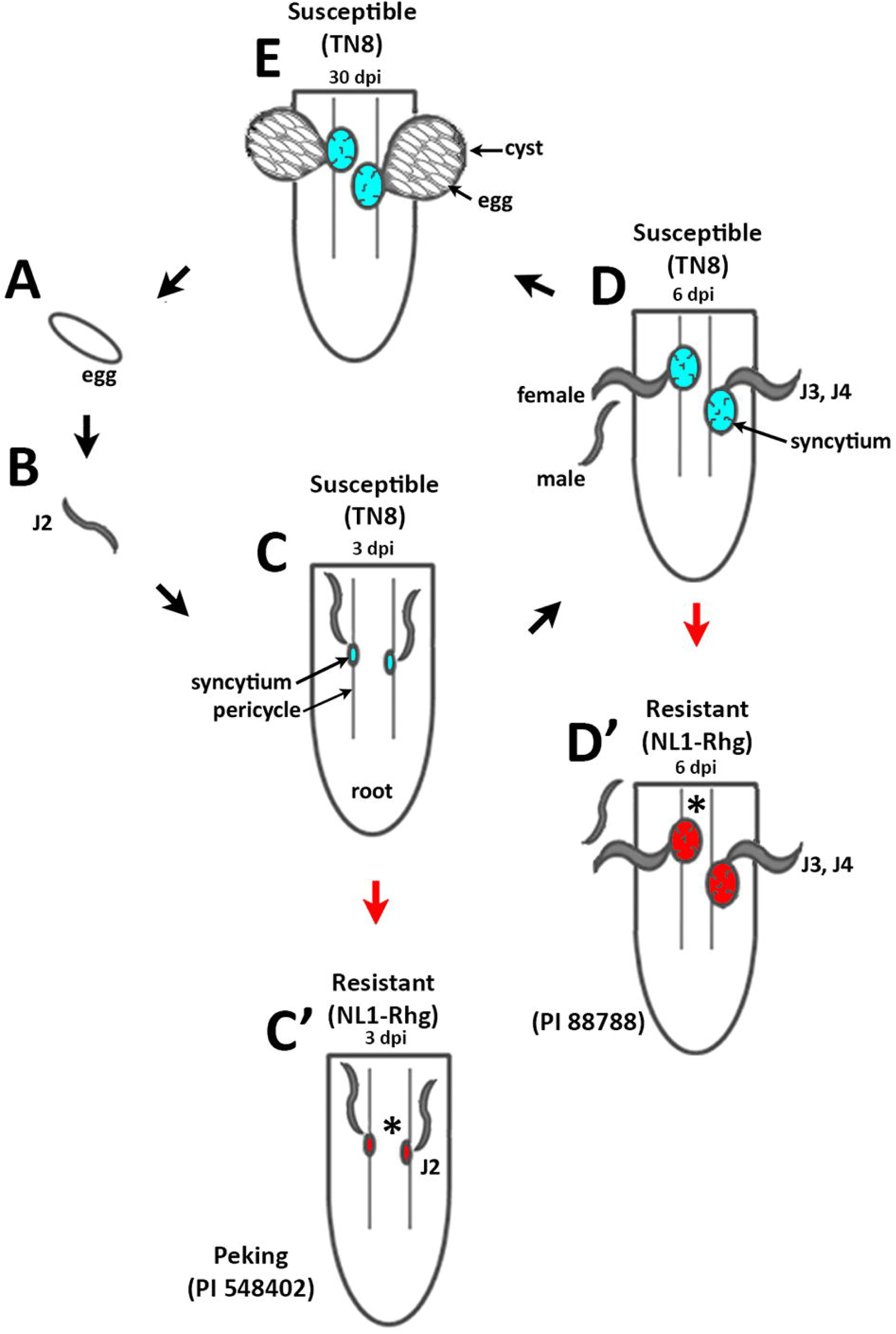
Life cycle of *H. glycines* in *G. max*. Egg; juvenile (J) stages, second (J2), third (J3), and fourth (J4). Sexually mature stages; Male, and female. The syncytium formed through the merging of cytoplasm of *H. glycines*-parasitized pericycle cells, and the cytoplasm of 200 to 250 neighboring cells. The cyst is the dead female carcass. **A**. Eggs. **B**. J2s infect *G. max*_[Peking/PI 548402]_ and *G. max* _[PI 88788]_. **C**. During the susceptible reaction in *G. max* _[Peking/PI 548402]_, *H. glycines* _[race 14/HG-type 1.3.6.7/TN8]_ continue to develop through the J2 stage as it produces a syncytium (cyan). **C’**. in *G. max* _[Peking/PI 548402]_, the defense response (red arrow) arrests *H. glycines* _[race 3/HG-type 7/NL1-Rhg]_ at the J2 stage producing a smaller syncytium (red). **D**. Susceptible reactions (syncytium colored in cyan) are obtained with *H. glycines* _[race 14/HG-type 1.3.6.7/TN8]_. **D’**. Infection with *H. glycines*_[race 3/HG-type 7/NL1-Rhg]_ results in a defense response in both *G. max* _[Peking/PI 548402]_ and *G. max*_[PI 88788]_. The defense response of *G. max* _[PI 88788]_ (red arrow) arrests *H. glycines*_[race 3/HG-type 7/NL1-Rhg]_ development at the J3–J4 stages, resulting in a larger syncytium that becomes non-functional. **E**. The *H. glycines* _[race 14/HG-type 1.3.6.7/TN8]_ life cycle during a susceptible reaction is completed by approximately 30 dpi in both *G. max*_[Peking/PI 548402]_ and *G. max*_[PI 88788]_ (Endo 1964, 1965, 1991; Riggs et al. 1973; Acido et al. 1984; Kim et al. 1987; Kim and Riggs 1992; Schmelzer 2002; Hardham et al. 2008; Colgrove and Niblack 2008).

**Table 2.** *GmPGIP* gene expression occurring during the defense response to *H. glycines* parasitism. *Footnote: The 9 dpi time point revealed that only G. max_[Peking/PI 548402]_ has measurable expression in each of the 3 replicates while G. max_[PI 88788]_ lacked expression in any one of the replicates.

| Gene | Accession | Affymetrix probe ID | Time point |  |  |  |
| --- | --- | --- | --- | --- | --- | --- |
|  |  |  | 0 | 3 | 6 | 9 |
| *GmPGIP1* | Glyma.05G123700.1 | GmaAffx.68517.2.S1_at | NM | NM | NM | NM |
| *GmPGIP2* | Glyma.05G123800.1 | no probe | n/a | n/a | n/a | n/a |
| *GmPGIP3* | Glyma.05G123900.1 | Gma.4142.1.S1_at | NM | NM | NM | NM |
| *GmPGIP4* | Glyma.05G124000.1 | GmaAffx.93403.1.A1_s_at | NM | NM | NM | NM |
| *GmPGIP5* | Glyma.08G078800.1 | no probe | n/a | n/a | n/a | n/a |
| *GmPGIP6* | Glyma.08G078900.1 | GmaAffx.18518.1.S1_at | NM | NM | NM | NM |
| *GmPGIP7* | Glyma.08G079100.1 | no probe | n/a | n/a | n/a | n/a |
| *GmPGIP8* | Glyma.08G079200.1 | no probe | n/a | n/a | n/a | n/a |
| *GmPGIP9* | Glyma.15G209200.1 | no probe | n/a | n/a | n/a | n/a |
| *GmPGIP10* | Glyma.15G209300.1 | GmaAffx.91749.1.S1_at | NM | NM | NM | NM |
| *GmPGIP11* | Glyma.19G145200.1 | GmaAffx.47891.1.S1_s_at | NM | M | M | N/M* |

### The production of transgenic *GmPGIP11* genetically mosaic composite plants

The *GmPGIP11* gene was targeted for genetically manipulated expression through transgenesis, altering its RTA as compared to the appropriate control. The *GmPGIP11* gene was engineered for OE in the *H. glycines* -susceptible *G. max* _[Williams 82/PI 518671]_. This experimental approach was based on the hypothesis that an increase in the targeted *GmPGIP11* RTA would change the *H. glycines*-susceptible *G. max* _[Williams 82/PI 518671]_ so it produced a defense response outcome resembling the *H. glycines*-resistant *G. max* _[Peking/PI 548402]_. In contrast, the *GmPGIP11* gene engineered for RNAi in the *H. glycines* -resistant *G. max* _[Peking/PI 548402]_ would decrease resistance. This approach was based on the hypothesis that decreasing the *GmPGIP11* RTA would change the *H. glycines* -resistant *G. max* _[Peking/PI 548402]_ so it resembled the observed impaired defense response (i.e., susceptibility) occurring in *G. max* _[Williams 82/PI 518671]_. Importantly, the combined hypothesized increase in *GmPGIP11* RTA in *G. max*_[Williams 82/PI 518671]_ that led to root systems that were more *H. glycines*-resistant and decrease in *GmPGIP11* RTA in the *G. max* _[Peking/PI 548402]_ that led to root systems that were more *H. glycines* -susceptible was accepted as evidence that the targeted *GmPGIP11* gene functioned in the defense response (Matsye et al. 2012; Pant et al. 2014). *GmPGIP11* OE and RNAi transformant roots, and their respective transgenic pRAP15-*ccd*B OE, and pRAP17-*ccd*B RNAi control roots, were made (**Figure 3)**. The RTAs of *GmPGIP11* in OE and RNAi roots are presented as compared to their respective transgenic controls (**Figure 3)**. The control gene employed in the RT-qPCR analysis was *GmRPS21* (**Figure 3)**. *GmPGIP11* RTAs in the OE roots increased 9.7-fold as compared to their control (p < 0.05, Student’s t-test) (**Figure 3**). In contrast, the *GmPGIP11* RTA decreased - 8.4-fold in the RNAi roots as compared to their control (p < 0.05, Student’s t-test) (**Figure 3**).

**Figure 3.**
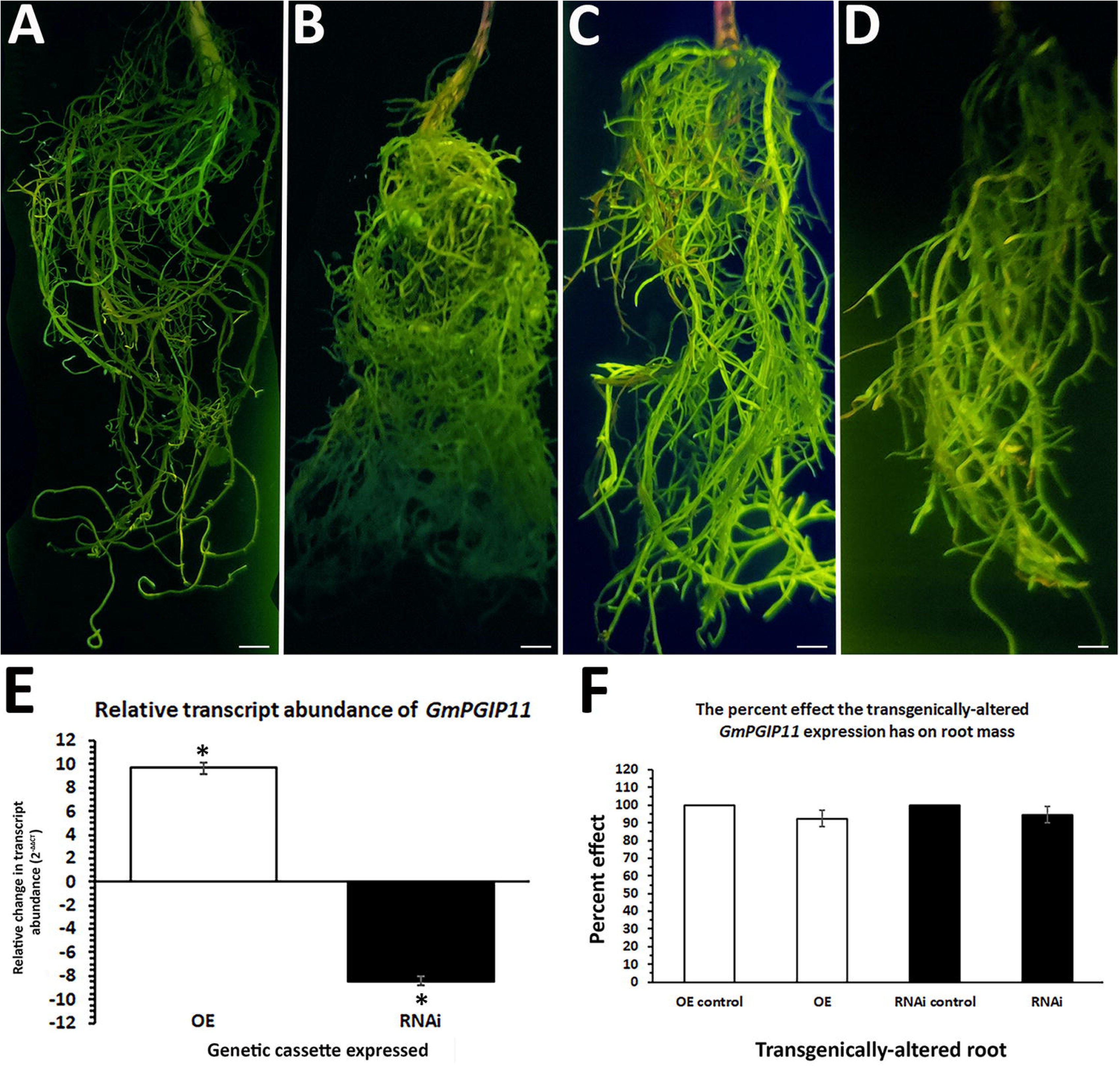
Generation of transgenic roots Terminology: overexpression (OE); RNA interference (RNAi).**A**. pRAP15-*ccd*B (OE control); **B**. *GmPGIP11-*OE; **C**. pRAP17-*ccd*B (RNAi control); **D**. *GmPGIP11-*RNAi; Bars = 1 cm. **E.** The change in *GmPGIP11* RTA, presented as FC, for the -OE and -RNAi roots, as compared to their pRAP15-*ccd*B and pRAP17-*ccd*B overexpression and RNAi controls, respectively, was calculated by 2^-ΔΔCT^ to determine FC (Livak and Schmittgen 2001). The RTA of the candidate defense genes, presented as a FC, in the transgenic roots was compared *GmRPS21.* **F**. The effect that the expression of the *GmPGIP11*-OE or -RNAi cassettes had on root mass as compared to their respective pRAP15-*ccd*B or pRAP17-*ccd*B controls. Statistical significance (*) for root mass difference was determined using the MWW Rank-Sum Test, p < 0.05 (Mann and Whitney, 1947).

### *GmPGIP11* expression does not alter root growth

The overexpression and RNAi of genes can impact root development. Consequently, an analysis was performed to address this issue. No statistically significant effect was observed for root system mass for the *GmPGIP11*-OE or *GmPGIP11*-RNAi root systems as compared to their respective controls (p ≥ 0.05, MWW) (**Figure 3**).

#### Altered *GmPGIP11* expression affects *H. glycines* parasitism in *G. max*

The experimental procedure for testing the effect of *GmPGIP11* overexpression is presented (**Figure 4)**. Soil in which transgenic *GmPGIP11*-OE roots of genetically mosaic, composite plants were growing were infested with 2,000 J2 *H. glycines*. *H. glycines* infection of *G. max* occurred, with subsequent parasitism proceeding for 30 days. The OE-controls were treated in the same manner. The experimental procedure for testing the effect of *GmPGIP11*-OE is presented (**Figure 4**). The results of the FI analysis for cysts per whole root (wr) system, also standardized to cysts per gram (pg) of root system for the *GmPGIP11*-OE transgenic roots are presented (**Figure 4**). The experimental procedure for testing the effect of *GmPGIP11*-RNAi is presented (**Figure 4**). The results of the FI analysis for the wr system, also standardized in the pg analyses for the *GmPGIP11*-RNAi transgenic roots are presented (**Figure 4)**.The FI analyses, showing a combined decrease in *H. glycines* cysts the *GmPGIP11*-OE experiment compared to its control and an increase in in *H. glycines* cysts the *GmPGIP11*-RNAi experiment, compared to its control, demonstrated that *GmPGIP11* performs a defense role toward *H. glycines* parasitism in the root.

**Figure 4.**
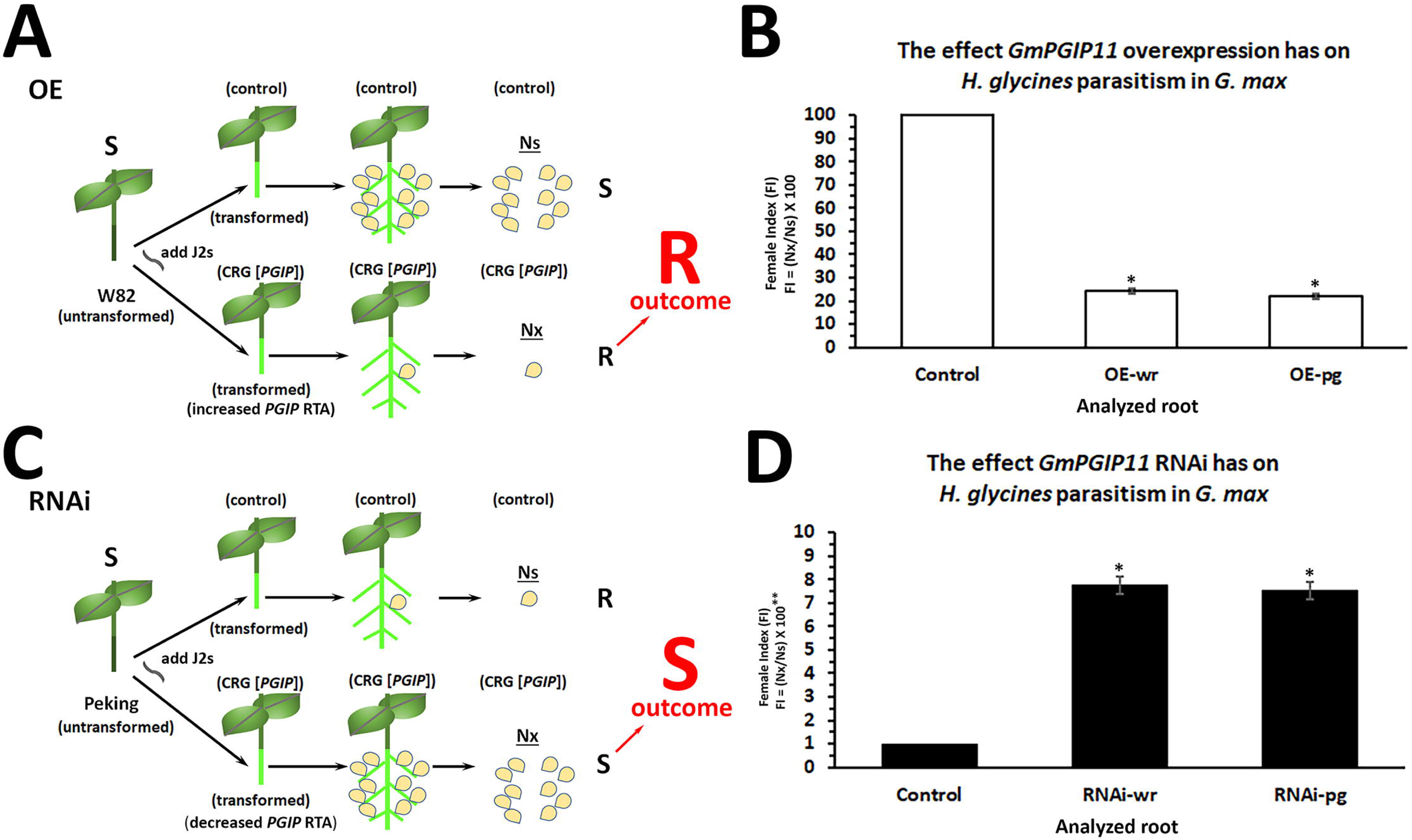
Experimental pipeline showing the calculation of the FI for *GmPGIP11* OE and RNAi roots as compared to their respective pRAP15-*ccd*B and pRAP17-*ccd*B controls. **A**. In the overexpression (OE) model, the susceptible (S), untransformed, non-transgenic *G. max* _[Williams 82/PI 518671]_ (W82) plant (shown with 2 leaves) had its root (dark gray) removed at the hypocotyl in the presence of K599 transformed either with the *GmPGIP11*-OE experimental CRG construct (below) or the pRAP15-*ccd*B control (above). After the generation of transgenic roots (green), evident by the visual green fluorescent protein (eGFP) reporter and confirmation by RT-qPCR of the e*GFP* and CRG, the soil of potted, genetically mosaic composite plants was infested with 2,000 J2 *H. glycines* with subsequent root infection and parasitism. *H. glycines* may develop into cysts (yellow oval). Under control conditions (Ns), the susceptible *G. max* _[Williams 82/PI 518671]_ permitted the unhindered development of *H. glycines* as evident by cysts (above). In contrast, the experimental overexpression of the *GmPGIP* gene condition (Ns), if it performs a defense role as hypothesized by its expression during the defense response, hindered cyst development (below). To determine whether the CRG overexpression performed a defense role that led to resistance (R) the FI was calculated by the equation Nx/Ns X 100 (Golden et al. 1970). **B**. *GmPGIP11*-OE FI results as compared to the pRAP15-*ccd*B OE control. The control, compared to itself, had a FI of 100. The FI analysis was done in 2 different manners. The first manner analyzed the number of *H. glycines* cysts per whole root system (wr) which did not account for any effect the expression of the transgene, *H. glycines* or combination of the transgene and *H. glycines* had on root development. The second manner took into consideration the development of the root by measuring the number of cysts per gram of root system (pg), thus standardizing the data. **C**.. In the RNAi model, the resistant (R), untransformed, non-transgenic G. max_[PI 548402/Peking]_ (Peking) plant (shown with 2 leaves) had its root (dark gray) removed at the hypocotyl in the presence of K599 transformed either with the *GmPGIP11*-RNAi experimental CRG construct (below) or the pRAP17-*ccd*B control (above). After the generation of transgenic roots (green), evident by the visual green fluorescent protein (eGFP) reporter and confirmation by RT-qPCR of the e*GFP* and *GmPGIP11* CRG, the soil of potted, genetically mosaic composite plants was infested with 2,000 J2 *H. glycines* with subsequent root infection and parasitism. *H. glycines* may develop into cysts (yellow oval). Under control conditions (Ns), the resistant *G. max*_[Williams 82/PI 518671]_ permitted only hindered development of *H. glycines* as evident by comparatively fewer cysts (above). In contrast, the experimental RNAi of the *GmPGIP11* CRG condition (Ns), if it performs a defense role as hypothesized by its expression during the defense response, permitted a comparatively less hindered cyst development (below). To determine whether the CRG RNAi impaired a defense role that led to susceptibility (S) the FI was calculated by the equation Nx/Ns X 100 (Golden et al. 1970). **D**. *GmPGIP11*-RNAi FI results as compared to the pRAP17-*ccd*B RNAi control (Golden et al. 1970). As in the OE experiment, the control, compared to itself, had a FI of 100. As explained for the CRG-OE experiment, the FI analysis analyzed the number of *H. glycines* cysts per whole root system (wr) and the number of cysts per gram of root system (pg), which standardized the data. The statistical significance (*) was determined by the MWW Rank-Sum Test, p < 0.05 (Mann and Whitney, 1947). The combination of a statistically significant decrease in FI in the CRG-OE and a statistically significant increase in FI in the CRG-RNAi experiment was taken as evidence that the gene performed a defense response that led to a resistant reaction (Pant et al. 2014). For details, please refer to the Results section, subsections: Genetic engineering of *G. max,* Real time-Quantitative PCR (RT-qPCR), *H. glycines* infection of *G. max* and sample collection, Female index (FI) calculation and data analysis.

### *GmPGIP11* RTA is regulated by components of PTI and ETI

Analyses demonstrated PTI and ETI components function in the defense response that *G. max* has to *H. glycines* (Pant et al. 2014; Aljaafri et al. 2017; McNeece et al. 2017; Klink et al. 2021; Niraula et al. 2022b). The PTI components included *GmBAK1-1* and *GmBIK1-6* (Pant et al. 2014; Klink et al. 2021). The ETI components included *GmNDR1-1* and *GmRIN4-4* (Aljaafri et al. 2017; McNeece et al. 2017; Niraula et al. 2022b). Experiments presented here demonstrated the overexpression of the PTI components *GmBAK1-1* and *GmBIK1-6* increased *GmPGIP11* RTA as compared to its pRAP15-*ccd*B OE control while their RNAi decreases it as compared to the pRAP17-*ccd*B RNAi control (**Figure 5**). Experiments presented here demonstrated the overexpression of the ETI components *GmNDR1-1* and *GmRIN4-4* increase *GmPGIP11* RTA as compared to its pRAP15-*ccd*B OE control while their RNAi decreased their RTAs as compared to the pRAP17-*ccd*B control (**Figure 5**).

**Figure 5.**
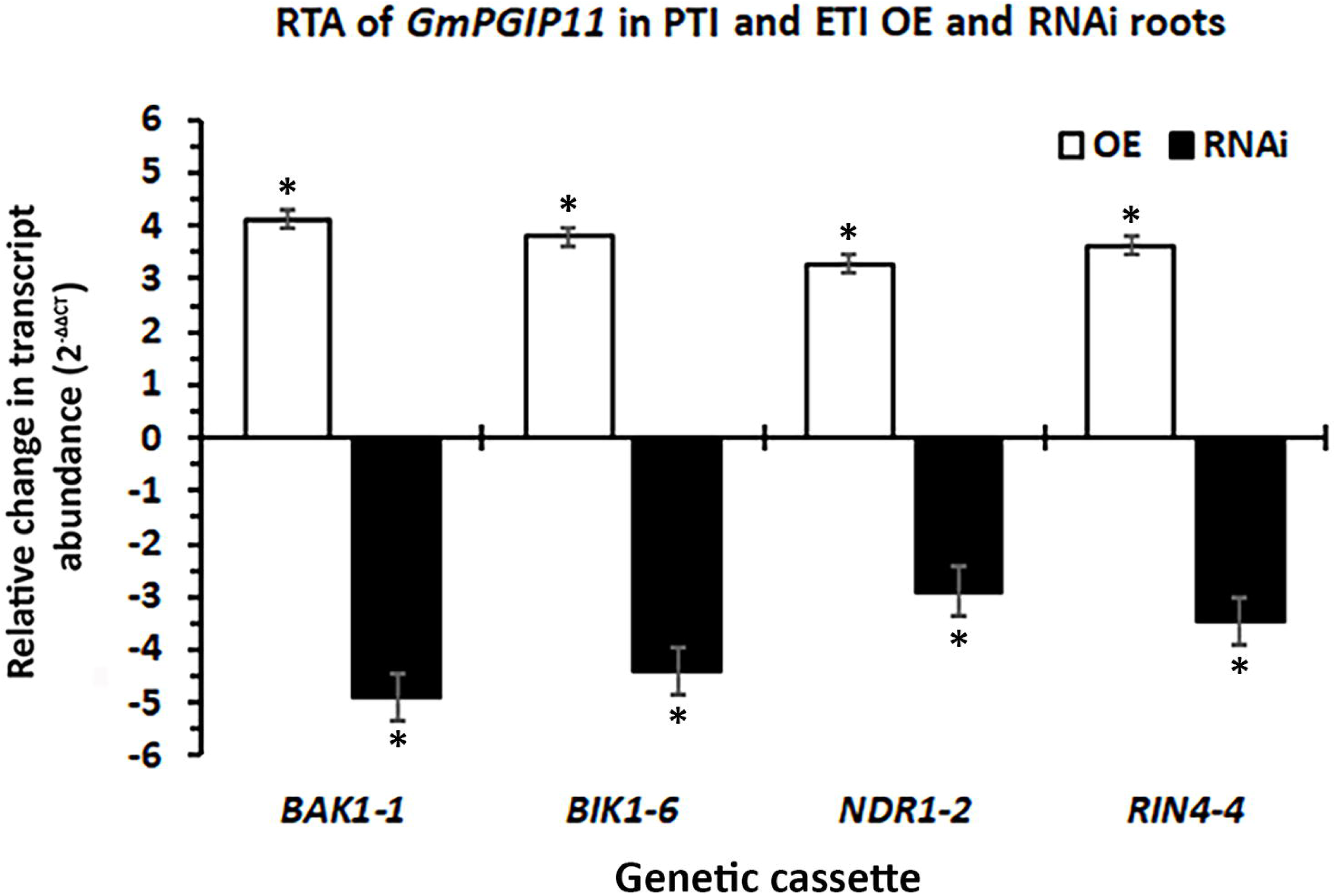
The altered expression of PTI and ETI genes influence *GmPGIP11* RTA. *GmPGIP11* RTA in the root RNA for the PTI genes *GmBAK1-1* and *GmBIK1-6* (OE [white histogram]and RNAi [black histogram]) as compared to the pRAP15-*ccd*B OE and pRAP17-*ccd*B RNAi controls, respectively. *GmPGIP* RTA in the root RNA of the ETI genes *GmNDR1-1* and *GmRIN4-4* (OE [white histogram] and RNAi [black histogram]) as compared to the pRAP15-*ccd*B OE and pRAP17-*ccd*B RNAi controls, respectively, calculated by 2^-ΔΔCT^ (Livak and Schmittgen 2001). The RT-qPCR analyses examined 3 experimental replicates (individual root systems) for *GmBAK1-1*, *GmBIK1-6*, *GmNDR1-1*, and *GmRIN4-4* -OE or -RNAi roots as compared to their pRAP15-*ccd*B, or pRAP17-*ccd*B controls, respectively, from each of the 3 biological replicates. Each experimental replicate is run in triplicate using the same RNA. (*), statistical significance of p < 0.05, Student’s *t*-test (Yuan et al. 2006).

### *GmPGIP* expression is regulated by *GmMAPK*s

McNeece et al. (2019) showed *GmMAPK3-1* and *GmMAPK3-2* have a defense role toward *H. glycines* parasitism. An in-depth analysis of RNA seq data for *GmMAPK3-1*-OE, *GmMAPK3-1*-RNAi, *GmMAPK3-2*-OE, and *GmMAPK3-2*-RNAi is presented (**Figure 6**). The RNA seq data was confirmed by RT-qPCR (**Figure 6**). *GmPGIP* genes not exhibiting expression by RNA seq analyses were not examined further.

**Figure 6.**
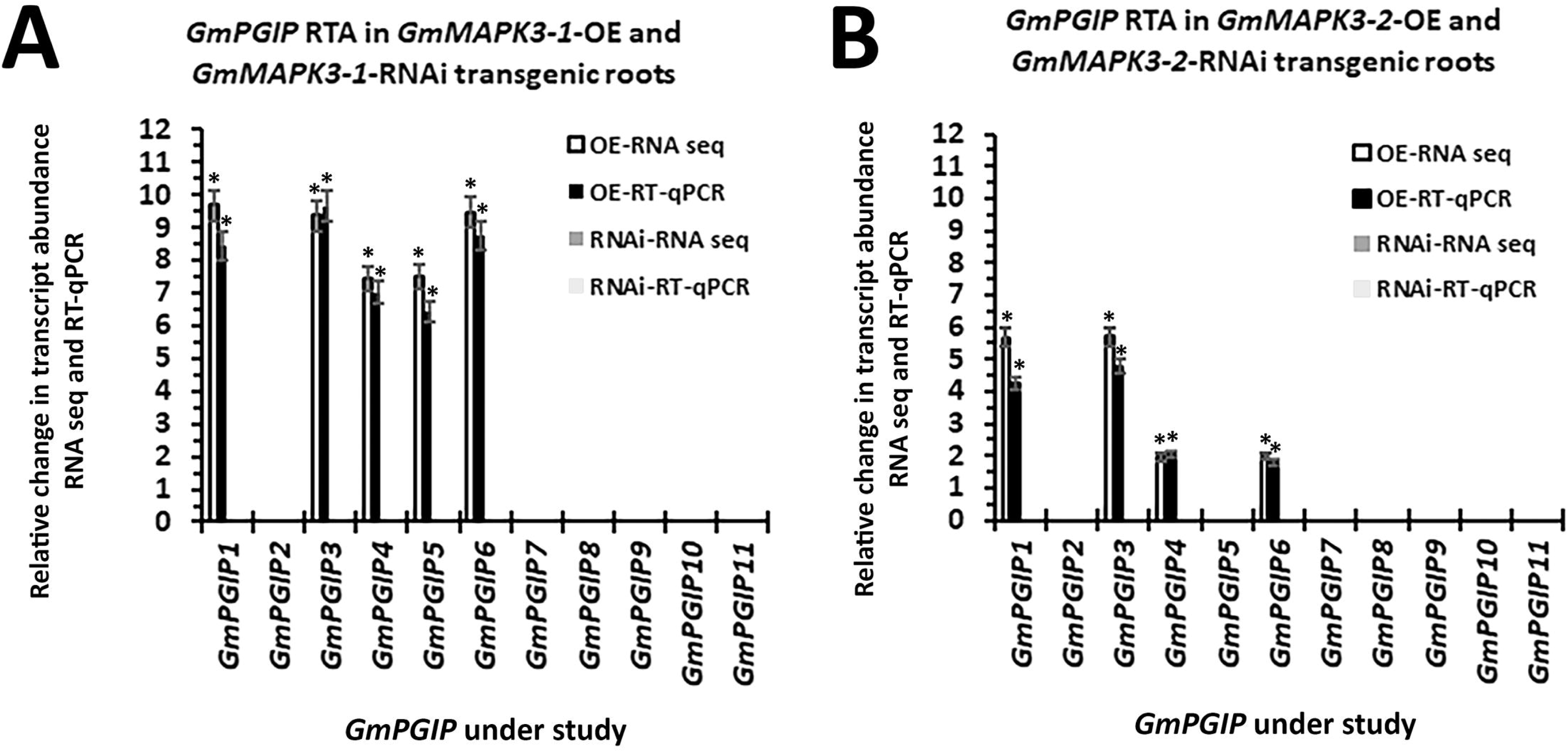
Analysis of RNA seq data for *GmMAPK3-1*-OE, *GmMAPK3-1*-RNAi, *GmMAPK3-2*-OE, and *GmMAPK3-2*-RNAi derived from BioProject ID PRJNA664992. *GmMAPK3-1*-OE and *GmMAPK3-2*-OE comparisons were made to its pRAP15-*ccd*B control. *GmMAPK3-1*-RNAi and *GmMAPK3-2*-RNAi comparisons were made to its pRAP17 control. The RNA seq resultswere confirmed by RT-qPCR. Relative change in transcript abundance of RNA seq data was calculated according to Wang and Wang (2021). The RT-qPCR transcript abundance was calculated by the 2^-ΔΔCt^ method (Livak and Schmittgen, 2001). The statistical significance of RT-qPCR analysis results was determined by a Student’s t-test, statistical significance of p < 0.05 (Yuan et al., 2006).

### The regulation of *GmPGIP* expression by MAPKs does not presage a defense role

The regulation of *GmPGIP1*, *GmPGIP3*, *GmPGIP4*, and *GmPGIP6* by both *GmMAPK3-1* and *GmMAPK3-2* OE, and RNAi contrasted with the LM expression data obtained from the syncytia of *G. max* _[Peking/PI 548402]_ and *G. max* _[PI 88788]_ genotypes which showed they lack expression (**Table 2**, **Supplemental Table 3**). *PGIP*s function in defense toward different types of pathogens, and their regulation may be influenced by *MAPK*s. Therefore, the MAPK regulation of *PGIP* expression may not necessarily relate to a defense role to *H. glycines*, especially if the genes lack expression in and around the cells where the defense response was expected to be happening (i.e., the syncytium). To test this hypothesis, *GmPGIP1* (*Glyma.05G123700*) which lacked expression in any LM-examined replicate and exhibited altered RTA in *GmMAPK3-1* and *GmMAPK3-2* OE roots, were cloned for their OE and RNAi. Experiments showed the expected increased *GmPGIP1* RTA in its OE root RNA as compared to its pRAP15-*ccd*B OE controls and the expected decreased *GmPGIP1* RTA in its RNAi root RNA as compared to its pRAP17-*ccd*B OE controls, respectively (**Figure 7**). The *GmPGIP1*-OE and *GmPGIP1*-RNAi FI, the relative measure of cysts, did not change as compared to their respective controls (**Figure 7**). The OE-of PTI and ETI genes led to increased *GmPGIP1* RTA, while their RNAi decreases *GmPGIP1* RTA, respectively (**Figure 7**).

**Figure 7.**
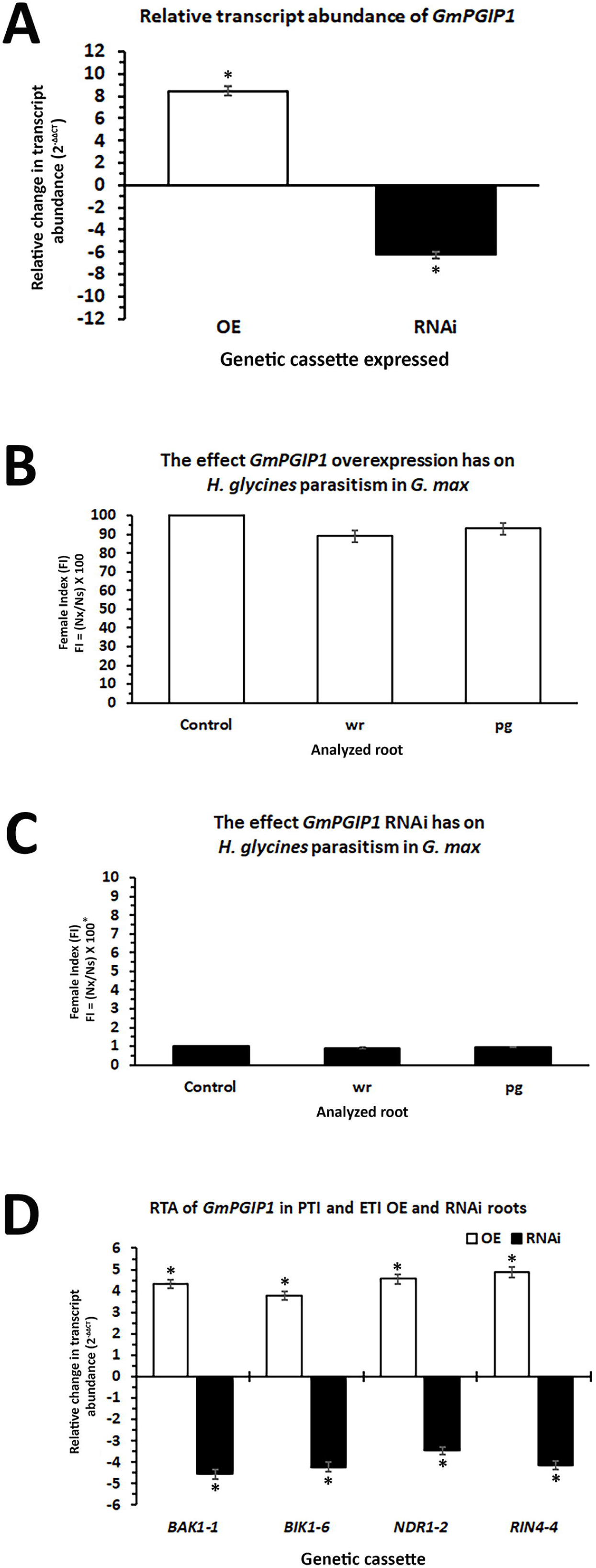
Altered *GmPGIP1* expression, effect on *H. glycines* parasitism and RTA in transgenic roots of PTI and ETI genes. **A.** The change in *GmPGIP1* RTA (OE-white histogram, RNAi-black histogram), calculated by 2^-ΔΔCT^ (Livak and Schmittgen 2001). **B**. The FI for *GmPGIP1*-OE roots as compared to their respective pRAP15-*ccd*B OE control (Golden et al. 1970). The statistical significance was determined by the MWW Rank-Sum Test, p < 0.05 (Mann and Whitney, 1947). **C**. The FI for *GmPGIP1*-RNAi roots as compared to their respective pRAP17-*ccd*B RNAi control (Golden et al. 1970). The statistical significance was determined by the MWW Rank-Sum Test, p < 0.05 (Mann and Whitney, 1947). **D**. *GmPGIP* RTA in the root RNA of the ETI genes *GmNDR1-1* and *GmRIN4-4* ETI as compared to the pRAP15-*ccd*B OE and pRAP17-*ccd*B RNAi controls, respectively. *GmPGIP1* RTA was calculated by 2^-ΔΔCT^ to determine RTA in *GmBAK1-1*, *GmBAK1-6*, *GmNDR1-2*, *GmRIN4-4*-OE (white histograms) and -RNAi (black histograms) expressing roots as compared to their respective controls (Livak and Schmittgen 2001). (*), statistical significance of p < 0.05, Student’s *t*-test (Yuan et al. 2006).

### Pairwise comparison of *Gm*PGIP1 and *Gm*PGIP11

An examination of the 335 aa *GmPGIP1* and 351 aa *GmPGIP11* accomplished through LM of control pericycle cells and syncytia undergoing two different forms of a defense response showed only that *GmPGIP11* was expressed within the syncytium under the study conditions. When examining *GmPGIP11* and *GmPGIP1*, only the overexpression of *GmPGIP11* led to defense to *H. glycines*. The outcome indicated that perhaps there were important aa signatures that could be present in *Gm*PGIP11 and not in *Gm*PGIP1, or vice versa, that relate to these differences in function. A conserved domain description (CDD) is provided to identify LRRs that are known to be present in PGIP proteins and act in the defense process (**Supplemental Figure 7**). A pairwise comparison identified protein signatures found in *Gm*PGIP11 that were not present in *Gm*PGIP1 (**Figure 8**). The 6 *Gm*PGIP11 LRRs demonstrated sequence divergence from *Gm*PGIP1 (**Figure 8**).

**Figure 8.**
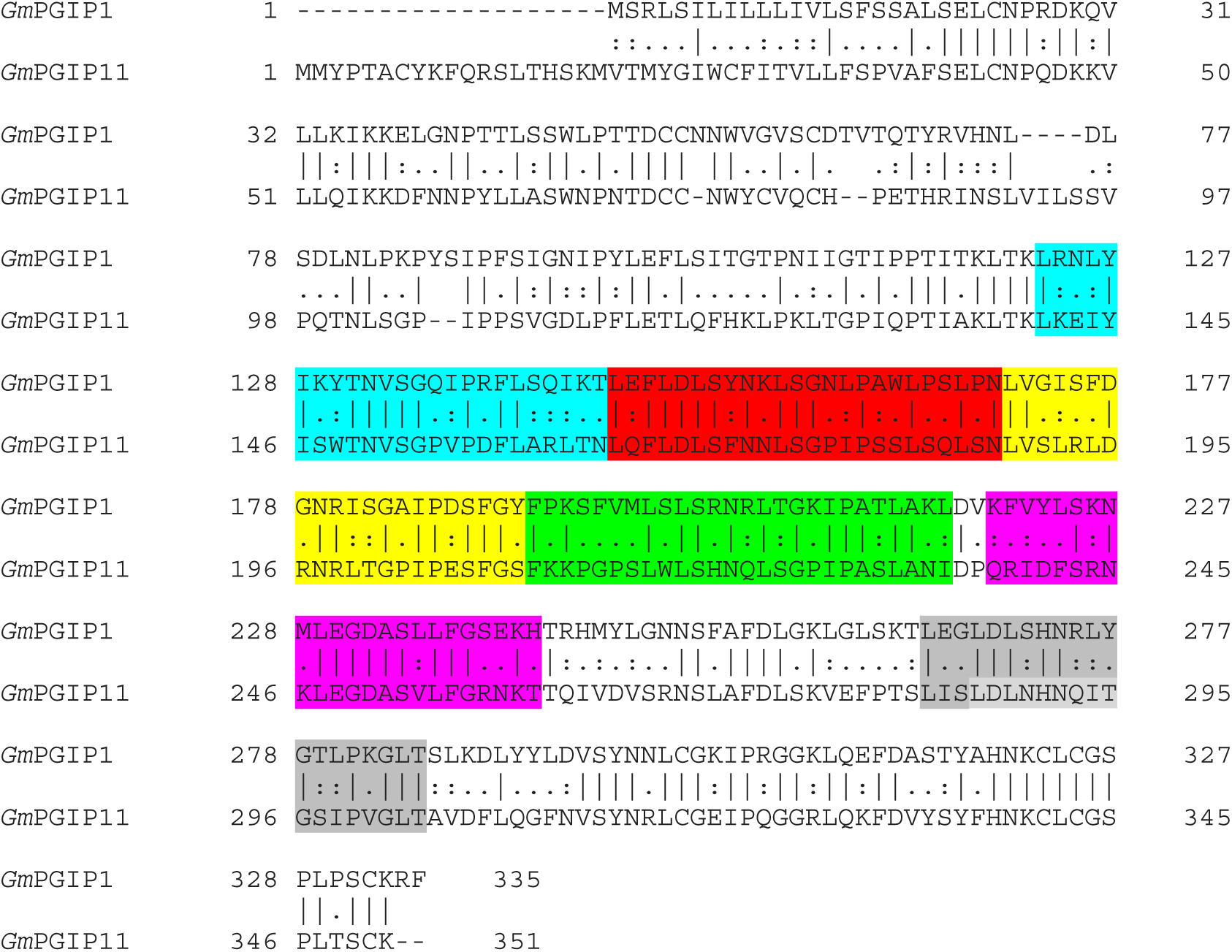
Pairwise comparison of *Gm*PGIP1 and *Gm*PGIP11. The analysis used the parameters of NEEDLE in EMBOSS (Needleman and Wunsch, 1970; Madeira et al. 2019). Details can be found in the Materials and Methods subsection: Pairwise protein sequence comparison. The sequential LRR repeats were highlighted in cyan, red, yellow, green, magenta, and gray. Please see the Materials and Methods, subsection: Pairwise protein sequence comparison, for details.

### 3-dimensional comparison of GmPGIP1 and GmPGIP11

Higher order structural arrangements perform important roles in protein-protein interactions between plants and pathogens (Gao et al. 2024). The dissimilarity in aa composition that exists between *Gm*PGIP11 and *Gm*PGIP1 indicated that there could be important differences in their 3-dimensional structures that may underlie their differences in defense function. A 3-dimentional comparison was performed between *Gm*PGIP11 and *Gm*PGIP1 to provide a basis for comparison (**Figure 9**).

**Figure 9.**
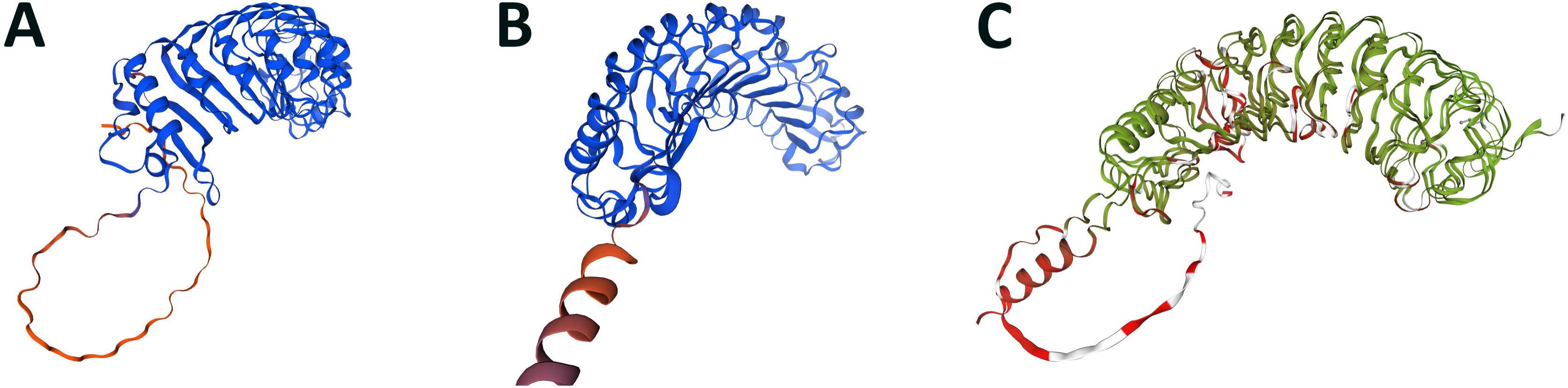
Pairwise 3-dimensional reconstruction and comparison of *Gm*PGIP11 and *Gm*PGIP1. For details, please see Materials and methods subsection: “3-dimensional modeling of *Gm*PGIP proteins. **A.** *Gm*PGIP11. **B**. *Gm*PGIP1. **C**. Comparative superimposed model of *Gm*PGIP11 and *Gm*PGIP1. Homology modeling of the protein sequences was performed using SWISS-MODEL under default settings. Red, identical; green very similar.

### The determination of PGIP protein families in agriculturally important crops

Plant protection from pathogens is an ongoing challenge (Thomas et al. 2023). An analysis is presented here that identified predicted PGIP protein families of important crop plants. The protein counts determined from the translated *PGIP*s from each plant species which included the products of splice variants from single genes are presented (**Table 3; Supplemental Tables 1, 4**).

**Table 3.** PGIP paralogs present in select crop species.

| Crop number | Species | genome | PGIP proteins |
| --- | --- | --- | --- |
| 1 | *Amborella trichopoda* | Amborella trichopoda v1.0 | 2 |
| 2 | *Amaranthus hypochondriacus* | Amaranthus hypochondriacus v2.1 | 6 |
| 3 | *Beta vulgaris* | Beta vulgaris EL10_1.0 | 9 |
| 4 | *Chenopodium quinoa* | Chenopodium quinoa v1.0 | 22 |
| 5 | *Spinacia oleracea* | Spinacia oleracea Spov3 | 6 |
| 6 | *Coffea arabica* | Coffea arabica v0.5 | 15 |
| 7 | *Daucus carota* | Daucus carota v2.0 | 9 |
| 8 | *Helianthus annuus* | Helianthus annuus r1.2 | 8 |
| 9 | *Lactuca sativa* | Lactuca sativa V8 | 13 |
| 10 | *Mimulus guttatus* | M.guttatus_TOL v5.0 | 10 |
| 11 | *Olea europaea* | Olea europaea v1.0 | 5 |
| 12 | *Solanum lycopersicum* | Solanum lycopersicum ITAG4.0 | 5 |
| 13 | *Solanum tuberosum* | Solanum tuberosum v6.1 | 3 |
| 14 | *Eucalyptus grandis* | Eucalyptus grandis v2.0 | 9 |
| 15 | *Vitis vinifera* | Vitis vinifera v2.1 | 4 |
| 16 | *Arachis hypogaea* | Arachis hypogaea v1.0 | 17 |
| 17 | *Castanea dentata* | Castanea dentata v1.1 | 5 |
| 18 | *Cicer arietinum* | Cicer arietinum v1.0 | 8 |
| 19 | *Cucumis sativus* | Cucumis sativus v1.0 | 3 |
| 20 | *Fragaria vesca* | Fragaria vesca v4.0.a2 | 4 |
| 21 | *Malus domestica* | Malus domestica v1.1 | 8 |
| 22 | *Medicago truncatula* | Medicago truncatula Mt4.0v1 | 23 |
| 23 | *Phaseolus vulgaris* | Phaseolus vulgaris v2.1 | 9 |
| 24 | *Prunus persica* | Prunus persica v2.1 | 5 |
| 25 | *Quercus rubra* | Quercus rubra v2.1 | 7 |
| 26 | *Trifolium pratense* | Trifolium pratense v2 | 16 |
| 27 | *Carya illinoensis* | Carya illinoensis v1.1 | 3 |
| 28 | *Vigna unguiculata* | Vigna unguiculata v1.2 | 10 |
| 29 | *Linum usitatissimum* | Linum usitatissimum v1.0 | 9 |
| 30 | *Manihot esculenta* | Manihot esculenta v8.1 | 6 |
| 31 | *Carica papaya* | Carica papaya ASGPBv0.4 | 3 |
| 32 | *Theobroma cacao* | Theobroma cacao v2.1 | 4 |
| 33 | *Arabidopsis thaliana* | Arabidopsis thaliana TAIR10 | 6 |
| 34 | *Schrenkiella parvula* | Schrenkiella parvula v2.2 | 6 |
| 35 | *Brassica oleracea capitata* | Brassica oleracea capitata v1.0 | 19 |
| 36 | *Brassica rapa* | Brassica rapa FPsc v1.3 | 19 |
| 37 | *Sinapis alba* | Sinapis alba v3.1 | 26 |
| 38 | *Gossypium hirsutum* | Gossypium hirsutum v2.1 | 6 |
| 39 | *Citrus sinensis* | Citrus sinensis v1.1 | 4 |
| 40 | *Ananas comosus* | Ananas comosus v3 | 5 |
| 41 | *Dioscorea alata* | Dioscorea alata v2.1 | 9 |
| 42 | *Musa acuminata* | Musa acuminata v1 | 5 |
| 43 | *Hordeum vulgare* | Hordeum vulgare r1 | 13 |
| 44 | *Oryza sativa* | Oryza sativa v7.0 | 9 |
| 45 | *Triticum aestivum* | Triticum aestivum v2.2 | 19 |
| 46 | *Brachypodium distachyon* | Brachypodium distachyon v3.2 | 7 |
| 47 | *Miscanthus sinensis* | Miscanthus sinensis v7.1 | 13 |
| 48 | *Sorghum bicolor* | Sorghum bicolor v3.1.1 | 8 |
| 49 | *Zea mays* | Zea mays RefGen_V4 | 9 |
| 50 | *Panicum hallii* | Panicum hallii v3.2 | 7 |
| 51 | *Vaccinium darrowii* | V.darrowii v1.2 | 13 |
| TOTAL |  |  | 469 |

### *PGIP* gene expression in other plant pathosystems

Ghaemi et al. (2020) investigated the susceptible and resistant reactions between *B. vulgaris* (sugar beet [SB]) and *H. schachtii* (beet cyst nematode [BCN]). The analyses identified significantly differentially expressed genes (DEGs) of sugar beet in the compatible sugar beet-BCN interaction. Included among the genes was a *B. vulgaris PG BV4G02530* (-1.49-fold). The authors also found a *B. vulgaris PG* significantly increased in its RTA related to cell wall architecture in the compatible SB-BCN interaction that included *BV4G02530* (1.36-fold). The authors then found significantly expressed sugar beet *PG*s in the incompatible SB-BCN interaction at 4 dpi *BV6G09860* (-1.10-fold). The authors also found significantly expressed sugar beet *PG*s in the incompatible SB-BCN interaction at 10 days after infection (dai) including the *PG*s *BV8G03010* (1.38-fold), and *BV4G02530* (1.37-fold). The authors subsequently examined significantly expressed SB DEGs related to cell wall architecture in the incompatible SB-BCN interaction. No significantly expressed *PG*s were identified in the comparison of resistant and susceptible SB cultivars in uninfected and BCN-infected conditions. Furthermore, no sugar beet genes show a differential response to BCN infection when comparing the resistant and susceptible cultivar. Analyses then were done to determine transcripts derived from BCN genes detected during the compatible and incompatible interaction with SB. Analyses of the 4 and 10 dai infected *H. schachtii* susceptible samples identified the *PG*s *SRR1125017.170284*, *SRR1125017.174045*, and *SRR1125017.154506*. Analyses of the 4 and 10 dai *H. schachtii* infected resistant samples identified the *PG*s *SRR1125017.170284*, *SRR1125017.174045*, and *SRR1125017.154506*. No genes annotated as *PGIP*s were identified. The analyses of Ghaemi et al. (2020) may highlight the drawbacks of using whole root RNA extracts for gene expression studies that involve very specific sets of plant and pathogen cells and the advantage of the LM technique presented here.

### 3-dimensional comparison of the GmPGIP11 and B. vulgaris BvPGIP1 and BvPGIP2

A *B. vulgaris PGIP* was shown to be expressed in roots infected by *T. myopaeformis* during a resistant reaction and functions in resistant reactions toward a number of pathogens (Li and Smigocki, 2018). A comparison of the primary structures of *Gm*PGIP11 and *Bv*PGIP1 reveals the 384 aa *Bv*PGIP1 is 35.7% identical and 50.1% similar to *Gm*PGIP11. Furthermore, the 384 aa *Bv*PGIP2 is 35.5% identical and 50.2% similar to *Gm*PGIP11. *Bv*PGIP1 and *Bv*PGIP2 are 97.9% identical and 99.2% similar to each other. There are no predicted *N*- or *O*-glycosylation sites conserved between *Gm*PGIP11 and *Bv*PGIP1 or *Bv*PGIP2. To better understand these proteins, a 3-dimensional analysis was done that compared *Gm*PGIP11 to *Bv*PGIP1 and also *Gm*PGIP11 to *Bv*PGIP2 (**Figure 10**).

**Figure 10.**
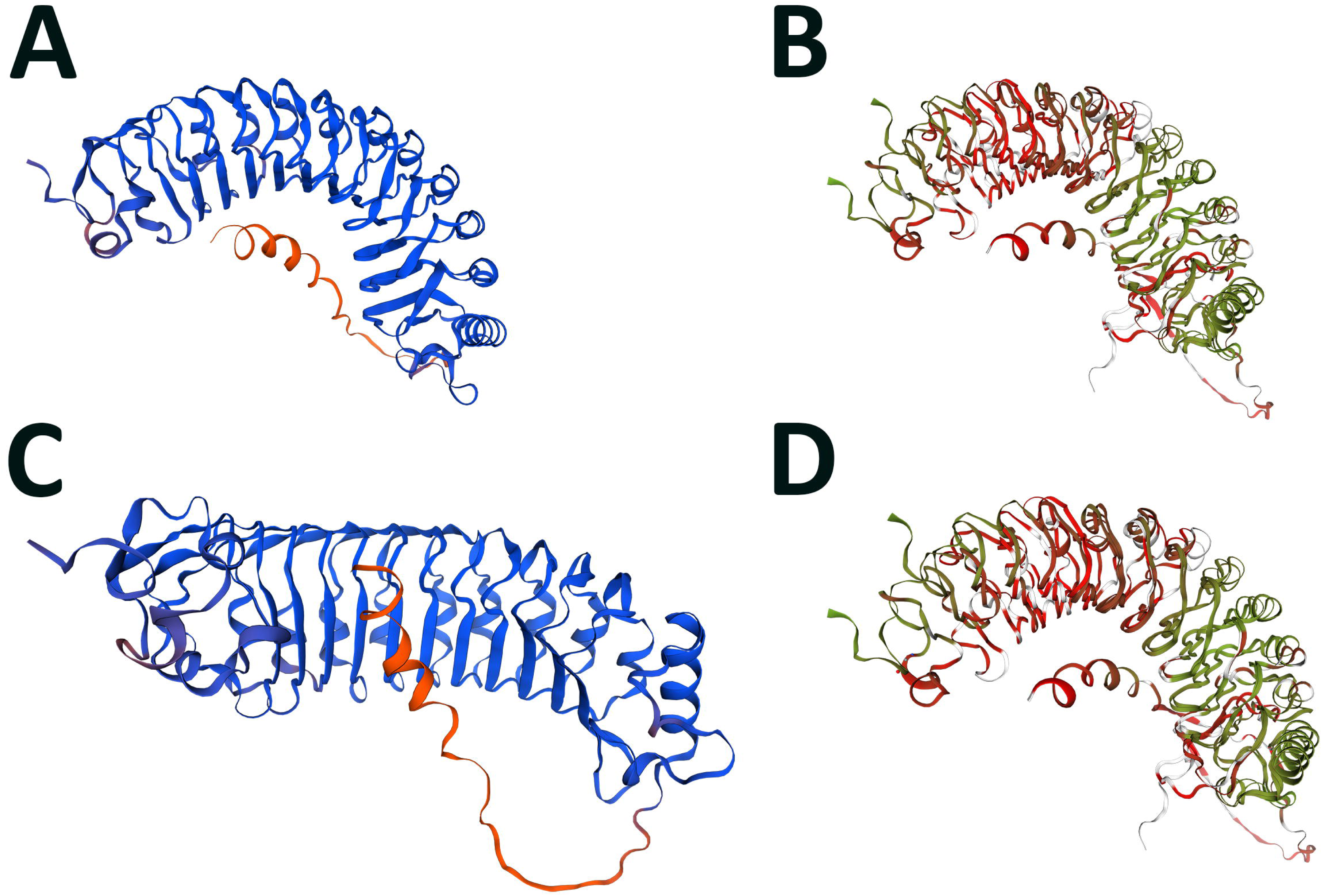
Pairwise 3-dimensional reconstruction and comparison of *Bv*PGIP1 and *Bv*PGIP2. For details, please see Materials and methods subsection: “3-dimensional modeling of *Gm*PGIP proteins. **A**. *Bv*PGIP1. **B**. Comparison of *Gm*PGIP11 and *Bv*PGIP1. **C**. *Bv*PGIP2. **D**. Comparison of *Gm*PGIP11 and *Bv*PGIP2. Homology modeling of the protein sequences was performed using SWISS-MODEL under default settings. Red, identical; green very similar. The *Bv*PGIP1 and *Bv*PGIP2 sequences, originally cloned from *B. vulgaris* genotypes sugar beet root maggot (SBRM) susceptible F1016 and SBRM resistant F1010, respectfully were obtained from Li and Smigocki, (2016).

## DISCUSSION

The analyses presented here were performed to evaluate a *GmPGIP* (*GmPGIP11*), found to be expressed in a nurse cell (syncytium) used by *H. glycines* as it would normally derive nourishment from during its parasitic life cycle. However, *GmPGIP11* was expressed during a defense response that led to the demise of *H. glycines* (Matsye et al. 2011). This expression contrasted with other *GmPGIP*s that could be studied under the experimental parameters presented here. Functional transgenic experimental evidence indicates *Gm*PGIP11 functions in the defense response that *G. max* has toward *H.* glycines. The results of the analyses presented here are discussed and put into the context of prior analyses.

### *GmPGIP*s belong to a small gene family

Mahalingam et al. (1999) identified a *Gm*PGIP protein sequence (AF130974.1 [AAD45503]), found in the *H. glycines* -resistant *G. max* _[PI 437654]_. These experiments were performed prior to the sequencing of the *G. max* _[Williams 82/PI 518671]_ genome, with no further functional transgenic studies reported (Mahalingam et al 1999; Schmutz et al. 2010). Bioinformatics analyses presented here determined that AF130974.1 was 100% identical to Glyma.08G079100.1. However, the AF130974.1 *Gm*PGIP sequence was missing the N-terminal, 19 aa polypeptide sequence MSKLSILFLLVLSFSSVLS. The next most closely related protein sequence to AF130974.1 was Glyma.05G123700.1 that had 72% identity. Consequently, AF130974.1 was most likely Glyma.08G079100. Subsequent analyses of *G. max* PGIPs identified Pgip1, Pgip2, Pgip3, and Pgip4, shown here, respectively (D’Ovidio et al. 2006). The D’Ovidio et al. (2006) studies showed the *G. max* Pgips to function effectively, broadly among different pathogen types, to impair pathogenicity. The effectiveness of these PGIPs in defense led to an examination of syncytium expression of soybean PGIPs in *G. max* _[Peking/PI 548402]_ and *G. max* _[PI 88788]_, each capable of an effective defense response to *H. glycines* parasitism (Caldwell et al. 1960; Matson and Williams, 1965). The analyses determined *G. max*_[Peking/PI 548402]_ and *G. max*_[PI 88788]_ express *GmPGIP11* (*Glyma.19G145200*) during their defense responses while not expressing them in control cells. Furthermore, *GmPGIP1*, *GmPGIP3*, *GmPGIP4*, *GmPGIP6*, and *GmPGIP10* were not expressed in the 0 dpi control, 3, or 6 dpi samples. Additionally, they were not found to be expressed in the 9 dpi time point sample. *GmPGIP11* was expressed in a 9 dpi time point *G. max* _[Peking/PI 548402]_ but not in *G. max* _[PI 88788]_. Cytological studies, however, have determined that the *G. max* defense response to *H. glycines* has concluded by 6 dpi, so it is possible that the collected cells were no longer transcriptionally active in *G. max*_[PI 88788]_ or the lack of expression has been controlled by some other process like apoptosis (Endo 1965, 1991). Gene expression assessment of *GmPGIP2*, *GmPGIP5*, *GmPGIP7*, *GmPGIP8*, and *GmPGIP9* could not be made by the analysis procedures presented here because they lacked probe sets on the microarray. *GmPGIP2*, *GmPGIP5*, and *GmPGIP8*, however, were predicted to lack a signal peptide, confirmed by Acharya et al. (2023) and may be undergoing a process of becoming genetically inert as described in the birth and death hypothesis presented by Kalunke et al. (2014). *GmPGIP2* is part of a locus that has 4 tandem *GmPGIP* copies (*GmPGIP1*, *GmPGIP2*, *GmPGIP3*, and *GmPGIP4*) on chromosome 5. *GmPGIP5* and *GmPGIP8* are part of 2 tandem duplications on chromosome that are interspersed by the N-acetyl-gamma-glutamyl-phosphate reductase/NAGSA dehydrogenase *Glyma.08G079000*. The *GmPGIP* of Mahalingam et al (1999) is *GmPGIP7* (*Glyma.08G079100*), but in their analysis it was expressed during mechanical wounding and during a susceptible reaction. D’Ovidio et al. (2006) identified their *Gmpgip1 AJ972660*, shown here as *Glyma.05G124000* (*GmPGIP4*). The *Gmpgip2 AJ972661* was shown here as *Glyma.05G123900* (*GmPGIP3*). The *Gmpgip3 AJ972662* was shown here as *Glyma.08G079100* (*GmPGIP7*). The *Gmpgip4 AJ972663* was shown here as *Glyma.08G079200.* (*GmPGIP8*). Expression in the *G. max* -*H. glycines* pathosystem under our parameters showed *GmPGIP3* and *GmPGIP4* lacked expression. Expression for *GmPGIP7* and *GmPGIP8* could not be determined under the analysis parameters. Our *GmPGIP* naming convention was presented in order to be comprehensive as it relates directly to the sequenced *G. max* genome in the order of the appearance of the genes, beginning with nucleotide 1 on chromosome 1. The evidence presented here supports the hypothesis that the expression of *GmPGIP11* is indicative that it functions in the defense response that *G. max* has to *H. glycines* parasitism. This hypothesis led to its cloning for functional transgenic studies in an attempt to understand any potential defense role.

### *G. max* PGIPs have features consistent with being secreted proteins

The defense response that *G. max* has toward *H. glycines* is dependent on its vesicular transport system (Matsye et al. 2011, 2012; Cook et al. 2012; Pant et al. 2014; Sharma et al. 2016, 2020; Liu et al. 2017; Lakhssassi et al. 2020a, b; Lawaju et al. 2020; Sharma et al. 2020; Song et al. 2021). These observations are supported by the results presented here that *Gm*PGIP11, with a level of specificity, impairs *H. glycines* parasitism. Analyses presented here and elsewhere report that *Gm*PGIPs have signal peptides, are *N*- and *O*-glycosylated, and have features that indicate they are transported through the secretion system (Acharya et al. 2023). The combination of the *GmPGIP11*-OE and RNAi results are compelling evidence they also function in defense in this studied pathosystem.

### *GmPGIP*s are expressed in *H. glycines*-parasitized root cells undergoing a defense response

The analysis presented here was based off of mining previously-generated data to determine whether members of the *G. max PGIP* gene family were expressed specifically during the defense response to *H. glycines*. This interest was because of the effectiveness that a root *PGIP* in *B. vulgaris* had on several pathogens (Li and Smigocki, 2018). The analysis identified that *GmPGIP11* was expressed specifically during 2 different forms of a defense response that *G. max* had toward the target pathogen, *H. glycines*. This expression contrasted to the expression of the control root cell, a population of pericycle cells, that lacked expression altogether. In contrast, the other *GmPGIP*s that had gene expression data available from RNA isolated from the nematode-parasitized root cell, lacked expression. The observation provided support that *GmPGIP11* could be an important candidate gene to study for a possible defense role in the *G. max*-*H. glycines* pathosystem.

### It is possible to express different *GmPGIP*s in the roots of *G max*

The pRAP series of plasmids were designed to study root pathogens and were particularly useful in the examination of genes expressed within the syncytium during a defense response (Matsye et al. 2012, Pant et al. 2014; Klink et al. 2009, 2021). The usefulness of the pRAP plasmids was expanded to include the study of *GmPGIP* genes. We showed that both *GmPGIP1* and *GmPGIP11*, when engineered into the pRAP15 overexpression plasmid, led to a significant increase in their RTAs as compared to their respective pRAP15-*ccd*B controls. In contrast, the engineering of *GmPGIP1* and *GmPGIP11*, into the pRAP17 RNAi plasmid, led to a significant decrease in their RTAs as compared to their respective pRAP17-*ccd*B controls. The specificity of the pRAP15 and pRAP17 plasmid’s ability to produce the expected result was demonstrated by the opposite outcome in the obtained RTA analyses (G*mPGIP*-OE: increased RTA, G*mPGIP*-RNAi: decreased RTA). The specificity of the functionality of the pRAP15 and pRAP17 plasmids was demonstrated in the outcomes of the pRAP15-*ccd*B OE and pRAP17-*ccd*B RNAi *GmPGIP11* FI analyses while no effect on *H. glycines* parasitism was observed in the *GmPGIP1* analyses. This aspect of the study was important because it is not always possible to obtain transformants when using the pRAP plasmids. For example, analyses of studied *G. max myosin XI* genes resulted in no transformants when using the pRAP15 overexpression plasmid while transformants using the pRAP17 RNAi plasmid were obtained (Austin et al. 2019). In that case, it was thought that balanced expression of the *myosin XI* was important for root development and that their overexpression may be toxic to root cell biology or root development (Austin et al. 2019). The usefulness of the plasmids in these studies has been tested in analyses of chlorophyll biosynthesis, defense signaling, cell wall biology, circadian rhythms, vesicle transport, and other cellular processes (Pant et al. 2014, Klink et al. 2021; Niraula et al. 2022b). Notably, vesicle transport relates to PGIPs (Haeger et al. 2020).

### *GmPGIP11* expression affects *H. glycines* parasitism

The hypothesis for the functional transgenic studies was that increased *GmPGIP11* RTA through overexpression changes the *H. glycines* -susceptible *G. max* _[Williams 82/PI 518671]_ so that it produces a defense response outcome resembling the *H. glycines*-resistant *G. max*_[Peking/PI 548402]_. In contrast a decrease in *GmPGIP11* RTA changes the *H. glycines*-resistant *G. max*_[Peking/PI 548402]_ so it produces the observed impaired defense response (i.e., susceptibility) of *G. max*_[Williams 82/PI 518671]_. The combined observed decrease in *H. glycines* parasitism in *GmPGIP11*-OE *G. max*_[Williams 82/PI 518671]_ roots leading to more *H. glycines*-resistant roots and increase in *H. glycines* parasitism in *GmPGIP-11*-RNAi in the *G. max*_[Peking/PI 548402]_ roots is accepted as evidence that the targeted *GmPGIP11* gene functions in the defense response (Pant et al. 2014). The analyses identified a 75.77% to a 77.81% decrease in *H. glycines* parasitism in *GmPGIP11*-OE *G. max*_[Williams 82/PI 518671]_ in cysts per wr and pg analyses, respectively. In contrast, the FI analysis identified a 7.5 to 7.7-fold increase in *H. glycines* parasitism in the *GmPGIP-11*-RNAi in the *G. max*_[Peking/PI 548402]_ in cysts per wr and pg analyses, respectively. The results demonstrated that the *GmPGIP11* gene functions in the defense process as shown for other syncytium-expressed genes with features of secreted proteins (Pant et al. 2014; Klink et al. 2017; 2021). The results are consistent with those of Shah et al. (2017). Shah et al. (2017) demonstrated the *A. thaliana AtPGIP1*, one of its 2 *PGIP*s, is strongly increased in its RTAs in response to *H. schacttii* root invasion, a process associated with mechanical damage which produces elicitors that activate defense signaling. Furthermore, *AtPGIP1* loss-of-function mutants were susceptible to *H. schachtii* parasitism but had no effect on *Meloidogyne incognita* (root knot nematode [RKN]) parasitism (Shah et al. 2017). Complementation studies demonstrated that overexpressed *AtPGIP1* resulted in a restoration of resistance to *H. schachtii* parasitism but had no effect on RKN parasitism (Shah et al. 2017). As a protein having a number of features consistent with secreted proteins, it is likely that the *Gm*PGIP11 protein is transported through the secretion system where it arrives at its site of function leading to the impairment of *H. glycines* parasitism. An important *H. glycines* resistance gene, *rhg1*, is believed widely to be alpha soluble *N*-Ethylmaleimide-Sensitive Factor Attachment Protein (α-SNAP) which functions with SNAP Receptor (SNARE) proteins at the fusion step of vesicle and target membranes (Matsye et al. 2011, 2012; Cook et al. 2012; Zick et al. 2015). This role is consistent with defense functions found for the vesicle transport system and secreted proteins, broadly, in plants for SNARE, and proven defense roles in *G. max* (Collins et al. 2003; Matsye et al. 2011, 2012; Cook et al. 2012; Pant et al. 2014; Sharma et al. 2016, 2020; Liu et al. 2017; Austin et al. 2019; Lakhssassi et al. 2020a, b; Lawaju et al. 2020; Song et al. 2021). Experiments show *Gm*MAPK-induced genes that were expressed within the syncytium and have signal peptides functioned effectively during the defense response to *H. glycines* parasitism, indicating PTI and/or ETI defense branches regulate their expression (Niraula et al. 2020; Acharya et al. 2024). The importance of signal peptide containing proteins was demonstrated in *A. thaliana* mutants of signal peptide peptidase (Han et al. 2009).

### *GmPGIP11* expression does not alter root growth

The results for the *GmPGIP11* FI analyses of cysts per whole root and per gram of root were obtained during the course of the experiments. The results, confirmed by the analysis of the root mass, showed *GmPGIP11*-OE and *GmPGIP11*-RNAi did not affect root growth to a statistically significant level (p > 0.05, MWW rank sum test). The results were consistent with the delimited expression that is found for *GmPGIP11*, indicating a localized, specialized role in defense that does not affect the rest of the root system. The result indicated any signals emanating from *GmPGIP* expression were not interfering with root development in a global manner. Regarding *PGIP*s in other plant systems, changes in growth were not observed in *Gossypium hirsutum*, *Oryza sativa*, *N. tabacum* or *T. aestivum* by their altered expression (Janni et al. 2008; Ferrari et al. 2012; Wang et al. 2013, 2015, 2018; Kalunk et al. 2015; Liu et al. 2018). The analysis presented here examined *GmPGIP11* expression in relation to the PTI genes *GmBAK1-1* and *GmBIK1-6*, as well as the ETI genes *GmNDR1-1*, and *GmRIN4-4* that were expressed in the syncytium during a defense response but whose OE and RNAi also did not affect root growth to a statistically significant level (Pant et al. 2014; McNeece et al. 2017; Klink et al. 2021). An examination of the 32 *GmMAPK*s revealed OE of *GmMAPK2*, and *GmMAPK5-3* which functioned during the defense response to *H. glycines,* decreased root mass to a statistically significant level (McNeece et al. 2019). In contrast, silencing of the defense *MAPK*s *GmMAPK3-2* and *GmMAPK4-2* increased root mass to a statistically significant level (McNeece et al. 2019). *GmMAPK*s not functioning in defense but whose OE affect root mass to a statistically significant level were limited to those experiments that resulted in a decrease in root mass, shown for *GmMAPK4-2*, *GmMAPK5-1*, *GmMAPK7*, *GmMAPK11-2*, *GmMAPK13-2*, *GmMAPK16-1*, and *GmMAPK19* (McNeece et al. 2019). In no case did RNAi of *GmMAPK*s not functioning defense affect root mass (McNeece et al. 2019). The results obtained for *GmPGIP11* expression, and limited to *GmPGIP11*, point toward localized expression as a means to achieve a potent response to a biotic challenge (*H. glycines* parasitism) while avoiding potential problems caused by global expression of genes functioning in a specialized function (Pino et al. 2007; Gwak et al. 2017). Perhaps the induced expression of *GmPGIP11* overcomes pathogen-secreted PGs or effectors that target them (Zhang et al. 2021; Wei et al. 2022).

### *GmPGIP11* RTA is affected by PTI and ETI component expression

PTI and ETI function in plants, coordinately, to produce a defense response (Jones and Dangl, 2006; Ngou et al. 2021). PTI functions in the *G. max*-*H. glycines* pathosystem as *GmBIK1-6* is expressed in syncytia undergoing a defense response to *H. glycines* with subsequent OE and RNAi experiments showing a reduction in parasitism by nearly 90% (Matsye et al. 2011; Pant et al. 2014). Subsequent experiments showed that *GmBIK1-6* regulates *GmMAPK3-1* and *GmMAPK3-2* expression (McNeece et al. 2019). Complimentary analyses demonstrated that the *GmBAK1-1*, one of 19 *BAK1*-like genes in *G. max*, was expressed in syncytia undergoing the *H. glycines* defense response and functioned in defense (Klink et al. 2021). *GmBAK1-1* also regulated the expression of *GmMAPK3-1* and *GmMAPK3-2* which indicated a role in PTI-mediated defense (Klink et al. 2021). Aljaafri et al. (2017) demonstrated that the bacterial effector harpin functioned effectively to activate a defense response to different nematodes that parasitize *G. max,* including *H. glycines, Meloidogyne incognita,* and *Rotylenchulus reniformis* (Wei et al. 1992). Subsequent experiments demonstrated that the *G. max GmNDR1-1*, a transcriptional target of harpin, became induced in its expression as a consequence of a topical harpin treatment (McNeece et al. 2017). *GmNDR1-1* functioned effectively to suppress *H. glycines* parasitism while also inducing the expression of proven defense genes (McNeece et al. 2017). In a broader understanding of this genetic pathway, *GmNDR1-1* was shown to induce the RTA of *GmMAPK3-1* whose induced expression increased the RTAs of a number of proven defense genes. For example, *MAPK3-1* positively regulated the dominant *Resistance to heterodera glycines 4* (*Rhg4*) *serine hydroxymethyltransferase-5* (*GmSHMT-5*) (*Glyma.08G108900*), *xyloglucan endotransglycosylase-hydrolase 43,* (*GmXTH43*) (*Glyma.17G065100*), *reticuline oxidase-40* (*GmRO-40*) (*Glyma.15G132800*), *galactinol synthase-3* (*GmGS-3*) (*Glyma.19G227800*), secreted *pathogenesis related 1-6* (*GmPR1-6*) (*Glyma.15G062400*), *GmMAPK3-2*, and *GmNDR1-1* (McNeece et al. 2019). Furthermore, *GmMAPK3-2* positively regulated the proven defense genes including the *NPR1* co-transcriptional regulator *GmTGA2-1* (*Glyma.10G296200*), *GmRO-40*, *GmSHMT-5*, *GmNPR1-2*, *GmMAPK3-1*, and *GmPR1-6*. These results were confirmed with RNAi of *GmMAPK3-1* and *GmMAPK3-2* that led to a decrease in the RTAs of these same genes, respectively, with the transgenic roots being accompanied by susceptibility to *H. glycines* (McNeece et al. 2019). These observations were consistent with the results presented here that showed the overexpression of the PTI genes *GmBAK1-1* and *GmBIK1-6* increased *GmPGIP11* RTA while their RNAi decreased it. Furthermore, these observations were consistent with the results presented here that showed the overexpression of the ETI genes *GmNDR1-1* and *GmRIN4-4* increased *GmPGIP11* RTA while their RNAi decreased it.

### *GmPGIP* expression is regulated by *GmMAPK*s

An analysis of RNA seq data generated for *GmMAPK3-1* and *GmMAPK3-2* overexpressing and RNAi roots in comparison to their controls, respectively, were performed (Alsherhi et al. 2018). The expression data for the *GmPGIP* gene family was extracted and analyzed here. The *GmMAPK3-1*-OE root RNA sample increased RTAs for *GmPGIP1*, *GmPGIP3*, *GmPGIP4*, *GmPGIP5*, and *GmPGIP6*. The *GmMAPK3-1*-RNAi root RNA samples did not yield a statistically significant change in their RTAs as compared to the control. The *GmMAPK3-2*-OE RNA sample increased RTAs for *GmPGIP1*, *GmPGIP3*, *GmPGIP4*, and *GmPGIP6*. The *GmMAPK3-2*-RNAi also did not yield a statistically significant change in their RTAs as compared to the control. Notably, through the LM syncytium gene expression analyses, none of these *GmPGIP*s were observed to be expressed during the defense response at 3 or 6 dpi or in control cells. The results indicated that the expression of these *GmPGIP* genes may be detrimentally affected by the pathogen and resembling some of the observations made by Mahalingam et al. (1999). It appears as though *GmPGIP11* did not increase its RTA through induced *GmMAPK3-1* or *GmMAPK3-2* expression. This observation is surprising since the overexpression of *GmBAK1-1*-OE and *GmBIK1-6*-OE increased *GmPGIP11* RTA while *GmBAK1-1*-RNAi and *GmBIK1-6*-RNAi decreased *GmPGIP11* RTA. However, there was substantial overlap observed between PTI and ETI in *G. max* (McNeece et al. 2019). Furthermore, more recent experiments confirmed this overlap (Yuan et al. 2021; Ngou et al. 2021; Nguyen et al. 2021). In *A. thaliana*, After ETI, PTI proteins including the BIK1 can function in pathways leading to the induction of the expression of a number of downstream genes acting in defense that may not require MAPKs (Ngou et al. 2021). Similar observations were made in the *G. max*-*H. glycines* pathosystem (Pant et al. 2014; McNeece et al. 2019). The *A. thaliana* genes included glutamate-like receptors (GLRs), calcium-dependent protein kinases (CDPKs), cyclic nucleotide-gated channels (CNGCs), cysteine-rich receptor-like kinases (CRKs), leucine-rich repeat receptor-like proteins (LRR-RLPs), and leucine-rich-repeat receptor-like protein kinases (LRR-RLKs) (Ngou et al. 2021). The results indicated the pathway to regulated expression was not necessarily dependent on MAPKs and could be pathogen-dependent as shown in the *G. max* -*H. glycines* pathosystem (Klink et al. 2011). Furthermore, pathogen effectors impair PTI and ETI in a race-dependent manner which likely is the case for *H. glycines,* supported by unique genomic compositions found in separately-sequenced individuals. The induced expression of *GmPGIP11* appeared to be specific and did not seem to rely on *GmMAPK3-1* or *GmMAPK3-2* under the study conditions. This point was important due to recent observations that showed the relationship between components involved in the common symbiosis pathway and defense to *H. glycines* involving specific MAPKs (Khatri et al. 2022).

### Induced *GmPGIP* expression does not presage a defense function

*GmMAPK3-1* and *GmMAPK3-2* overexpression was capable of inducing *GmPGIP1*, *GmPGIP3*, *GMPGIP4*, *GmPGIP5*, and *GmPGIP6* expression. The results contrast with the LM expression data obtained for these genes including *GmPGIP1*, *GmPGIP3*, *GmPGIP4*, and *GmPGIP6* showing they lacked expression. *GmPGIP5* does not have a probe set fabricated onto the microarray and could not be examined by the LM experimental procedures. The results indicated the inductive signals that led to increased *GmPGIP11* expression, while involving PTI and ETI components as shown by its altered RTA in the transgenic *GmBAK1-1*, *GmBAK1-6*, *GmNDR1-1*, and *GmRIN4-4* root RNA, may be different that those obtained for *GmPGIP1*, *GmPGIP3*, *GmPGIP4*, and *GmPGIP6*. The LM experiments that identified the presence of *GmPGIP11* employed *H. glycines* -infected roots that were not transgenic while the RT-qPCR experiments examining transgenic roots undergoing overexpression or RNAi of *GmMAPK3-1*, *GmMAPK3-2*, *GmBAK1-1*, *GmBAK1-6*, *GmNDR1-1*, and *GmRIN4-4* were not infected, complicating direct comparison. The results showed the presence of *GmPGIP11* transcript in the syncytia undergoing a defense response did presage its defense function, while the absence of *GmPGIP1* did presage a lack of a defense function against *H. glycines* parasitism. The different *G. max PGIP* s likely have different defense spectra in relation to the pathogen of interest. The *Gm*PGIP1 presented here was not the same Gmpgip1 examined in the analyses of D’Ovidio et al. (2006), nor any of the others.

### Comparative primary structure analysis of *Gm*PGIP1 and *Gm*PGIP11

A comparative analysis of *Gm*PGIP1 and *Gm*PGIP11 has revealed the 358 aa protein alignments are 46.4% (166/358) identical, 63.7% (228/358) similar, and having 8.4% (30/358) gaps. There is a 1 aa N-terminal extension found only in *Gm*PGIP11. There are additional indels of 2 and 3 amino acids found outside of the LRR domains in the N-terminal region with an additional 2 aa indel found at the C-terminus. Protein alignments between the *G. max* PGIPs show the locations of the *N*- and *O*-glycosylation sites with some of them being conserved (Nguema-Ona et al. 2014; Acharya et al. 2023). *Gm*PGIP11 has the NVSG *N*-glycosylation site at a similar position to the *Gm*PGIP1 NVSG but, otherwise, The two proteins do not share *N*-or *O*-glycosylation sites, possibly contributing to their specificity and/or different roles (Acharya et al. 2023). Protein folding prediction models have revealed similarities and differences between the different *G. max PGIP*s. The importance of the folding likely relates to substrate (PG) binding.

### Comparative 3-dimentional protein modeling of *Gm*PGIP1 and *Gm*PGIP11

A 3-dimensional protein folding modeling of *Gm*PGIP1 as compared to *Gm*PGIP11 was performed to better understand the different outcomes obtained in the functional transgenic studies. The analyses produced a primary aa sequence alignment that was analyzed for identical and similar aa positions (Waterhouse et al. 2018). The produced alignment led to the generation of a 3-dimensional structure for both *Gm*PGIP11 and *Gm*PGIP1. Furthermore, the 3-dimensional models were superimposed on each other, allowing for visualization of the identical, similar, and non-similar regions. The produced alignment identified identical, similar, and non-similar regions that would relate to *O*- and *N*-glycosylation, protein features that are important to protein function, including defense (Kingsley et al. 1986; Hartweck et al. 2002; Koiwa et al. 2003; Chen et al. 2006). *N*-glycosylation has a selective and critical role in relation to different layers of plant immunity that appear to function through membrane-localized regulator quality control (Jin et al. 2007; Saijo et al. 2009). It is possible that the presentation of glycosylation mediated by the 3-dimensional structure of *Gm*PGIP11 relates to its role in defense to *H. glycines* parasitism.

### PGIPs are found broadly among important crop plants

Globally-important agricultural systems are experiencing changes in climate conditions (Tilman et al. 2011; Burkhead and Klink, 2018; Ray et al. 2019). Kalunk et al. (2015) have identified PGIPs from several plant species, demonstrating their broad existence. The analysis of the 50 additional proteomes results in the identification of their PGIPs with a low of 2 in *A. trichopoda* to a high of 26 for *S. alba.* The significance of *A. trichopoda* is that plant phylogeneticists place it at the base of the flowering plants in a monotypic order, the Amborellales. In contrast, *S. alba* is a member of the Brassicales that comprise approximately 17 families, 398 genera, and 4,450 species. The analyses of the genomes examined here provide a more extensive candidate pool of genes whose expression could be tested for functions in plant defense and could result in the generation of durable defense processes that perhaps could otherwise fail due to the pathogen and/or climatic factors (Jablonska et al. 2007; Tilman et al. 2011; Burkhead and Klink, 2018; Ray et al. 2019; Chiu et al. 2021). PGIPs such as *Gm*PGIP11 shown here have great potential not only to impair pathogen invasion but mitigate the effects of climate change (Rathinam et al. 2020). Conditions relevant to climate change do have large scale effects on plant roots (Sriden and Charoensawan, 2022). The experiments presented here for the root relates to a greatly understudied plant organ regarding climate change and defense, in particular, parasitic nematodes and may be relevant to other root pathogens such as insects (Ammati et al. 1986; Cap et al. 1993; Yaghoobi et al. 1995; Veremis and Roberts, 1996; Ammiraju et al. 2003; Claverie et al. 2004; Yaghoobi et al. 2005; Cooper et al. 2005; Marques et al. 2015; Li and Smigocki, 2018; Du et al. 2020; Alkharouf et al. 2024).

### *PGIP* gene expression in other plant pathosystems

Experiments demonstrated root expression of the *B. vulgaris PGIP* s *BvPgip1* and *BvPgip2* during a root infection by *Tetanops myopaeformis*, the sugar beet root maggot (SBRM) insect (Li and Smigocki, 2018; Alkharouf et al. 2024). The analysis then led to a literature search that showed Ghaemi et al. (2020) analyzed the infection that *B. vulgaris* had toward the root pathogens *H. schachtii* and *M. incognita.* The observations supported the hypothesis that *GmPGIP11* could function in the defense process that *G. max* has toward *H. glycines.* The observations also led to comparative studies of the *G. max* and *B. vulgaris* PGIPs to determine what similarities they may have.

### Comparative analysis of *Gm*PGIP11 to the sugar beet *Bv*PGIP1 and *Bv*PGIP2 defense PGIPs

A functional transgenic analysis of the *B. vulgaris BvPgip1* and *BvPgip2* revealed its heterologous transgenic expression in *N. benthamiana* produces resistance to fungal and insect shoot pathogens (Li and Smigocki, 2018). These observations provided an opportunity to understand any structural elements in *Bv*PGIP1 or *Bv*PGIP2 in common to *Gm*PGIP11 as they may relate to the resistance outcome. The 384 aa *Bv*PGIP1 is 35.7% identical and 50.1% similar to *Gm*PGIP11. The 384 aa *Bv*PGIP2 is 35.5% identical and 50.2% similar to *Gm*PGIP11. *Bv*PGIP1 and *Bv*PGIP2 are 97.9% identical and 99.2% similar to each other. There are no predicted *N*- or *O*-glycosylation sites conserved between *Gm*PGIP11 and *Bv*PGIP1 or *Bv*PGIP2. Experiments to transgenically express *BvPGIP1* or *Bv*P*GIP2* in *G. max* and *GmPGIP11* in *B. vulgaris* could provide important insight into their defense roles.

### Global and localized expression of PGIPs and the resistant reaction

The analysis here presented the use of LM as a means to isolate RNA from localized cell populations that relate to a specific cellular response to infection by a root pathogen. The experiment was used to identify a *PGIP* that has a specific expression pattern in cells undergoing the resistant reaction to *H. glycines* in 2 different *G. max* genotypes that are capable of a resistant reaction. However, over the era of transcriptomics many gene expression studies examined the interaction between *G. max* and *H. glycines*. The results of those analyses have been summarized here (**Figure 11; Supplemental Table 5)**. The analyses relating to the whole root includes 17 gene expression studies relating to the susceptible reaction that *G. max* whole root systems have to *H. glycines* spanning the 15 time points including 0, 8, and 12 hpi, 2, 3, 4, 5, 6, 8, 10, 12, 15, 16, 20, 30 dpi, and a pooled 3-21 dpi sample (Vaghchhipawala et al. 2001; Khan et al. 2004; Tucker et al. 2007; Klink et al. 2007; Ithal et al. 2007; Puthoff et al. 2007; Afzal et al. 2009; Tucker et al. 2011; Wan et al. 2015; Li et al. 2018; Song et al. 2019; Neupane et al. 2019; Guo et al. 2020; Miraeiz et al. 2020; Shi et al. 2021; Kofsky et al. 2021; Torabi et al. 2023). That discussion is followed by the gene expression papers relating to the resistant reaction that *G. max* whole root systems have to *H. glycines*, spanning 12 time points including (0, 8, 12 hpi, 2, 3, 4, 5, 6, 8, 10, 15, 30 dpi, and the combined 3-21 dpi timepoints analyses (Alkharouf et al. 2004; Klink et al. 2007; Wan et al. 2015; Li et al. 2018; Song et al. 2019; Neupane 2019; Miraeiz et al. 2020; Miraeiz et al. 2020; Guo et al. 2020; Shi et al. 2021; Kofsky et al. 2021; Chu et al. 2022; Torabi et al. 2023) (**Figure11**; **Supplemental Table 5)**. Analyses specifically limited to the syncytium have also been made. The analysis of the susceptible reaction span 2, 3, 5, 6, 8, 9, and 10 dpi (Klink et al. 2005, 2007b, 2009, 2010a, b, 2011; Ithal et al. 2007; Matsye et al. 2011). Analysis of the resistant syncytium include span 3, 6, 9 dpi (Klink et al. 2007b, 2009, 2010a,b, 2011; Kandoth et al. 2011; Matsye et al. 2011)) (**Figure 11**; **Supplemental Table 5)**. Presented here is information generated for *GmPGIP*s and *PG*s since PGIPs act on PGs. The results show the point in time when *PGIP*s and *PG*s have been observed to be expressed, the type of reaction (susceptible/resistant) were *PGIP*s and *PG*s are expressed, whether the expression is found broadly in the root or locally within the syncytium, and/or whether there is contrasting expression observed between the whole root and localized response. This conglomerate expression that has examined many gene expression analyses provides a window into the expression of *PGIP* and *PG* expression. It also lays the foundation for more broadly aimed analyses that may consider various important aspects of biology that relate to pathogenicity such as circadian rhythms along with more specific aspects that relate to root cell integrity and signaling (Niraula et al. 2020, 2021, 2022b).

**Figure 11.**
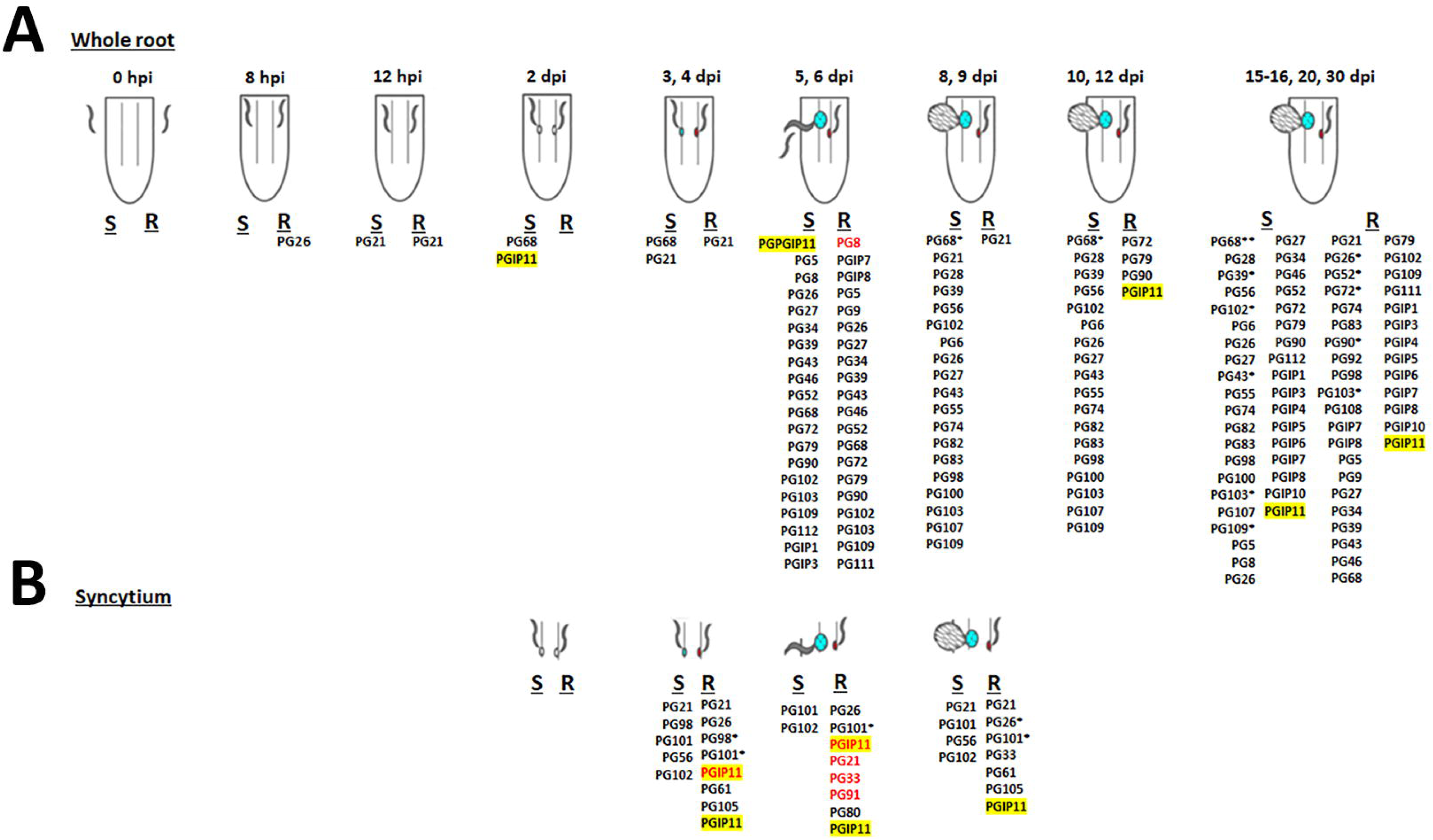
Reported *PGIP* and *PG* gene expression. Gene expression pertaining to the whole root susceptible reaction was extracted (Vaghchhipawala et al. 2001; Khan et al. 2004; Tucker et al. 2007; Klink et al. 2007; Ithal et al. 2007; Puthoff et al. 2007; Afzal et al. 2009; Tucker et al. 2011; Wan et al. 2015; Li et al. 2018; Song et al. 2019; Neupane et al. 2019; Guo et al. 2020; Miraeiz et al. 2020; Shi et al. 2021; Kofsky et al. 2021; Torabi et al. 2023). Gene expression pertaining to the whole root resistant reaction was extracted (Alkharouf et al. 2004; Klink et al. 2007; Wan et al. 2015; Li et al. 2018; Song et al. 2019; Neupane 2019; Miraeiz et al. 2020; Miraeiz et al. 2020; Guo et al. 2020; Shi et al. 2021; Kofsky et al. 2021; Chu et al. 2022; Torabi et al. 2023). (Klink et al. 2005, 2007b, 2009, 2010a, 2010b, 2011; Ithal et al. 2007; Matsye et al. 2011). Analysis of the resistant syncytium include span 3, 6, 9 dpi (Klink et al. 2007b, 2009, 2010a, 2010b, 2011; Matsye et al. 2011). Details of the analyses are available in those references. White syncytium, prior to cytological signs of outcome type (susceptible or resistant. Red, resistant syncytium. Blue, susceptible syncytium. Yellow highlight, *GmPGIP11*. Red lettering, decreased RTA. Black, increased RTAs. *, found to be expressed in multiple experiments. Note: the Klink et al. 2011 data is for *G. max*_[Peking/PI 540402]_ compared to *G. max*_[PI 88788]_ so a suppressed outcome means suppressed compared to another resistant reaction that may actually be induced compared to a control.

### Conclusion

*H. glycines* is a significant pathogen of *G. max,* devastating to agriculture on a global scale. *GmPGIP11* is expressed specifically in *G. max* root cells undergoing a defense response to *H. glycines* parasitism, indicating it may function in the defense response to stave off parasitism and, thus, a benefit to agriculture. The transgenic expression of *GmPGIP11* in *G. max*_[Williams 82/PI 518671]_ led to a statistically significant decrease in *H. glycines* parasitism in comparison to the pRAP15-*ccd*B control while not affecting root growth. In contrast, the transgenic expression of a *GmPGIP11* RNAi cassette in *G. max* _[Peking/PI 548402]_ led to a statistically significant increase in *H. glycines* parasitism in comparison to the pRAP17-*ccd*B control while not affecting root growth. The combination of the results indicates *Gm*PGIP11 has a defense role, in contrast to *Gm*PGIP1. PGIPs are expressed broadly in different angiosperms where they have defense roles. Most studies have focused on the roles that PGIPs have in the shoot. The analysis presented here reveals an important defense role in the root and that the expression relates to ETI and PTI signaling. Other studies that involve wild relatives of *G. max* will unravel additional layers of regulation (Zhang et al. 2017a, 2017b). Future, more comprehensive studies will likely provide additional details on the *GmPGIP* gene family as it relates to defense and other aspects of growth and development.

## STATEMENTS AND DECLARATIONS

### Ethics Declarations

The authors understand the ethics disclosure statement.

### Data Availability

All data is available in this work.

### Ethics Approval

An ethics approval is not required for the work.

## CONTRIBUTIONS

**SA** Performed the PGIP functional transgenic analyses, editing

**HT** Performed the PGIP functional transgenic analyses, editing

**RB** Performed the PGIP functional transgenic analyses, editing

**DM:** Bioinformatics, editing

**KL** Bioinformatics, editing

**NA** Bioinformatics, contributed to writing the PGIP aspect of the manuscript, bioinformatics

**VK** Generated research, wrote, edited the manuscript

## ACKNOWLEDGEMENTS

VK is thankful to the Department of Biological Sciences and the Department of Biochemistry, Molecular Biology, Entomology and Plant Pathology (BMBEPP) at Mississippi State University. Furthermore, VK is thankful to Gary Lawrence (retired) (BMBEPP) for all of his support over the years. Robert Nichols and Kater Hake of Cotton Incorporated are thanked for their support during this project. Yixiu (Jan) Pinnix BMBEPP is thanked for her technical support. Jeff Dean (BMBEPP) is thanked for his generosity of providing greenhouse, headhouse, storage, and field space for the experiments and maintenance of plant stocks. The authors thank Scott Willard, Wes Burger, George Hopper, and Reuben Moore, Mississippi Agricultural and Forestry Experiment Station (MAFES), and Mississippi State University for their support. The College of Arts and Sciences and MAFES at Mississippi State University have each provided crucial Special Research Initiative funding for this work. Funding is also provided by the USDA-ARS NP 8042-21220-262-000D project, and Beet Sugar Development Foundation under agreement number 58-3012-1-001-N to VK.

## Summary

For the whole root system studies, Vaghchhipawala et al. (2001) performed a PCR study of 12 select genes during *H. glycines* _[VL1]_ infection of susceptible *G. max* _[PI 89008]_. The listed genes included Catalase (Z12021), 2 Cyclins (D50868, X62820), Elongation factor EF-1α (X56856), 5 β-1,3-endoglucanases (U41323, U08405, U34755, U00730, U00731), Heat shock protein HSP70 (X62799), Hydroxymethylglutaryl CoAreductase (U97653), and Late embryonic abundant protein type 14 (U08108). *PGIP*s and *PG*s were not presented.

Alkharouf et al. (2004) performed an analysis of cDNA libraries spanning a total of 3,454 cDNA clones generated from 12-hours post infection (hpi), 2, 4, 6 and 8 days post infection (dpi). The analysis highlighted cDNA library transcripts accounting for at least t 0.3% of the expressed sequence tags (ESTs) from the *G. max*_[Peking/PI 548402]_ cDNA library generated from roots 12 h after inoculation with *H. glycines* _[race 3/NL1-RHp]_ that culminate in a resistant reaction. No *GmPGIP*s nor *GmPG*s meeting the criteria were identified (Alkharouf et al. 2004). An additional analysis of all cDNAs from the pooled libraries also did not identify any *GmPGIP* nor *GmPG* transcripts accounting for at least t 0.3% of the ESTs from the cDNA libraries generated (Alkharouf et al. 2004).

Khan et al. (2004) performed a microarray analysis revealing gene expression *G. max*_[Kent/PI 548586]_ roots susceptible to *H. glycines*_[race 3/ NL1-RHp]_ at 2 dpi. In those analyses, of the 1,305 ESTs printed on the cDNA microarray, 99 measured induced and one measured suppressed gene expression in *H. glycines*_[race 3/ NL1-RHp]_-inoculated *G. max*_[Kent/PI 548586]_ relative to the non-inoculated control (Khan et al. 2004). Neither *G. max PGIP*s nor *PG*s were highlighted in those analyses (Khan et al. 2004.

Alkharouf et al. (2006) performed a time course microarray analysis examining more than 6,000 cDNA inserts. The analysis examined 7 different time points (6, 12 hpi, 1, 2, 4, 6, and 8 dpi) during a susceptible reaction in *G. max*_[Kent/PI 548586]_ infected with *H. glycines*_[race 3/NL1-RHp]_, leading to the identification of hundreds of induced and suppressed genes per time point. None of the highlighted genes were *GmPGIP*s or *GmPG*s (Alkharouf et al. 2006).

Tucker et al. (2007) infected the *H. glycines*-susceptible *G. max*_[Williams 82/PI 518671]_ with *H. glycines*_[race 3/NL1-RHp/HG-type 7]_ and nematode-colonized root pieces had their RNA isolated at 2, 4, 8, 12, and 20 dpi for hybridization to the Affymetrix soybean GeneChip that contains 37,744 *G. max* probe sets (Tucker et al. 2007). *GmPGIP*s were not highlighted although *G. max PG1* (AF128266), *PG2* (AF128267), *PG3* (AW101688), *PG4*, *PG5*, *PG6* (CA819645)*, PG7*, *PG9*, *PG10* (CD392205), *PG11*, *PG12*, *PG15*, *PG16* (AW734461), and *PG17* (CD409001), were studied. The analysis demonstrated that *PG11* RTAs increased with *H. glycines* infection during a susceptible reaction (Tucker et al. 2007). This observation is consistent with the cell wall loosening that would likely accommodate the expansion of the developing syncytium during a susceptible reaction.

Klink et al. (2007a) performed a time-course comparative microarray analysis of an incompatible (leading to a resistant reaction) and compatible (leading to a susceptible reaction) response by *G. max* to *H. glycines* infection. In those analyses, *H.* glycines_[NL1-RHg/HG-type117]_ (incompatible) and TN8 (compatible) were used to infect *G. max*_[Peking/ PI 548402]_ and the interaction was allowed to proceed with whole root samples collected at 12 hpi, 3 and 8 dpi (Klink et al. 2007). Affymetrix Soybean GeneChip microarrays were used to study gene expression occurring within the whole root (Klink et al. 2007). In those studies, *PGIP1* (*CF806249*) was identified to exhibit an increased RTA in the incompatible interaction, at the 12 and 3 dpi time points and suppressed at the 8 dpi time point. The identified *PGIP1* identified in Klink et al. (2007), when blasting a conceptually translated *CF806249* is most closely (100% identical) related over a stretch of 88 amino acids as *Gm*PGIP7, presented here. No other *G. max* PGIP was greater than 77% identical. The analysis identified the increased RTA of *GmPG021* during an incompatible reaction at 12 hpi, 3, and 8 dpi, respectfully. The analysis also identified an increase in *GmPG021* RTA during the compatible reaction at 12 hpi, 3, and 8 dpi, respectfully. This observation is in agreement with the increased expression of *PG*s accommodating the expansion of the developing syncytium during a susceptible reaction as observed in Tucker et al. (2007) found in *G. max*_[Williams 82/PI 518671_. Therefore, simultaneous expression of *GmPGIP*s and *GmPG*s occur during the resistant reaction, but not all of them exhibit increased transcriptional activity as compared to the control.

Ithal et al. (2007), in a related study that employed the Affymetrix Soybean GeneChip microarrays, using root tissues excised from *G. max*_[Williams 82/PI 518671]_ infected with the inbred population *H. glycines* _[PA3/HG-type 0/race 3]_ identified 429 soybean genes that showed statistically significant differential expression occurring between uninfected and nematode-infected root tissues. The analysis identified *PGIP1* (*GmPGIP7*) to be induced during the compatible interaction at 5 dpi (Ithal et al. 2007). These results are interesting when comparing these observations to Klink et al. (2007). As Klink et al. (2007) demonstrated, increased *GmPGIP7* RTAs at the 12 and 3 dpi time points and decreased RTAs at the 8 dpi time point during an incompatible reaction. *H. glycines.* PGs were not described in Ithal et al. (2007).

Puthoff et al. (2007) infected 14 day old *G. max* _[Williams 82/PI 518671]_ with *H. glycines*_[race 3/NL1-RHp/HG-type 7]_. *H. glycines*-colonized root pieces had their RNA isolated at 8, 12, and 16 dpi for hybridization to the Affymetrix Soybean GeneChip. Of the thousands of identified genes that exhibited differential expression, *PGIP* (Gma.8542.1.S1_at [*Glyma08g08380*]) was identified to have decreased RTAs at 8, 12, and 16 dpi in the *H. glycines*-infected roots as compared to the control (Puthoff et al. 2007). Gma.8542.1.S1_at is not annotated as a *GmPGIP* in the analysis presented here. The *PGIP* GmaAffx.90524.1.S1_s_at was identified to have decreased RTAs at 8, 12, and 16 dpi in the *H. glycines*-infected roots as compared to the control (Puthoff et al. 2007). GmaAffx.90524.1.S1_s_at is not annotated as a *GmPGIP* here. There were several *GmPG*s identified including those with increased RTAs including the BURP domain-containing protein/polygalacturonase, putative (Gma.8368.1.S1_at, GmaAffx.29105.1.S1_at, GmaAffx.91025.1.S1_x_at, Gma.14822.1.S1_at, GmaAffx.60353.1.S1_at), five glycoside hydrolase family 28 protein/polygalacturonase (pectinase) family protein (Gma.15979.1.A1_at, Gma.15979.2.S1_at [*GmPG102*], GmaAffx.37897.1.A1_at, GmaAffx.90527.1.S1_at [*GmPG039*], Gma.3308.1.S1_at [*GmPG028*]), and one suppressed glycoside hydrolase family 28 protein/polygalacturonase (pectinase) family protein (GmaAffx.87008.1.S1_at [*GmPG056*]). The increased expression of *PG*s is consistent with their purported cell wall loosening role(s). The decreased expression on some PGs may indicate a different role or that different PGs function differently or successively during a susceptible reaction.

Afzal et al. (2009) examined the nematode resistance allele at the *rhg1* locus, showing it alters the proteome and primary metabolism of soybean roots during *H. glycines* parasitism. In those studies, the analysis used 14 day old *G. max* near isogenic line (NIL) 34-23 (resistant haplotype between markers Satt214 and Satt570, *G. max*_[NIL 34-23-R]_) and NIL 34-3 (susceptible haplotype between Satt214 and Satt570, *G. max*_[NIL 34-3-S]_) at the F5:13 generation infected with *H. glycines* _[Race 3/HG type 0]_. The root protein was harvested 10 days after infection (dai). The proteomic analysis was able to reproducibly resolve more than 1,000 protein spots on each gel. Neither PGIPs or PGs were identified or mentioned.

Tucker et al. (2011) further utilized gene expression profiling and also performed shared promoter motif analyses for cell wall-modifying proteins expressed in *H. glycines*-infected roots. In those studies, 14 day old SCN-susceptible *G. max* _[Williams 82/PI 518671]_ was infected with *H. glycines*_[race 3/NL1-RHp/HG-type 7]_. Nematode-colonized root pieces had their RNA isolated at 8, 12, and 16 dpi for hybridization to the Affymetrix Soybean GeneChip. The analysis identified *PG1*, *PG2*, *PG3a/b*, *PG4a/b*, *PG5*, *PG6a*, *PG7a/b, PG9a/b*, *PG10a/b*, *PG11a*, *PG12a/b*, *PG15a*, *PG16*, *PG17a*. Analyses showed GUS-*PG11a* promoter fusion produced staining at the ends of the expanding syncytium during the susceptible reaction. This observation is consistent with the role of PGs in macerating cell walls which in this case would be hypothesized to permit expansion of the growing syncytium occurring during a compatible interaction. A conserved WGCATGTG promoter motif, referred to as an SCN-box in the sequence 51 to the start of translation of all the genes for cell wall-modifying proteins was identified and used in primers that PCR amplified *PG3a*, *PG3b*, *PG7b*, *PG11a*, *PG12b*, and *PG16a*. The results indicate that *Gm*PGIPs could be important to deactivate plant and or putative nematode PGs to evoke a resistant reaction. Analyses of whole roots then then shifted to the use of RNA-seq which could provide a comprehensive snapshot of gene RTAs.

Wan et al. (2015) performed RNA-seq analyses of one *H. glycines* -susceptible soybean cultivar, *G. max* _[Magellan]_, and two *H. glycines* -resistant soybean plant introductions (PIs) that included *G. max* _[PI 437654]_ and *G. max* _[PI 567516C],_. The analysis involved infecting them with *H. glycines*_[HG type 0/PA 3]_ (Wan et al. 2015). Both *H. glycines*-inoculated and mock-inoculated root samples were harvested at 0, 3, and 8 dpi. The analysis identified a single *PG* (*GLYMA09G05610*) that was differentially expressed in the at the 0 and 3 dpi sample in the susceptible *G. max* _[Magellan]_. While this observation is consistent with the role of PG function during a compatible interaction, the gene is not annotated as a PG in Wang et al. (2016) and would require further functional analysis to understand its function.

Li et al. (2018) analyzed the *H. glycines*-resistant *G. max* line known as Huipizhi Heidou, ZDD2315 (*G. max* _[Huipizhi Heidou/ZDD2315]_, and an *H. glycines* -susceptible cultivar known as Liaodou15 (*G. max*_[Liaodou15]_). The two lines were infected with *H. glycines*_[race 3]._ and harvested at 5, 10, and 15 dpi. However, the susceptible reaction was not investigated further. In the 5 dpi resistant analysis, Li et al. (2018) identified 2 *PGIP*s, *GLYMA_08G079100* (*GmPGIP7*), and *GLYMA_08G079200* (*GmPGIP8*) to have an increase in their RTAs. Additionally, there was one *GmPG* (*GLYMA_02G037300* [*GmPG008*]) identified to have a decrease in its RTA in the resistant reaction. The 10 dpi analysis identified one *GmPGIP*, *GLYMA_19G145200* (*GmPGIP11*). Furthermore, 3 *GmPG*s with increased RTAs were identified, including, *GLYMA_13G364700* (*GmPG072*), *GLYMA_15G008900* (*GmPG079*), and *GLYMA_15G275400* (*GmPG090*). Two *GmPGIP*s with increased RTAs were identified at the 15 dpi time point including *GLYMA_08G079200* (*GmPGIP8*) and *GLYMA_08G079100* (*GmPGIP7*) which were identified at the 5 dpi time point. A number of *GmPG*s were identified at the 15 dpi time point, with 2 having increased RTAs, including *GLYMA_09G224800* (*GmPG052*) and *GLYMA_16G033000* (*GmPG091*), and 9 *GmPG*s with decreased RTAs including *GLYMA_02G034700*, *GLYMA_18G116400* (*GmPG098*), *GLYMA_03G224600* (*GmPG021*), *GLYMA_15G146800* (*GmPG083*), *GLYMA_19G221600* (*GmPG108*), *GLYMA_14G032900* (*GmPG074*), *GLYMA_13G364700* (*GmPG072*), *GLYMA_19G006200* (*GmPG103*), and *GLYMA_05G005800* (*GmPG026*). *GLYMA_02G034700* is not annotated as a *GmPG* by Wang.

Song et al. (2019) used *G. max*_[Lee/PI 548656]_ which is susceptible to their *H. glycines* population SCN_S_ (*H. glycines*_[SCNS]_) but resistant to SCN_T_ for their use in infection of mock and infected roots occurring for 3 days. At 3 dpi, the authors did not identify any *GmPGIP*s or *GmPG*s among their 1,444 genes with increased RTAs or 246 genes with decreased RTAs during the resistant reaction or 2,320 genes with increased RTAs or 356 genes with decreased RTAs during the susceptible reaction.

Neupane et al. (2019a) transcriptionally profiled the interaction of *G. max* in relation to infection by *H. glycines* and soybean aphids on soybean, sampled at 5 and 30 dpi. The *H. glycines* _[race 3/HG Type 0]_ and *Aphis* glycines (soybean aphid)-susceptible G*. max*_[Williams 82/PI 518671]_ and *H. glycines*_[race 3/HG Type 0]_-resistant but *A*. glycines-susceptible *G. max*_[MN1806CN]_ were infected and RNA samples were collected at 5 and 30 dpi. Of the top 6,000 differentially expressed genes presented, 17 *GmPG*s were identified, including increased RTAs for *GmPG005*, *GmPG008*, *GmPG026*, *GmPG027*, *GmPG034*, *GmPG039*, *GmPG043*, *GmPG046*, *GmPG052*, *GmPG068*, *GmPG072*, *GmPG079*, *GmPG090*, *GmPG102*, *GmPG103*, *GmPG109*, and *GmPG112* at the 5 and 30 dpi time points during both the susceptible and resistant reactions (Neupane et al. 2019a). There were 9 *GmPGIP*s that exhibit differential expression, including the induced *GmPGIP1*, *GmPGIP3*, *GmPGIP4*, *GmPGIP5*, *GmPGIP6*, *GmPGIP7*, *GmPGIP8*, *GmPGIP10*, and *GmPGIP11* during the susceptible and resistant reactions.

Guo et al. (2020) employed the *H. glycines* -susceptible *G. max* _[Magellan]_, and *H. glycines* resistant *G. max*_[Pingliang]_ to be infected with *H. glycines*_[race 3/ HG type 7]_ or to be left uninfected for 10 days prior to the extraction of the total RNA. Of the total, 5,799 and 5,976 significant DEGs were detected between the infected and uninfected roots in the resistant *G. max* _[Pingliang]_ and susceptible *G. max* _[Magellan]_ strains, respectively. In *G. max*_[Pingliang]_, 2,591 and 3,208 up- and downregulated genes were detected, respectively (Guo et al. 2020). In *G. max*_[Magellan]_, 2,596 and 3,380 up- and downregulated genes were detected. *GmPGIP*s and *GmPG*s were not described.

Miraeiz et al. (2020) analyzed *G. max* _[Williams 82/PI 518671]_, *G. max* _[Peking/PI 548402]_, *G. max*_[Fayette/PI 518674]_, and *G. soja*_[PI 468916]._ Roots were infected with *H. glycines*_[race 3/HG type 0]_ for 8 hpi. Miraeiz et al (2020) determined a total of 50 genes in *G. max*_[Williams 82/PI 518671]_, 93 genes in *G. max*_[Fayette/PI 518674]_, and 174 genes in *G. max*_[Peking/PI 548402],_ and 364 genes in *G. soja* _[PI 468916]_ were significantly differentially expressed. Miraeiz et al (2020) identified the induced expression of *GmPG026* at 8 hpi in *G. max* _[Peking/PI 548402]_ demonstrating that induced *PG* expression can occur early during infection, prior to the initiation of the formation of the syncytium in a SCN-resistant genotype where it was not observed in a SCN-susceptible genotype.

Shi et al. (2021) combined targeted metabolite analyses and transcriptomics to reveal the specific chemical composition and associated genes in the *H. glycines*_[HG1.2.3.5.7]-_incompatible *G. max* _[PI 437654]_ and the 3 selected *H. glycines* _[HG1.2.3.5.7]-_ compatible genotypes including *G. max* _[Williams 82/PI 518671_, *G. max* _[Zhonghuang 13]_ and *G. max*_[Hefeng 47]_ (*G. max*_[WM82]_*, G. max*_[ZH13]_, and *G. max*_[HF47]_, respectfully) at 8 dpi. While the comparative transcriptome analyses identified 15,835 DEGs (6,922 increased RTA vs 8,913 decreased RTA), 12,225 DEGs (6,001 increased RTA vs 6,224 decreased RTA), 18,362 DEGs (9,589 increased RTA vs 8,773 decreased RTA) and 19,528 DEGs (8,944 increased RTA vs 10,584 decreased RTA) in the incompatible soybean variety *G. max*_[PI 437654]_, and the three compatible soybean varieties, *G. max* _[WM82]_*, G. max* _[ZH13]_, and *G. max*_[HF47]_, infected by *H. glycines* _[HG1.2.3.5.7]_, respectively. Neither *GmPGIP*s nor *GmPG*s were identified in the transcriptomic study of Shi et al. (2021).

Kofsky et al. (2021) investigated novel resistance strategies to *H. glycines* by examining infection in *G. max*_[Peking/PI 548402]_ and *G. max*_[Williams 82/PI 518671]_ as well as the *H. glycines*-resistant *G. soja* _[PI 468916]_ “NRS100” and *H. glycines* -susceptible *G. soja* _[PI 468916]_ “S-soja” infected with *H. glycines* _[race 5/HG 2.5.7]_. Hundreds of genes were identified. The analysis identified the differential expression of 8 *GmPG*s, including *GmPG005*, *GmPG033*, *GmPG040*, *GmPG043*, *GmPG046*, *GmPG054*, *GmPG055*, and *GmPG101*. No differentially expressed *GmPGIP*s were identified.

Chu et al. (2022) performed an analysis of the *H. glycines*-resistant *G. max*_[Peking_, along with other SCN resistant lines that have a Peking-like response, including *G. max*_[PI 90763]_, *G. max* _[PI 404166]_, *G. max* _[PI 89772]_, *G. max* _[PI 437654]_, and *G. max* _[PI 438489B]_. A second analysis was done on *G. max* _[PI 88788]_ and other PI 88788-like genotypes including *G. max*_[PI 437655]_, *G. max* _[PI 495017C]_, *G. max* _[PI 209332]_, *G. max* _[PI 438503 A]_, and *G. max* _[PI 467312]_. Results from these analyses were compared to the *H. glycines* -susceptible *G. max*_[Hutcheson/PI 518664]_ and genotypes with a similar susceptible response including *G. max*_[Williams 82/PI 518671]_, and *G. max*_[Magellan/PI 595362]_. Chu et al. (2022) identified suppressed expression of *GmPGIP11* in the pooled 3-21 dpi pooled time point sample. This observation is consistent with the suppressed expression by 9 dpi presented here which may indicate its function is not needed after recovery from the resistant reaction.

Torabi et al. (2023) conducted a dual RNA-seq analysis on SCN-resistant *G. max*_[PI 437654]_, *G. max*_[Peking/PI 548402]_, and *G. max*_[PI 88788]_ as well as a susceptible line *G. max*_[Lee 74]_ under exposure to *H. glycines* _[HG1.2.3.5.7]_ to identify specific SCN-responsive genes in soybean and host-specific pathogenesis genes in *H. glycines.* The 5 and 10 dpi time points were chosen as the optimal time points based on a previous study. The analysis at 5 dpi identified that 167, 1,966, 581, and 161 genes (DEGs) were upregulated *G. max*_[PI 437654],_ *G. max* _[Peking/PI 548402],_ and *G. max* _[PI 88788]_ as well as a susceptible line *G. max*_[Lee 74]_, respectively, at 5 dpi (Torabi et al. 2023). The analysis went on to identify 145, 123, 361, and 312 *G. max*_[PI 437654],_ *G. max*_[Peking/PI 548402],_ and *G. max*_[PI 88788]_ as well as a susceptible line *G. max* _[Lee 74]_, at 10 dpi. The differential expression of *GmPGIP*s and *GmPG*s was not observed (Torabi et al. 2023).

For the cell-type-specific syncytium studies, Klink et al. (2005) generated a cDNA library that was produced from RNA extracted from laser microdissected (LM) syncytia developing in *H. glycines* _[race 3/NL1-RHp]_-infected *G. max* _[Kent/PI 548586]_ roots 8 dpi. In the analysis, approximately 800 cDNA clones were one-pass sequenced and the resulting expressed sequence tags (ESTs) were assembled into 174 tentative consensus unigene sequences. *G. max PGIP*s or *PG*s were not among the highlighted genes (Klink et al. 2005).

Klink et al. (2007b) employed the *G. max* _[Peking/PI 548402]_-incompatible *H. glycines*_[NL1-RHg/HG-type117]_ and *G. max* _[Peking/ PI 548402]_-compatible *H. glycines*_[race 14/TN8/HG-type 1.3.6.7]_ to infect *G. max* _[Peking/PI 548402]_. The interaction was allowed to proceed with syncytial samples collected at 3 and 8 dpi (Klink et al. 2007). Affymetrix Soybean GeneChip microarrays were used to study gene expression occurring within the syncytial cell, comparing incompatible and compatible interactions (Klink et al. 2007). The analysis demonstrated that there is little overlap in gene expression signatures between the whole root and syncytium RNA samples in both the incompatible and compatible interaction studies (Klink et al. 2007). This analysis was proof of concept that vastly different sets of genes, possibly important defense or susceptibility genes, were differentially expressed within the syncytium as compared to the whole root that may otherwise escape detection. The analysis went on to identify genes expressed specifically in the incompatible syncytium as compared to the compatible syncytium (Klink et al. 2007). Neither *PGIP*s or *PG*s were highlighted among the induced or suppressed genes (Klink et al. 2007b).

Ithal et al. (2007) studied the 2, 5, and 10 dpi time points. Ithal et al. (2007) employed the *H. glycines* _[race 3/PA3/HG-type 0]_-compatible *G. max*_[Williams 82/PI 518671]_. Neither *PGIP*s or *PG*s were highlighted in their analyses (Ithal et al. 2007).

Klink et al. (2009) performed a gene expression analysis of syncytia laser microdissected from the roots of the *G. max* _[Peking/PI 548402]_ undergoing a resistant reaction after infection by *H. glycines*_[NL1-RHg/HG-type117]_. In the *G. max* _[Peking/PI 548402]_ 6 dpi resistant syncytium as compared to its pericycle cells Gma.8542.1.S1_at measured that the PGIP precursor (AI437934) exhibited induced gene expression (Klink et al. 2009). AI437934 became *Glyma08g08380* in the original Wm82.a1.v1.1 genome annotation but subsequently was eliminated from the latter annotations Wm82.a2.v1 and Wm82.a4.v1 so no *GmPGIP* identifier is given and will not be presented further.

Klink et al. (2010a) Then examined a different form of the resistant reaction that occurs in *G. max*_[PI 88788]_. *G. max*_[PI 88788]_ has resistance loci that are both common to and different from *G. max* _[Peking’PI 540402]_. Klink et al. (2010b) infected *G. max* _[PI 88788]_ with either *H. glycines*_[NL1-RHg/HG-type117]_ and *H. glycines*_[TN8/HG-type111.3.6.7]_. The analysis identified induced *GmPG026*, and *GmPG101* at the 3, 6, and 9 dpi time points during the resistant reaction. During the susceptible reaction, the analysis identified induced expression of *GmPG098* at the 3 dpi time point and induced *GmPG101* at the 3, 6, and 9 dpi time points. The analysis also identified suppressed *GmPG026* at the 3 dpi time point during the susceptible reaction. It is unclear why *GmPG101* is induced constitutively during the resistant and susceptible reaction. In contrast, *GmPG026* is induced at the 3 dpi time point when susceptible and resistant reactions appear similar, cytologically, but then is not differentially expressed at the 6 or 9 dpi time point during the susceptible reaction. *GmPG098* is expressed at the 3 dpi time point during the susceptible reaction like *GmPG026* but then no longer experiences differential expression. This characteristic may indicate an important early role for *GmPG098* and *GmPG026* but only during the susceptible reaction. Perhaps the soybean defense strategy that employs PGIPs is circumvented by the nematode activating different host *PG*s during parasitism. The question then became whether gene expression at the syncytium was different or similar between soybean genotypes with different forms of the resistant reaction.

Klink et al. (2010b) then focused on identifying genes expressed only within the parasitized root cells undergoing either an incompatible or compatible interaction. These are the genes that are normally discarded in a differential expression analysis since a statistical determination of differential expression to a control where expression must occur for the comparison to be made is not possible. A single *G. max* genotype (*G. max*_[PI11548402/Peking]_) was used in the experiments to obtain both incompatible and compatible reactions by the use of two different populations of *H. glycines, H. glycines*_[NL1-RHg/HG-type117]_ and *H. glycines*_[TN8/HG-type111.3.6.7]_. Hundreds of genes were identified in microarray comparisons of genes found only in the compatible or incompatible syncytium where they were not expressed in the control cells (syncytium) with the focus of the study being on select genes annotated as belonging to Disease and Defense, Signaling, and Transcription categories. Neither *PGIP*s nor *PG*s were highlighted in the analyses. However, a more in depth analysis was presented in a later study.

Klink et al. (2011) by comparing two different forms of the resistant reaction identified differences in gene expression amplitude overlie a conserved transcriptomic program occurring between the rapid and potent localized resistant reaction at the syncytium of *G. max* _[Peking/PI 548402]_ as compared to the prolonged and potent resistant reaction of *G. max* _[PI 88788]_. There were a number of *PG*s that were found to be differentially expressed. A comparison of the 3 dpi syncytium to the control identified induced expression of Gma.8494.1.S1_at (*GmPG101*), and Gma.8493.1.S1_at (*GmPG026*), while Gma.7587.1.S1_at (*GmPG098*) was suppressed. PGs were found to be constitutively induced throughout the time course analysis at 3, 6 and 9 dpi, including Gma.8494.1.S1_at (*GmPG101*), and Gma.8493.1.S1_at (*GmPG026*). A pathway analysis of 9 dpi to the 6 dpi resistant syncytium identified induced *PG*s including Gma.6599.1.S1_at (*GmPG049*) (Klink et al. 2011). Klink et al. (2011) then performed cross-comparative analyses between the previously studied 0, 3, 6 and 9 dpi time points of *G. max* _[Peking/PI 548402]_ and *G. max* _[PI 88788]_. In the first analysis of the 0 dpi pericycle control, no *PGIP*s were found to be expressed at a higher level in *G. max*_[Peking/PI 548402]_ as compared to *G. max* _[PI 88788]_. However, 4 *PG*s were found to be expressed at a higher level in *G. max* _[Peking/PI 548402]_ as compared to *G. max* _[PI 88788]_. Gma.15979.1.A1_at (*CD414564*), Gma.8494.1.S1_at (*AF128266*), GmaAffx.92741.1.S1_s_at (*CF808466*), and Gma.3308.1.S1_at *(CD414773*). An expression analyses, comparing *G. max* _[Peking/PI 548402]_ syncytium gene expression directly to *G. max*_[PI 88788]_ syncytium gene expression at 3 dpi, identified no statistically significand differences in gene expression. An expression analyses comparing *G. max*_[Peking/PI 548402]_ syncytium gene expression directly to *G. max*_[PI 88788]_ syncytium gene expression at 6 dpi identified suppressed expression of 3 *PGIP*s including Gma.8542.1.S1_at (*AI437934*), GmaAffx.90524.1.S1_s_at (*CF806249*), Gma.9637.1.S1_at (*AF529302*). The same analyses identified suppressed expression of two *PG*s including GmaAffx.92774.1.S1_at (*CF808124*), GmaAffx.37897.1.A1_at (*BU545459*). Furthermore, the analysis of the 3 dpi *G. max* _[Peking/PI 548402 + PI 88788]_ syncytium sample identified induced expression of the *PGIP* Gma.8542.1.S1_at (*AI437934*) as compared to the pericycle. The same analysis identified induced expression of 7 PGs including GmaAffx.60035.1.S1_at (*BM271340*), Gma.8494.1.S1_at (*AF128266*), Gma.8493.1.S1_at (*AF128267*), Gma.14822.1.S1_at (*AW348778*), GmaAffx.92774.1.S1_at (*CF808124*), GmaAffx.47891.1.S1_s_at (*BQ785880*), GmaAffx.87008.1.S1_at (*BE804610*), and GmaAffx.37897.1.A1_at (*BU545459*). The analysis of the 6 dpi *G. max* _[Peking/PI 548402 + PI 88788]_ syncytium sample identified induced expression of the same Gma.8542.1.S1_at (*AI437934*) as compared to the pericycle. The same analysis identified induced expression of 6 of the same 7 same *PG*s including GmaAffx.60035.1.S1_at (*BM271340*), Gma.8494.1.S1_at (*AF128266*), Gma.8493.1.S1_at (*AF128267*), GmaAffx.92774.1.S1_at (*CF808124*), GmaAffx.47891.1.S1_s_at (*BQ785880*), and GmaAffx.37897.1.A1_at (*BU545459*) with GmaAffx.87008.1.S1_at (*BE804610*) dropping out. The analysis of the 9 dpi *G. max*_[Peking/PI 548402 + PI 88788]_ syncytium sample identified induced expression of the same Gma.8542.1.S1_at (*AI437934*) as compared to the pericycle. The same analysis identified three *PG*s were still induced, including Gma.8494.1.S1_at (*AF128266*), Gma.8493.1.S1_at (*AF128267*), and GmaAffx.37897.1.A1_at (*BU545459*). A final analysis representing the time course combined samples identified probe sets that measure induced or suppressed levels of gene expression across all time points (3, 6 and 9 dpi *G. max* _[Peking/PI 548402 + PI 88788]_) as compared to the pericycle *G. max* _[Peking/PI 548402 + PI 88788]_. As expected, the analysis identified the induced *PGIP* Gma.8542.1.S1_at (*AI437934*) and induced *PG*s Gma.8494.1.S1_at (*AF128266*), Gma.8493.1.S1_at (*AF128267*), and GmaAffx.37897.1.A1_at (*BU545459*).

Matsye et al. (2011) mapped cell fate decisions that occur during soybean defense responses and, importantly, provided the first cell-type specific RNA-seq analysis of the syncytium undergoing a defense response. Matsye et al. (2011) identified *GmPG056* as being induced during the susceptible reaction at 3 and 9 dpi. *GmPG102* is induced during the susceptible reaction at the 3, 6, and 9 dpi time points. Matsye identified *GmPG026* as expressed at the 9 dpi time point during a resistant reaction in *G. max*_[PI 88788]_. Matsye identified *GmPG033* as expressed at the 9 dpi time point during a resistant reaction in *G. max* _[Peking/PI 548402]_. Matsye identified *GmPG061* as expressed during the resistant reaction at the 3 dpi time point in *G. max*_[Peking/PI 548402]_ and the 9 dpi time point in *G. max* _[PI 88788]_. Matsye identified *GmPG080* as expressed during the resistant reaction at the 6 dpi time point in *G. max* _[Peking/PI 548402]_. Matsye identified *GmPG105* as expressed during the resistant reaction at the 3 and 9 dpi time points in *G. max*_[PI 88788]_. Matsye identified *GmPG033* as expressed during the resistant reaction at the 3 and 6 dpi time points in *G. max*_[Peking/PI 548402]_ and *G. max*_[PI 88788]_. Matsye identified *GmPG049* as expressed during the resistant reaction at the 3 and 6 dpi time points in *G. max*_[Peking/PI 548402]_ and *G. max* _[PI 88788]_. Matsye identified *GmPG061* as expressed during the resistant reaction at the 6 dpi time point in *G. max*_[Peking/PI 548402]_ and *G. max*_[PI 88788]_. Matsye identified *GmPG098* as expressed during the resistant reaction at the 3 and 6 dpi time points in *G. max*_[Peking/PI 548402]_ and *G. max*_[PI 88788]_. Matsye identified *GmPG105* as expressed during the resistant reaction at the 6 dpi time point in *G. max*_[Peking/PI 548402]_ and *G. max* _[PI 88788]_. The two probe sets for *GmPGIP11* measured expression. GmaAffx.47891.1.S1_s_at (*BQ785880*), measured expression during the resistant reaction at the 3 and 6 dpi time points in *G. max*_[Peking/PI 548402]_ and *G. max*_[PI 88788]_ while GmaAffx.92774.1.S1_at (*CF808124*) measured expression at the 3 and 9 dpi time points in *G. max*_[Peking/PI 548402]_ and *G. max*_[PI 88788]_ (Matsye et al. 2011). GmaAffx.92774.1.S1_at measured *GmPGIP11* expression only in *G. max*_[PI 88788]_ at the 6 dpi time point.

Kandoth et al. (2011) examined a *G. max* near isogenic line (NIL) differing at the *Rhg1* locus (NIL-R and NIL-S) that was derived from a cross between the *H. glycines*-resistant PI 209332 and susceptible Evans (*G. max*_[Evans/ PI 548560]_). The NILs were infected with *H. glycines* _[PA3/HG type 0]_ and *H. glycines* _[TN19/HG type 1-7]._ Laser microdissection was performed, and microarray analyses done on the RNA isolated from the syncytia at 5 and 8 dpi with SCN from either the NIL-S or NIL-R. Since there was no significant evidence of interaction between NIL and dpi, the analysis focused on the genes of NIL-S or NIL-R as a whole. This observation was in contrast to those identified by Klink et al. 2007, 2009, 2010a, b, 2011; Matsye et al. 2011). It could be explained that the same *H. glycines* population that would have the same sets of parasitism genes was used to infect the 2 closely related NILs in Kandoth et al. (2011) while Klink et al. (2007b, 2009, 2010a, 2010b, 2011) and Matsye et al. 2011) used the identical soybean genotype but different *H. glycines* populations to achieve their susceptible and resistant reactions. Kandoth et al. (2011) identified 1,447 differentially expressed genes occurring between NIL-S or NIL-R with 828 being up-regulated and 619 being down-regulated. Neither *PGIP*s nor *PG*s were highlighted in those analyses. Three suppressed glycoside hydrolase family 28 protein / polygalacturonase (pectinase) family protein probe sets measured suppressed expression, including GmaAffx.82513.1.S1_at (*Glyma07g07280.1* [*GmPG033*]), GmaAffx.37897.1.A1_at (*Glyma16g03680.1* [*GmPG091*]), and GmaAffx.17518.1.S1_at (*Glyma03g38350.3* [*GmPG021*]).

