## Supplementary figures and images for "*Glycine max* polygalacturonase inhibiting protein 11 (*Gm*PGIP11) functions in the root to suppress *Heterodera glycines* parasitism"

### Supplemental Figure 1

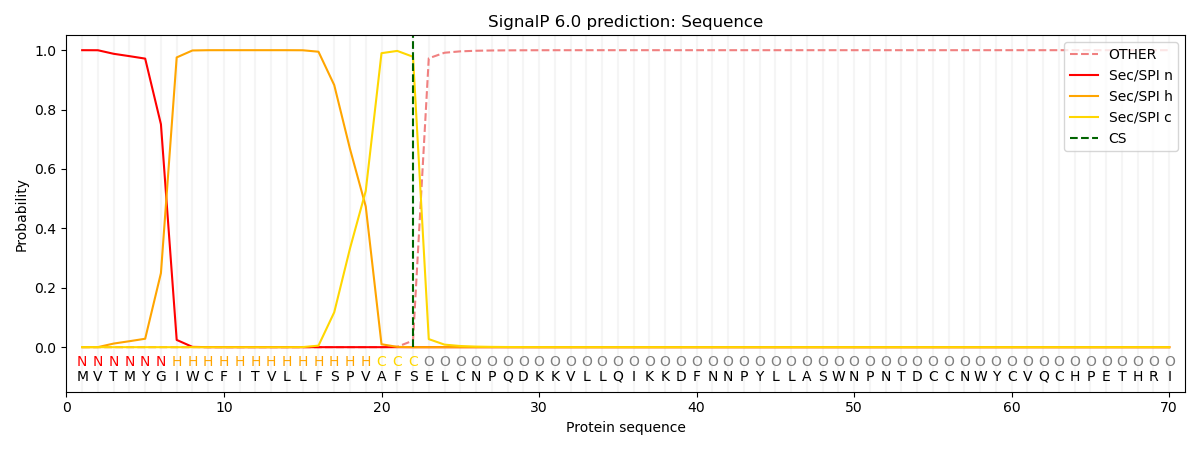

### Supplemental Figure 2

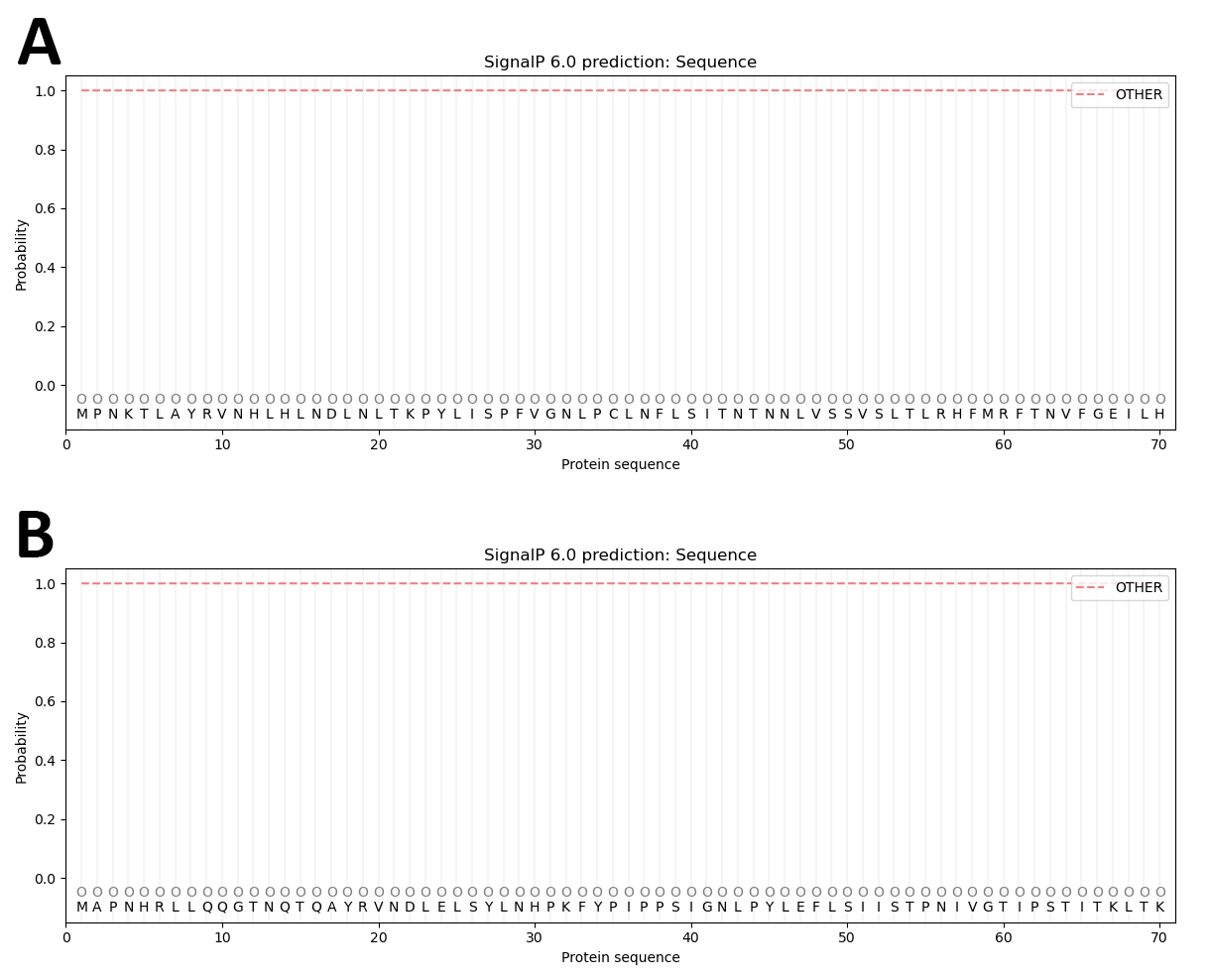

### Supplemental Figure 5

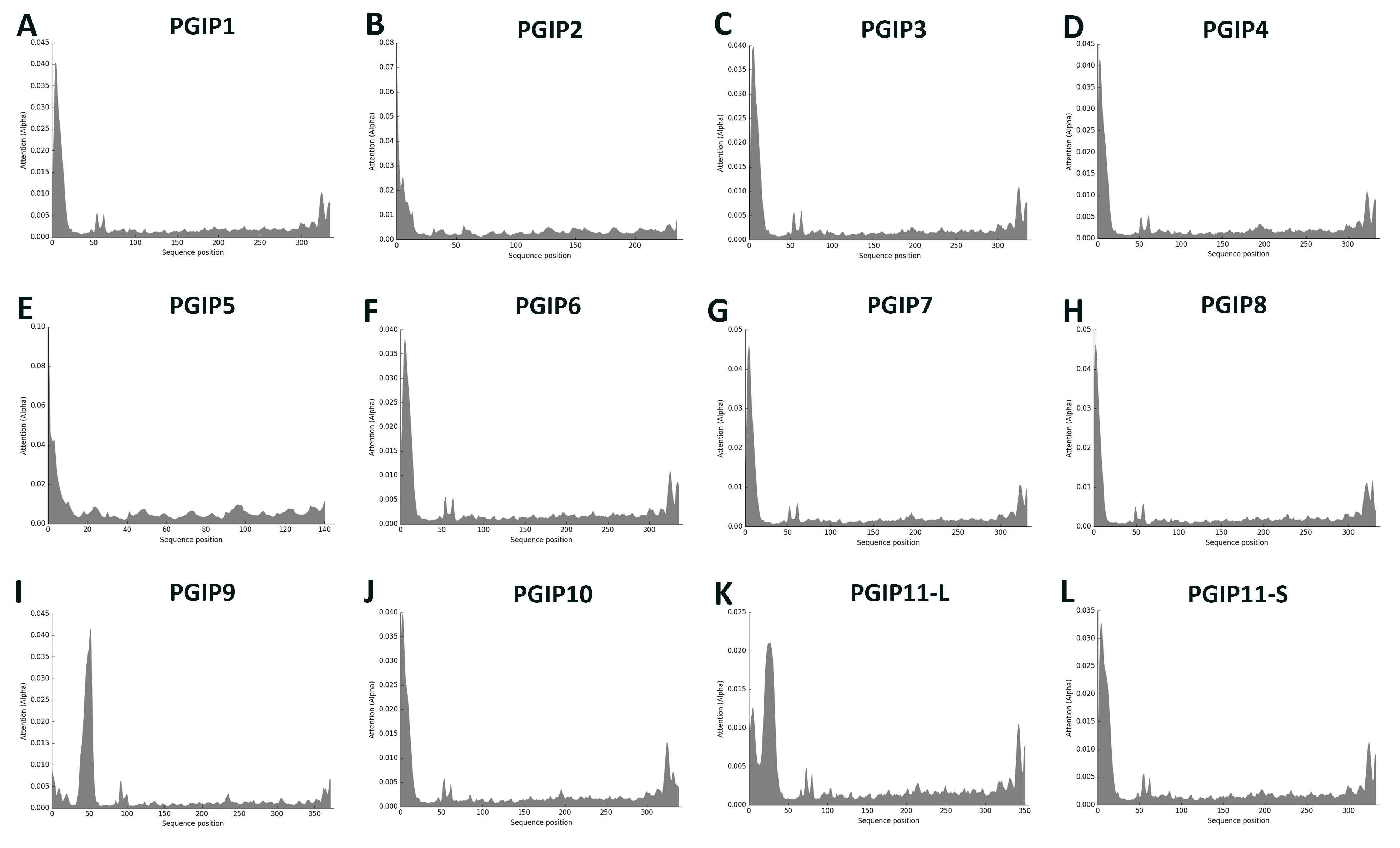

### Supplemental Figure 6

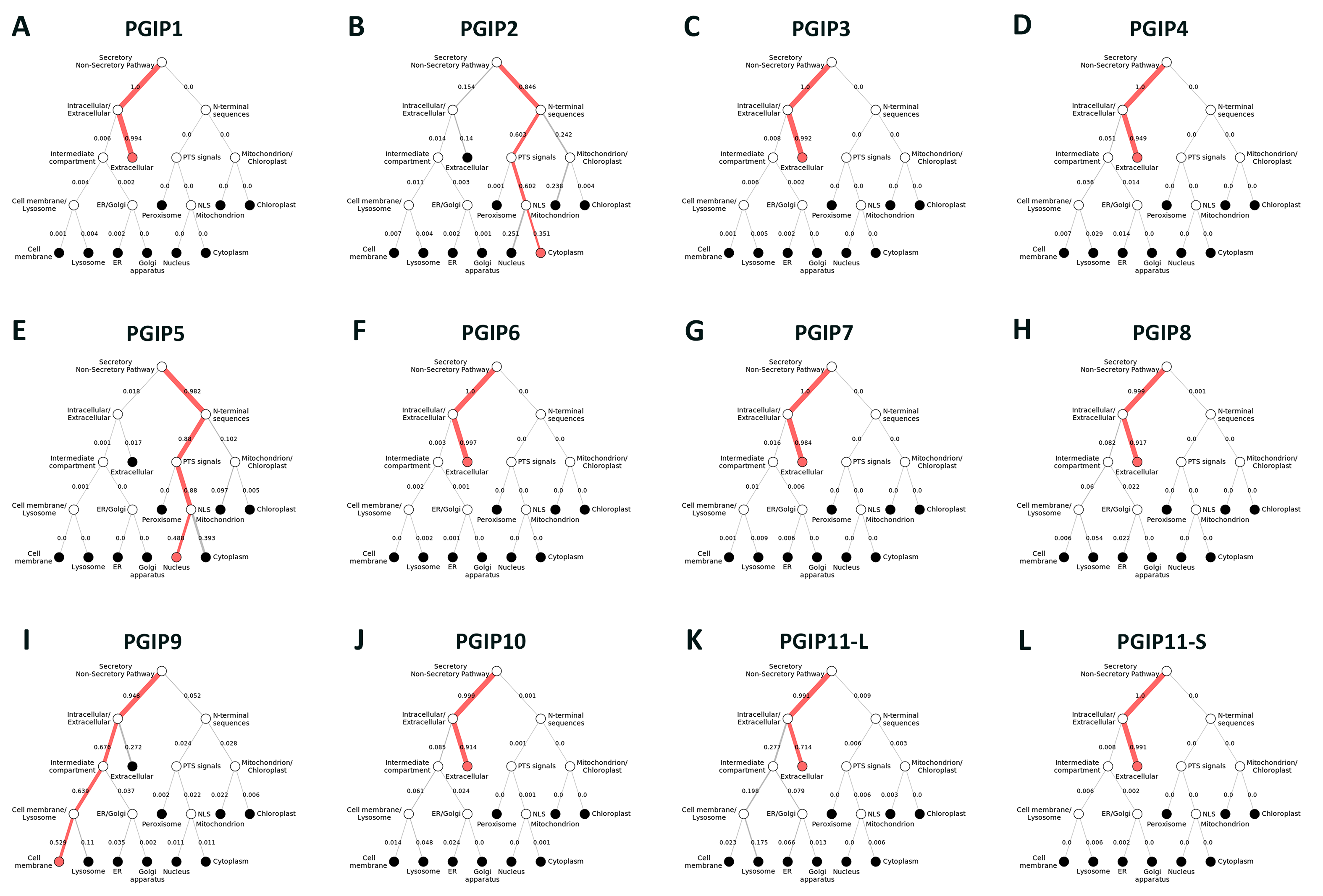

### Supplemental Figure 7

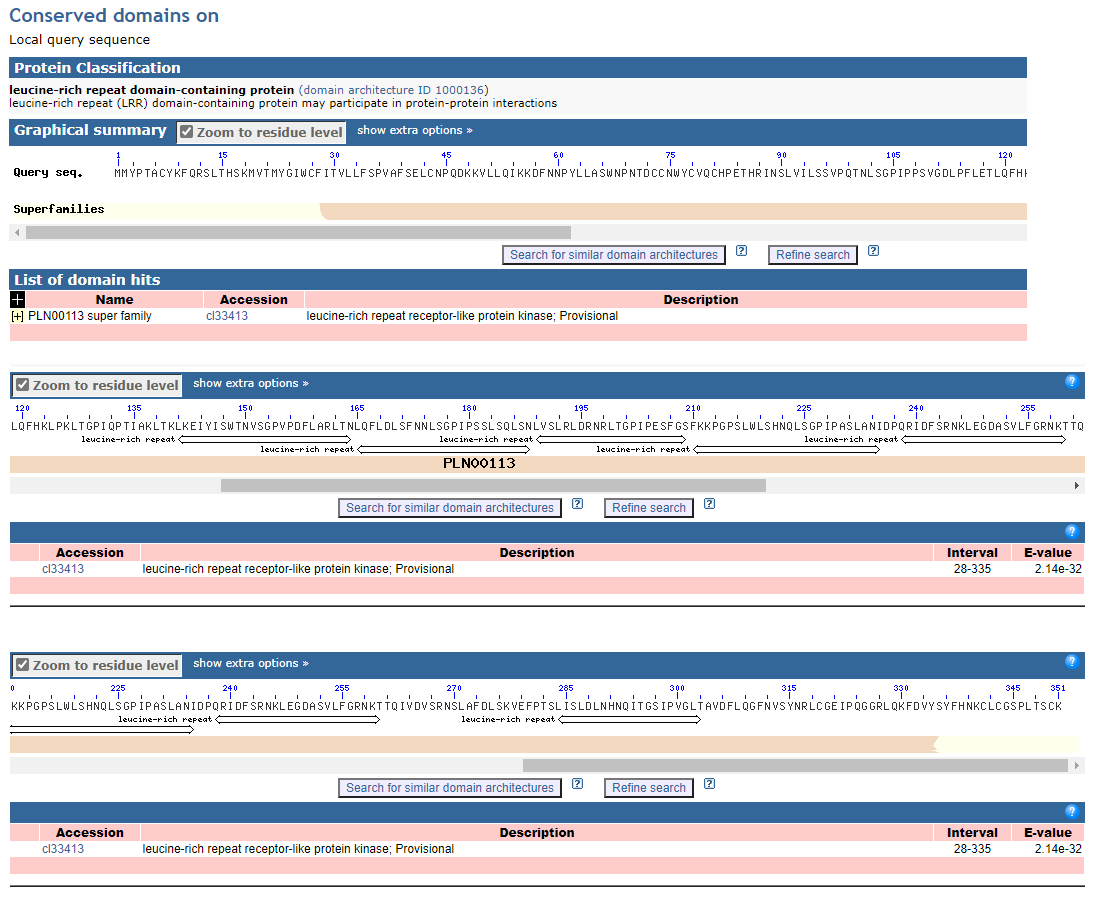
