## Supplemental Figure 3 for "*Glycine max* polygalacturonase inhibiting protein 11 (*Gm*PGIP11) functions in the root to suppress *Heterodera glycines* parasitism"

***Gm*PGIP1**

```
>G.max Wm82.a2.v1|Glyma.05G123700.1.p
(Threshold=0.5)
--------------------------------------------------------------------------------------
      		   		Jury    	N-Glyc
```

**SeqName^a^	Position^b^	Potential^c^	agreement^d^	result^e^**

**--------------------------------------------------------------------------------------**

**GmPGIP1	41	NPTT	0.6289	(8/9)	+	WARNING: PRO-X1.**

**GmPGIP1	132 NVSG	0.7063	(9/9)	++**

**GmPGIP1	250 NNSF	0.4126	(7/9)	-**

^a^The GmPGIP under study.

^b^The position refers to the amino acid position N terminal to C terminal along the protein.

^c^The potential scores shown are the averaged output of nine neural networks.

^d^Jury agreement indicates how many of the nine networks support the prediction.

^e^N-Glyc results are indicated (+) for a potential N-glycosylation site >0.5 threshold, (++) for a potential N-glycosylation site >0.5 threshold and jury agreement of 9/9 and (+++) for a potential N-glycosylation site >0.75 and jury agreement of 9/9. (−−−) indicates a potential N-glycosylation score <0.5 threshold.

***Note: The** WARNING: PRO-X1. **Proline**, when occurring just after the asparagine residue, makes it highly unlikely that the asparagine is glycosylated, presumably due to conformational constraints.

***Gm*PGIP2**

```
>G.max Wm82.a2.v1|Glyma.05G123800.1.p
(Threshold=0.5)
--------------------------------------------------------------------------------------
      		   	Jury    	N-Glyc
```

**SeqName^a^	Position^b^	Potential^c^	agreement^d^	result^e^**

```
--------------------------------------------------------------------------------------
GmPGIP2       3 NKTL   	0.7646     	(9/9)   		+++
GmPGIP2      19 NLTK   	0.7558     	(9/9)   		+++
GmPGIP2      87 NFSG   	0.5789     	(7/9)   		+
GmPGIP2     165 NNSF   	0.2583     	(9/9)   		---
```

^a^The GmPGIP under study.

^b^The position refers to the amino acid position N terminal to C terminal along the protein.

^c^The potential scores shown are the averaged output of nine neural networks.

^d^Jury agreement indicates how many of the nine networks support the prediction.

^e^N-Glyc results are indicated (+) for a potential N-glycosylation site >0.5 threshold, (++) for a potential N-glycosylation site >0.5 threshold and jury agreement of 9/9 and (+++) for a potential N-glycosylation site >0.75 and jury agreement of 9/9. (−−−) indicates a potential N-glycosylation score <0.5 threshold.

***Gm*PGIP3**

```
>G.max Wm82.a2.v1|Glyma.05G123900.1.p
(Threshold=0.5)
--------------------------------------------------------------------------------------
      		   				Jury    	N-Glyc
```

**SeqName^a^	Position^b^	Potential^c^	agreement^d^	result^e^**

```
--------------------------------------------------------------------------------------
GmPGIP3      41 NPTN   	0.6902     	(8/9)   	+	WARNING: PRO-X1.
GmPGIP3      44 NLSS   	0.6370     	(9/9)   		++
```

^a^The GmPGIP under study.

^b^The position refers to the amino acid position N terminal to C terminal along the protein.

^c^The potential scores shown are the averaged output of nine neural networks.

^d^Jury agreement indicates how many of the nine networks support the prediction.

^e^N-Glyc results are indicated (+) for a potential N-glycosylation site >0.5 threshold, (++) for a potential N-glycosylation site >0.5 threshold and jury agreement of 9/9 and (+++) for a potential N-glycosylation site >0.75 and jury agreement of 9/9. (−−−) indicates a potential N-glycosylation score <0.5 threshold.

***Note: The** WARNING: PRO-X1. **Proline**, when occurring just after the asparagine residue, makes it highly unlikely that the asparagine is glycosylated, presumably due to conformational constraints.

***Gm*PGIP4**

```
>G.max Wm82.a2.v1|Glyma.05G124000.1.p
(Threshold=0.5)
--------------------------------------------------------------------------------------
      		   				Jury    	N-Glyc
```

**SeqName^a^	Position^b^	Potential^c^	agreement^d^	result^e^**

```
--------------------------------------------------------------------------------------
GmPGIP4      39 NPTT   	0.6266    	(8/9)   	+    WARNING: PRO-X1.
GmPGIP4      54 NNSW   	0.5838    	(6/9)   	+
GmPGIP4     131 NISG   	0.5928    	(8/9)   	+
```

^a^The GmPGIP under study.

^b^The position refers to the amino acid position N terminal to C terminal along the protein.

^c^The potential scores shown are the averaged output of nine neural networks.

^d^Jury agreement indicates how many of the nine networks support the prediction.

^e^N-Glyc results are indicated (+) for a potential N-glycosylation site >0.5 threshold, (++) for a potential N-glycosylation site >0.5 threshold and jury agreement of 9/9 and (+++) for a potential N-glycosylation site >0.75 and jury agreement of 9/9. (−−−) indicates a potential N-glycosylation score <0.5 threshold.

***Note: The** WARNING: PRO-X1. **Proline**, when occurring just after the asparagine residue, makes it highly unlikely that the asparagine is glycosylated, presumably due to conformational constraints.

***Gm*PGIP5**

```
>G.max Wm82.a2.v1|Glyma.08G078800.1.p
(Threshold=0.5)
--------------------------------------------------------------------------------------
      		   				Jury    	N-Glyc
```

**SeqName^a^	Position^b^	Potential^c^	agreement^d^	result^e^**

```
--------------------------------------------------------------------------------------
GmPGIP5      13 NQTQ   	0.6107    	 (9/9)  		++
GmPGIP5      80 NLSG   	0.5152    	(5/9)  		+
GmPGIP5     120 NFSE   	0.5788    	(7/9)  		+
```

^a^The GmPGIP under study.

^b^The position refers to the amino acid position N terminal to C terminal along the protein.

^c^The potential scores shown are the averaged output of nine neural networks.

^d^Jury agreement indicates how many of the nine networks support the prediction.

^e^N-Glyc results are indicated (+) for a potential N-glycosylation site >0.5 threshold, (++) for a potential N-glycosylation site >0.5 threshold and jury agreement of 9/9 and (+++) for a potential N-glycosylation site >0.75 and jury agreement of 9/9. (−−−) indicates a potential N-glycosylation score <0.5 threshold.

***Gm*PGIP6**

```
>G.max Wm82.a2.v1|Glyma.08G078900.1.p
(Threshold=0.5)
--------------------------------------------------------------------------------------
      		   				Jury    	N-Glyc
```

**SeqName^a^	Position^b^	Potential^c^	agreement^d^	result^e^**

```
--------------------------------------------------------------------------------------
GmPGIP6      41 NPTT   	0.5917    	(7/9)   		+    WARNING: PRO-X1.
GmPGIP6      66 NKTQ   	0.7202     	(9/9)   		++
GmPGIP6     133 NVSG   	0.5514     	(7/9)   		+
GmPGIP6     157 NFSG  	0.5312     	(5/9)   		+
GmPGIP6     251 NNSF   	0.2423     	(9/9)   		---
```

^a^The GmPGIP under study.

^b^The position refers to the amino acid position N terminal to C terminal along the protein.

^c^The potential scores shown are the averaged output of nine neural networks.

^d^Jury agreement indicates how many of the nine networks support the prediction.

^e^N-Glyc results are indicated (+) for a potential N-glycosylation site >0.5 threshold, (++) for a potential N-glycosylation site >0.5 threshold and jury agreement of 9/9 and (+++) for a potential N-glycosylation site >0.75 and jury agreement of 9/9. (−−−) indicates a potential N-glycosylation score <0.5 threshold.

***Note: The** WARNING: PRO-X1. **Proline**, when occurring just after the asparagine residue, makes it highly unlikely that the asparagine is glycosylated, presumably due to conformational constraints.

***Gm*PGIP7**

```
>G.max Wm82.a2.v1|Glyma.08G079100.1.p
(Threshold=0.5)
--------------------------------------------------------------------------------------
      		   				Jury    	N-Glyc
```

**SeqName^a^	Position^b^	Potential^c^	agreement^d^	result^e^**

```
--------------------------------------------------------------------------------------
GmPGIP7      39 NPTT   	0.6136     	(7/9)   	+    WARNING: PRO-X1*.
GmPGIP7      54 NRTW   	0.6153     	(8/9)   	+
GmPGIP7     131 NVSG   	0.5373     	(6/9)   	+
```

**GmPGIP7	293 NVSF	0.4779	(6/9)	-**

^a^The GmPGIP under study.

^b^The position refers to the amino acid position N terminal to C terminal along the protein.

^c^The potential scores shown are the averaged output of nine neural networks.

^d^Jury agreement indicates how many of the nine networks support the prediction.

^e^N-Glyc results are indicated (+) for a potential N-glycosylation site >0.5 threshold, (++) for a potential N-glycosylation site >0.5 threshold and jury agreement of 9/9 and (+++) for a potential N-glycosylation site >0.75 and jury agreement of 9/9. (−−−) indicates a potential N-glycosylation score <0.5 threshold.

*Note: The WARNING: PRO-X1. **Proline**, when occurring just after the asparagine residue, makes it highly unlikely that the asparagine is glycosylated, presumably due to conformational constraints.

***Gm*PGIP8**

```
>G.max Wm82.a2.v1|Glyma.08G079200.1.p
(Threshold=0.5)
--------------------------------------------------------------------------------------
      		   				Jury    	N-Glyc
```

**SeqName^a^	Position^b^	Potential^c^	agreement^d^	result^e^**

```
--------------------------------------------------------------------------------------
GmPGIP8      36 NPTK   	0.7197     	(9/9)   	++   WARNING: PRO-X1.
GmPGIP8      65 NQTC   	0.5721     	(6/9)   	+
GmPGIP8     116 NLTN   	0.5719     	(8/9)   	+
GmPGIP8     124 NITY   	0.7316     	(9/9)   	++
GmPGIP8     129 NVSG   	0.5883     	(7/9)   	+
GmPGIP8     153 NLSG   	0.6071     	(8/9)   	+
```

**GmPGIP8	291 NVSN	0.6118	(9/9)	++**

^a^The GmPGIP under study.

^b^The position refers to the amino acid position N terminal to C terminal along the protein.

^c^The potential scores shown are the averaged output of nine neural networks.

^d^Jury agreement indicates how many of the nine networks support the prediction.

^e^N-Glyc results are indicated (+) for a potential N-glycosylation site >0.5 threshold, (++) for a potential N-glycosylation site >0.5 threshold and jury agreement of 9/9 and (+++) for a potential N-glycosylation site >0.75 and jury agreement of 9/9. (−−−) indicates a potential N-glycosylation score <0.5 threshold.

***Note: The** WARNING: PRO-X1. **Proline**, when occurring just after the asparagine residue, makes it highly unlikely that the asparagine is glycosylated, presumably due to conformational constraints.

***Gm*PGIP9**

```
>G.max Wm82.a2.v1|Glyma.15G209200.1.p
(Threshold=0.5)
--------------------------------------------------------------------------------------
      		   				Jury    	N-Glyc
```

**SeqName^a^	Position^b^	Potential^c^	agreement^d^	result^e^**

```
--------------------------------------------------------------------------------------
GmPGIP9     121 NASG   	0.4621     	(7/9)   	-
GmPGIP9     170 NISG   	0.5773     	(6/9)   	+
GmPGIP9     194 NLSG   	0.5879     	(7/9)   	+
GmPGIP9     333 NVSY   	0.5652     	(6/9)   	+
```

^a^The GmPGIP under study.

^b^The position refers to the amino acid position N terminal to C terminal along the protein.

^c^The potential scores shown are the averaged output of nine neural networks.

^d^Jury agreement indicates how many of the nine networks support the prediction.

^e^N-Glyc results are indicated (+) for a potential N-glycosylation site >0.5 threshold, (++) for a potential N-glycosylation site >0.5 threshold and jury agreement of 9/9 and (+++) for a potential N-glycosylation site >0.75 and jury agreement of 9/9. (−−−) indicates a potential N-glycosylation score <0.5 threshold.

***Gm*PGIP10**

```
>G.max Wm82.a2.v1|Glyma.15G209300.1.p
(Threshold=0.5)
--------------------------------------------------------------------------------------
      		   				Jury    	N-Glyc
```

**SeqName^a^	Position^b^	Potential^c^	agreement^d^	result^e^**

```
--------------------------------------------------------------------------------------
GmPGIP10      82 NLSA   	0.6827     	(7/9)   	+
GmPGIP10     131 NLSG   	0.6373     	(9/9)   	++
GmPGIP10     155 NLSG   	0.5572     	(6/9)   	+
```

^a^The GmPGIP under study.

^b^The position refers to the amino acid position N terminal to C terminal along the protein.

^c^The potential scores shown are the averaged output of nine neural networks.

^d^Jury agreement indicates how many of the nine networks support the prediction.

^e^N-Glyc results are indicated (+) for a potential N-glycosylation site >0.5 threshold, (++) for a potential N-glycosylation site >0.5 threshold and jury agreement of 9/9 and (+++) for a potential N-glycosylation site >0.75 and jury agreement of 9/9. (−−−) indicates a potential N-glycosylation score <0.5 threshold.

***Gm*PGIP11**

```
>G.max Wm82.a2.v1|Glyma.19G145200.1.p
--------------------------------------------------------------------------------------
      		   				Jury    	N-Glyc
```

**SeqName^a^	Position^b^	Potential^c^	agreement^d^	result^e^**

**--------------------------------------------------------------------------------------**

**GmPGIP11	101 NLSG	0.6096	(8/9)	+**

**GmPGIP11	150 NVSG	0.6658	(9/9)	++**

**GmPGIP11	174 NLSG	0.5431	(6/9)	+**

**GmPGIP11	258 NKTT	0.5788	(5/9)	+**

**GmPGIP11	312 NVSY	0.5620	(8/9)	+**

^a^The GmPGIP under study.

^b^The position refers to the amino acid position N terminal to C terminal along the protein.

^c^The potential scores shown are the averaged output of nine neural networks.

^d^Jury agreement indicates how many of the nine networks support the prediction.

^e^N-Glyc results are indicated (+) for a potential N-glycosylation site >0.5 threshold, (++) for a potential N-glycosylation site >0.5 threshold and jury agreement of 9/9 and (+++) for a potential N-glycosylation site >0.75 and jury agreement of 9/9. (−−−) indicates a potential N-glycosylation score <0.5 threshold.
