## Supplemental Figure 4 for "*Glycine max* polygalacturonase inhibiting protein 11 (*Gm*PGIP11) functions in the root to suppress *Heterodera glycines* parasitism"

**(*Gm*PGIP1)**

```
##gff-version 2
##source-version NetOGlyc 4.0.0.13
##date 22-2-18
##Type Protein
```

#seqname^a^	source^b^	feature^c^	start^d^	end^e^	score^f^	strand^g^	frame^h^	comment^i^

```
PGIP1	netOGlyc-4.0.0.13	CARBOHYD	2	2	0.106902	.	.
PGIP1	netOGlyc-4.0.0.13	CARBOHYD	5	5	0.0544594	.	.
PGIP1	netOGlyc-4.0.0.13	CARBOHYD	15	15	0.0473706	.	.
PGIP1	netOGlyc-4.0.0.13	CARBOHYD	17	17	0.110401	.	.
PGIP1	netOGlyc-4.0.0.13	CARBOHYD	18	18	0.0644824	.	.
PGIP1	netOGlyc-4.0.0.13	CARBOHYD	21	21	0.0545614	.	.
PGIP1	netOGlyc-4.0.0.13	CARBOHYD	43	43	0.430489	.	.
PGIP1	netOGlyc-4.0.0.13	CARBOHYD	44	44	0.448902	.	.
PGIP1	netOGlyc-4.0.0.13	CARBOHYD	46	46	0.35457	.	.
PGIP1	netOGlyc-4.0.0.13	CARBOHYD	47	47	0.75321	.	.	#POSITIVE
PGIP1	netOGlyc-4.0.0.13	CARBOHYD	51	51	0.270979	.	.
PGIP1	netOGlyc-4.0.0.13	CARBOHYD	52	52	0.167065	.	.
PGIP1	netOGlyc-4.0.0.13	CARBOHYD	62	62	0.435372	.	.
PGIP1	netOGlyc-4.0.0.13	CARBOHYD	65	65	0.140508	.	.
PGIP1	netOGlyc-4.0.0.13	CARBOHYD	67	67	0.0699338	.	.
PGIP1	netOGlyc-4.0.0.13	CARBOHYD	69	69	0.171824	.	.
PGIP1	netOGlyc-4.0.0.13	CARBOHYD	78	78	0.0173664	.	.
PGIP1	netOGlyc-4.0.0.13	CARBOHYD	87	87	0.0655367	.	.
PGIP1	netOGlyc-4.0.0.13	CARBOHYD	91	91	0.0773823	.	.
PGIP1	netOGlyc-4.0.0.13	CARBOHYD	102	102	0.135274	.	.
PGIP1	netOGlyc-4.0.0.13	CARBOHYD	104	104	0.157077	.	.
PGIP1	netOGlyc-4.0.0.13	CARBOHYD	106	106	0.265962	.	.
PGIP1	netOGlyc-4.0.0.13	CARBOHYD	112	112	0.396215	.	.
PGIP1	netOGlyc-4.0.0.13	CARBOHYD	116	116	0.329586	.	.
PGIP1	netOGlyc-4.0.0.13	CARBOHYD	118	118	0.351908	.	.
PGIP1	netOGlyc-4.0.0.13	CARBOHYD	121	121	0.205909	.	.
PGIP1	netOGlyc-4.0.0.13	CARBOHYD	131	131	0.110461	.	.
PGIP1	netOGlyc-4.0.0.13	CARBOHYD	134	134	0.174633	.	.
PGIP1	netOGlyc-4.0.0.13	CARBOHYD	142	142	0.163045	.	.
PGIP1	netOGlyc-4.0.0.13	CARBOHYD	146	146	0.0851498	.	.
PGIP1	netOGlyc-4.0.0.13	CARBOHYD	153	153	0.0329424	.	.
PGIP1	netOGlyc-4.0.0.13	CARBOHYD	158	158	0.0362299	.	.
PGIP1	netOGlyc-4.0.0.13	CARBOHYD	167	167	0.0682406	.	.
PGIP1	netOGlyc-4.0.0.13	CARBOHYD	175	175	0.0942006	.	.
PGIP1	netOGlyc-4.0.0.13	CARBOHYD	182	182	0.172069	.	.
PGIP1	netOGlyc-4.0.0.13	CARBOHYD	188	188	0.0527801	.	.
PGIP1	netOGlyc-4.0.0.13	CARBOHYD	195	195	0.0426644	.	.
PGIP1	netOGlyc-4.0.0.13	CARBOHYD	200	200	0.149131	.	.
PGIP1	netOGlyc-4.0.0.13	CARBOHYD	202	202	0.0825999	.	.
PGIP1	netOGlyc-4.0.0.13	CARBOHYD	207	207	0.0827936	.	.
PGIP1	netOGlyc-4.0.0.13	CARBOHYD	213	213	0.227997	.	.
PGIP1	netOGlyc-4.0.0.13	CARBOHYD	225	225	0.0380826	.	.
PGIP1	netOGlyc-4.0.0.13	CARBOHYD	234	234	0.0306814	.	.
PGIP1	netOGlyc-4.0.0.13	CARBOHYD	239	239	0.100477	.	.
PGIP1	netOGlyc-4.0.0.13	CARBOHYD	243	243	0.112375	.	.
PGIP1	netOGlyc-4.0.0.13	CARBOHYD	252	252	0.0366045	.	.
PGIP1	netOGlyc-4.0.0.13	CARBOHYD	263	263	0.0865906	.	.
PGIP1	netOGlyc-4.0.0.13	CARBOHYD	265	265	0.169018	.	.
PGIP1	netOGlyc-4.0.0.13	CARBOHYD	272	272	0.201844	.	.
PGIP1	netOGlyc-4.0.0.13	CARBOHYD	279	279	0.175716	.	.
PGIP1	netOGlyc-4.0.0.13	CARBOHYD	285	285	0.0833163	.	.
PGIP1	netOGlyc-4.0.0.13	CARBOHYD	286	286	0.0931463	.	.
PGIP1	netOGlyc-4.0.0.13	CARBOHYD	296	296	0.0289366	.	.
PGIP1	netOGlyc-4.0.0.13	CARBOHYD	316	316	0.413738	.	.
PGIP1	netOGlyc-4.0.0.13	CARBOHYD	317	317	0.249855	.	.
PGIP1	netOGlyc-4.0.0.13	CARBOHYD	327	327	0.149154	.	.
PGIP1	netOGlyc-4.0.0.13	CARBOHYD	331	331	0.131789	.	.
```

**(*Gm*PGIP2)**

```
##gff-version 2
##source-version NetOGlyc 4.0.0.13
##date 22-2-19
##Type Protein
```

#seqname^a^	source^b^	feature^c^	start^d^	end^e^	score^f^	strand^g^	frame^h^	comment^i^ PGIP2	netOGlyc-4.0.0.13	CARBOHYD	5	5	0.285019	.	.

```
PGIP2	netOGlyc-4.0.0.13	CARBOHYD	21	21	0.0620045	.	.
PGIP2	netOGlyc-4.0.0.13	CARBOHYD	27	27	0.106577	.	.
PGIP2	netOGlyc-4.0.0.13	CARBOHYD	40	40	0.0532804	.	.
PGIP2	netOGlyc-4.0.0.13	CARBOHYD	42	42	0.0746558	.	.
PGIP2	netOGlyc-4.0.0.13	CARBOHYD	44	44	0.107543	.	.
PGIP2	netOGlyc-4.0.0.13	CARBOHYD	49	49	0.316814	.	.
PGIP2	netOGlyc-4.0.0.13	CARBOHYD	50	50	0.233589	.	.
PGIP2	netOGlyc-4.0.0.13	CARBOHYD	52	52	0.168602	.	.
PGIP2	netOGlyc-4.0.0.13	CARBOHYD	54	54	0.324339	.	.
PGIP2	netOGlyc-4.0.0.13	CARBOHYD	62	62	0.145439	.	.
PGIP2	netOGlyc-4.0.0.13	CARBOHYD	73	73	0.250146	.	.
PGIP2	netOGlyc-4.0.0.13	CARBOHYD	77	77	0.075417	.	.
PGIP2	netOGlyc-4.0.0.13	CARBOHYD	80	80	0.085196	.	.
PGIP2	netOGlyc-4.0.0.13	CARBOHYD	89	89	0.0592841	.	.
PGIP2	netOGlyc-4.0.0.13	CARBOHYD	100	100	0.0920973	.	.
PGIP2	netOGlyc-4.0.0.13	CARBOHYD	106	106	0.187581	.	.
PGIP2	netOGlyc-4.0.0.13	CARBOHYD	113	113	0.121582	.	.
PGIP2	netOGlyc-4.0.0.13	CARBOHYD	115	115	0.029837	.	.
PGIP2	netOGlyc-4.0.0.13	CARBOHYD	117	117	0.108007	.	.
PGIP2	netOGlyc-4.0.0.13	CARBOHYD	119	119	0.0691519	.	.
PGIP2	netOGlyc-4.0.0.13	CARBOHYD	122	122	0.0347433	.	.
PGIP2	netOGlyc-4.0.0.13	CARBOHYD	124	124	0.0815734	.	.
PGIP2	netOGlyc-4.0.0.13	CARBOHYD	131	131	0.0575329	.	.
PGIP2	netOGlyc-4.0.0.13	CARBOHYD	133	133	0.0300775	.	.
PGIP2	netOGlyc-4.0.0.13	CARBOHYD	138	138	0.0484255	.	.
PGIP2	netOGlyc-4.0.0.13	CARBOHYD	144	144	0.133615	.	.
PGIP2	netOGlyc-4.0.0.13	CARBOHYD	156	156	0.0586479	.	.
PGIP2	netOGlyc-4.0.0.13	CARBOHYD	167	167	0.043674	.	.
PGIP2	netOGlyc-4.0.0.13	CARBOHYD	178	178	0.286404	.	.
PGIP2	netOGlyc-4.0.0.13	CARBOHYD	180	180	0.190493	.	.
PGIP2	netOGlyc-4.0.0.13	CARBOHYD	187	187	0.130877	.	.
PGIP2	netOGlyc-4.0.0.13	CARBOHYD	194	194	0.0905052	.	.
PGIP2	netOGlyc-4.0.0.13	CARBOHYD	200	200	0.05998	.	.
PGIP2	netOGlyc-4.0.0.13	CARBOHYD	201	201	0.043964	.	.
PGIP2	netOGlyc-4.0.0.13	CARBOHYD	211	211	0.0611155	.	.
PGIP2	netOGlyc-4.0.0.13	CARBOHYD	219	219	0.547485	.	.	#POSITIVE
PGIP2	netOGlyc-4.0.0.13	CARBOHYD	231	231	0.170928	.	.
PGIP2	netOGlyc-4.0.0.13	CARBOHYD	232	232	0.0644876	.	.
```

**(*Gm*PGIP3)**

```
##gff-version 2
##source-version NetOGlyc 4.0.0.13
##date 22-2-19
##Type Protein
```

#seqname^a^	source^b^	feature^c^	start^d^	end^e^	score^f^	strand^g^	frame^h^	comment^i^ PGIP3	netOGlyc-4.0.0.13	CARBOHYD	2	2	0.139691	.	.

```
PGIP3	netOGlyc-4.0.0.13	CARBOHYD	5	5	0.0374852	.	.
PGIP3	netOGlyc-4.0.0.13	CARBOHYD	15	15	0.0891424	.	.
PGIP3	netOGlyc-4.0.0.13	CARBOHYD	17	17	0.08735	.	.
PGIP3	netOGlyc-4.0.0.13	CARBOHYD	21	21	0.0778075	.	.
PGIP3	netOGlyc-4.0.0.13	CARBOHYD	43	43	0.250842	.	.
PGIP3	netOGlyc-4.0.0.13	CARBOHYD	46	46	0.185174	.	.
PGIP3	netOGlyc-4.0.0.13	CARBOHYD	47	47	0.682001	.	.	#POSITIVE
PGIP3	netOGlyc-4.0.0.13	CARBOHYD	52	52	0.0515613	.	.
PGIP3	netOGlyc-4.0.0.13	CARBOHYD	63	63	0.318247	.	.
PGIP3	netOGlyc-4.0.0.13	CARBOHYD	66	66	0.204302	.	.
PGIP3	netOGlyc-4.0.0.13	CARBOHYD	68	68	0.0642075	.	.
PGIP3	netOGlyc-4.0.0.13	CARBOHYD	70	70	0.131251	.	.
PGIP3	netOGlyc-4.0.0.13	CARBOHYD	79	79	0.0240035	.	.
PGIP3	netOGlyc-4.0.0.13	CARBOHYD	84	84	0.102406	.	.
PGIP3	netOGlyc-4.0.0.13	CARBOHYD	92	92	0.102175	.	.
PGIP3	netOGlyc-4.0.0.13	CARBOHYD	105	105	0.292693	.	.
PGIP3	netOGlyc-4.0.0.13	CARBOHYD	107	107	0.254877	.	.
PGIP3	netOGlyc-4.0.0.13	CARBOHYD	113	113	0.232998	.	.
PGIP3	netOGlyc-4.0.0.13	CARBOHYD	116	116	0.520571	.	.	#POSITIVE
PGIP3	netOGlyc-4.0.0.13	CARBOHYD	117	117	0.356747	.	.
PGIP3	netOGlyc-4.0.0.13	CARBOHYD	119	119	0.407833	.	.
PGIP3	netOGlyc-4.0.0.13	CARBOHYD	122	122	0.225487	.	.
PGIP3	netOGlyc-4.0.0.13	CARBOHYD	132	132	0.088474	.	.
PGIP3	netOGlyc-4.0.0.13	CARBOHYD	133	133	0.225848	.	.
PGIP3	netOGlyc-4.0.0.13	CARBOHYD	135	135	0.278884	.	.
PGIP3	netOGlyc-4.0.0.13	CARBOHYD	143	143	0.259795	.	.
PGIP3	netOGlyc-4.0.0.13	CARBOHYD	147	147	0.0546035	.	.
PGIP3	netOGlyc-4.0.0.13	CARBOHYD	154	154	0.0232602	.	.
PGIP3	netOGlyc-4.0.0.13	CARBOHYD	159	159	0.0330734	.	.
PGIP3	netOGlyc-4.0.0.13	CARBOHYD	164	164	0.206648	.	.
PGIP3	netOGlyc-4.0.0.13	CARBOHYD	168	168	0.0979734	.	.
PGIP3	netOGlyc-4.0.0.13	CARBOHYD	183	183	0.136896	.	.
PGIP3	netOGlyc-4.0.0.13	CARBOHYD	189	189	0.0530441	.	.
PGIP3	netOGlyc-4.0.0.13	CARBOHYD	194	194	0.130315	.	.
PGIP3	netOGlyc-4.0.0.13	CARBOHYD	201	201	0.270726	.	.
PGIP3	netOGlyc-4.0.0.13	CARBOHYD	203	203	0.041935	.	.
PGIP3	netOGlyc-4.0.0.13	CARBOHYD	208	208	0.0450542	.	.
PGIP3	netOGlyc-4.0.0.13	CARBOHYD	213	213	0.183299	.	.
PGIP3	netOGlyc-4.0.0.13	CARBOHYD	214	214	0.247533	.	.
PGIP3	netOGlyc-4.0.0.13	CARBOHYD	222	222	0.0587386	.	.
PGIP3	netOGlyc-4.0.0.13	CARBOHYD	226	226	0.0537432	.	.
PGIP3	netOGlyc-4.0.0.13	CARBOHYD	235	235	0.0403272	.	.
PGIP3	netOGlyc-4.0.0.13	CARBOHYD	240	240	0.0571037	.	.
PGIP3	netOGlyc-4.0.0.13	CARBOHYD	244	244	0.0791049	.	.
PGIP3	netOGlyc-4.0.0.13	CARBOHYD	269	269	0.366571	.	.
PGIP3	netOGlyc-4.0.0.13	CARBOHYD	280	280	0.167221	.	.
PGIP3	netOGlyc-4.0.0.13	CARBOHYD	286	286	0.0942431	.	.
PGIP3	netOGlyc-4.0.0.13	CARBOHYD	287	287	0.0744777	.	.
PGIP3	netOGlyc-4.0.0.13	CARBOHYD	297	297	0.0313847	.	.
PGIP3	netOGlyc-4.0.0.13	CARBOHYD	317	317	0.102545	.	.
PGIP3	netOGlyc-4.0.0.13	CARBOHYD	328	328	0.0654874	.	.
PGIP3	netOGlyc-4.0.0.13	CARBOHYD	332	332	0.11642	.	.
```

**(*Gm*PGIP4)**

```
##gff-version 2
##source-version NetOGlyc 4.0.0.13
##date 22-2-18
##Type Protein
```

#seqname^a^	source^b^	feature^c^	start^d^	end^e^	score^f^	strand^g^	frame^h^	comment^i^

```
PGIP4	netOGlyc-4.0.0.13	CARBOHYD	2	2	0.12991	.	.
PGIP4	netOGlyc-4.0.0.13	CARBOHYD	5	5	0.0187181	.	.
PGIP4	netOGlyc-4.0.0.13	CARBOHYD	13	13	0.0440324	.	.
PGIP4	netOGlyc-4.0.0.13	CARBOHYD	15	15	0.0624475	.	.
PGIP4	netOGlyc-4.0.0.13	CARBOHYD	19	19	0.0440615	.	.
PGIP4	netOGlyc-4.0.0.13	CARBOHYD	29	29	0.11937	.	.
PGIP4	netOGlyc-4.0.0.13	CARBOHYD	41	41	0.374741	.	.
PGIP4	netOGlyc-4.0.0.13	CARBOHYD	42	42	0.322685	.	.
PGIP4	netOGlyc-4.0.0.13	CARBOHYD	44	44	0.288358	.	.
PGIP4	netOGlyc-4.0.0.13	CARBOHYD	45	45	0.741025	.	.	#POSITIVE
PGIP4	netOGlyc-4.0.0.13	CARBOHYD	50	50	0.0992123	.	.
PGIP4	netOGlyc-4.0.0.13	CARBOHYD	56	56	0.11016	.	.
PGIP4	netOGlyc-4.0.0.13	CARBOHYD	61	61	0.313486	.	.
PGIP4	netOGlyc-4.0.0.13	CARBOHYD	64	64	0.241357	.	.
PGIP4	netOGlyc-4.0.0.13	CARBOHYD	66	66	0.0777742	.	.
PGIP4	netOGlyc-4.0.0.13	CARBOHYD	68	68	0.150217	.	.
PGIP4	netOGlyc-4.0.0.13	CARBOHYD	77	77	0.0241823	.	.
PGIP4	netOGlyc-4.0.0.13	CARBOHYD	90	90	0.129016	.	.
PGIP4	netOGlyc-4.0.0.13	CARBOHYD	93	93	0.0509513	.	.
PGIP4	netOGlyc-4.0.0.13	CARBOHYD	103	103	0.25649	.	.
PGIP4	netOGlyc-4.0.0.13	CARBOHYD	111	111	0.190102	.	.
PGIP4	netOGlyc-4.0.0.13	CARBOHYD	114	114	0.470496	.	.
PGIP4	netOGlyc-4.0.0.13	CARBOHYD	115	115	0.355274	.	.
PGIP4	netOGlyc-4.0.0.13	CARBOHYD	117	117	0.410986	.	.
PGIP4	netOGlyc-4.0.0.13	CARBOHYD	120	120	0.202019	.	.
PGIP4	netOGlyc-4.0.0.13	CARBOHYD	130	130	0.0630914	.	.
PGIP4	netOGlyc-4.0.0.13	CARBOHYD	133	133	0.134926	.	.
PGIP4	netOGlyc-4.0.0.13	CARBOHYD	141	141	0.0707323	.	.
PGIP4	netOGlyc-4.0.0.13	CARBOHYD	152	152	0.0337603	.	.
PGIP4	netOGlyc-4.0.0.13	CARBOHYD	157	157	0.0383514	.	.
PGIP4	netOGlyc-4.0.0.13	CARBOHYD	162	162	0.244587	.	.
PGIP4	netOGlyc-4.0.0.13	CARBOHYD	166	166	0.107073	.	.
PGIP4	netOGlyc-4.0.0.13	CARBOHYD	174	174	0.0356626	.	.
PGIP4	netOGlyc-4.0.0.13	CARBOHYD	181	181	0.0725944	.	.
PGIP4	netOGlyc-4.0.0.13	CARBOHYD	190	190	0.0386204	.	.
PGIP4	netOGlyc-4.0.0.13	CARBOHYD	192	192	0.060602	.	.
PGIP4	netOGlyc-4.0.0.13	CARBOHYD	199	199	0.22837	.	.
PGIP4	netOGlyc-4.0.0.13	CARBOHYD	201	201	0.0495568	.	.
PGIP4	netOGlyc-4.0.0.13	CARBOHYD	212	212	0.17359	.	.
PGIP4	netOGlyc-4.0.0.13	CARBOHYD	224	224	0.041682	.	.
PGIP4	netOGlyc-4.0.0.13	CARBOHYD	233	233	0.0390995	.	.
PGIP4	netOGlyc-4.0.0.13	CARBOHYD	238	238	0.0759056	.	.
PGIP4	netOGlyc-4.0.0.13	CARBOHYD	242	242	0.140526	.	.
PGIP4	netOGlyc-4.0.0.13	CARBOHYD	262	262	0.14559	.	.
PGIP4	netOGlyc-4.0.0.13	CARBOHYD	264	264	0.124133	.	.
PGIP4	netOGlyc-4.0.0.13	CARBOHYD	278	278	0.117633	.	.
PGIP4	netOGlyc-4.0.0.13	CARBOHYD	284	284	0.0613649	.	.
PGIP4	netOGlyc-4.0.0.13	CARBOHYD	285	285	0.0801022	.	.
PGIP4	netOGlyc-4.0.0.13	CARBOHYD	295	295	0.0190512	.	.
PGIP4	netOGlyc-4.0.0.13	CARBOHYD	315	315	0.162948	.	.
PGIP4	netOGlyc-4.0.0.13	CARBOHYD	326	326	0.0779128	.	.
PGIP4	  netOGlyc-4.0.0.13  CARBOHYD	 330	   330	     0.116629	.	.
```

**(*Gm*PGIP5)**

```
##gff-version 2
##source-version NetOGlyc 4.0.0.13
##date 22-2-19
##Type Protein
```

#seqname^a^	source^b^	feature^c^	start^d^	end^e^	score^f^	strand^g^	frame^h^	comment^i^ PGIP5	netOGlyc-4.0.0.13	CARBOHYD	12	12	0.692175	.	.	#POSITIVE

```
PGIP5	netOGlyc-4.0.0.13	CARBOHYD	15	15	0.542454	.	.	#POSITIVE
PGIP5	netOGlyc-4.0.0.13	CARBOHYD	26	26	0.0988309	.	.
PGIP5	netOGlyc-4.0.0.13	CARBOHYD	39	39	0.104798	.	.
PGIP5	netOGlyc-4.0.0.13	CARBOHYD	50	50	0.113261	.	.
PGIP5	netOGlyc-4.0.0.13	CARBOHYD	53	53	0.113435	.	.
PGIP5	netOGlyc-4.0.0.13	CARBOHYD	54	54	0.185613	.	.
PGIP5	netOGlyc-4.0.0.13	CARBOHYD	60	60	0.237932	.	.
PGIP5	netOGlyc-4.0.0.13	CARBOHYD	63	63	0.613237	.	.	#POSITIVE
PGIP5	netOGlyc-4.0.0.13	CARBOHYD	64	64	0.336403	.	.
PGIP5	netOGlyc-4.0.0.13	CARBOHYD	66	66	0.417473	.	.
PGIP5	netOGlyc-4.0.0.13	CARBOHYD	69	69	0.150541	.	.
PGIP5	netOGlyc-4.0.0.13	CARBOHYD	79	79	0.105283	.	.
PGIP5	netOGlyc-4.0.0.13	CARBOHYD	82	82	0.157774	.	.
PGIP5	netOGlyc-4.0.0.13	CARBOHYD	90	90	0.162872	.	.
PGIP5	netOGlyc-4.0.0.13	CARBOHYD	94	94	0.103515	.	.
PGIP5	netOGlyc-4.0.0.13	CARBOHYD	101	101	0.0360346	.	.
PGIP5	netOGlyc-4.0.0.13	CARBOHYD	108	108	0.0355527	.	.
PGIP5	netOGlyc-4.0.0.13	CARBOHYD	117	117	0.0944908	.	.
PGIP5	netOGlyc-4.0.0.13	CARBOHYD	122	122	0.0710341	.	.
PGIP5	netOGlyc-4.0.0.13	CARBOHYD	125	125	0.0871464	.	.
PGIP5	netOGlyc-4.0.0.13	CARBOHYD	132	132	0.241125	.	.
PGIP5	netOGlyc-4.0.0.13	CARBOHYD	134	134	0.0639483	.	.
PGIP5	netOGlyc-4.0.0.13	CARBOHYD	136	136	0.0783221	.	.
PGIP5	netOGlyc-4.0.0.13	CARBOHYD	138	138	0.0531838	.	.
PGIP5	netOGlyc-4.0.0.13	CARBOHYD	140	140	0.0379705	.	.
```

**(*Gm*PGIP6)**

```
##gff-version 2
##source-version NetOGlyc 4.0.0.13
##date 22-2-18
##Type Protein
```

#seqname^a^	source^b^	feature^c^	start^d^	end^e^	score^f^	strand^g^	frame^h^	comment^i^

```
PGIP6	netOGlyc-4.0.0.13	CARBOHYD	2	2	0.252197	.	.
PGIP6	netOGlyc-4.0.0.13	CARBOHYD	4	4	0.05439	.	.
PGIP6	netOGlyc-4.0.0.13	CARBOHYD	5	5	0.12634	.	.
PGIP6	netOGlyc-4.0.0.13	CARBOHYD	6	6	0.036501	.	.
PGIP6	netOGlyc-4.0.0.13	CARBOHYD	17	17	0.0984094	.	.
PGIP6	netOGlyc-4.0.0.13	CARBOHYD	21	21	0.0720879	.	.
PGIP6	netOGlyc-4.0.0.13	CARBOHYD	43	43	0.309031	.	.
PGIP6	netOGlyc-4.0.0.13	CARBOHYD	44	44	0.280976	.	.
PGIP6	netOGlyc-4.0.0.13	CARBOHYD	46	46	0.241123	.	.
PGIP6	netOGlyc-4.0.0.13	CARBOHYD	47	47	0.511318	.	.	#POSITIVE
PGIP6	netOGlyc-4.0.0.13	CARBOHYD	51	51	0.19477	.	.
PGIP6	netOGlyc-4.0.0.13	CARBOHYD	52	52	0.182987	.	.
PGIP6	netOGlyc-4.0.0.13	CARBOHYD	63	63	0.28425	.	.
PGIP6	netOGlyc-4.0.0.13	CARBOHYD	68	68	0.0592674	.	.
PGIP6	netOGlyc-4.0.0.13	CARBOHYD	70	70	0.19109	.	.
PGIP6	netOGlyc-4.0.0.13	CARBOHYD	92	92	0.088854	.	.
PGIP6	netOGlyc-4.0.0.13	CARBOHYD	103	103	0.0783322	.	.
PGIP6	netOGlyc-4.0.0.13	CARBOHYD	105	105	0.079148	.	.
PGIP6	netOGlyc-4.0.0.13	CARBOHYD	107	107	0.162485	.	.
PGIP6	netOGlyc-4.0.0.13	CARBOHYD	113	113	0.403148	.	.
PGIP6	netOGlyc-4.0.0.13	CARBOHYD	117	117	0.308201	.	.
PGIP6	netOGlyc-4.0.0.13	CARBOHYD	119	119	0.294086	.	.
PGIP6	netOGlyc-4.0.0.13	CARBOHYD	122	122	0.0933981	.	.
PGIP6	netOGlyc-4.0.0.13	CARBOHYD	132	132	0.09119	.	.
PGIP6	netOGlyc-4.0.0.13	CARBOHYD	135	135	0.177907	.	.
PGIP6	netOGlyc-4.0.0.13	CARBOHYD	143	143	0.135771	.	.
PGIP6	netOGlyc-4.0.0.13	CARBOHYD	147	147	0.0616554	.	.
PGIP6	netOGlyc-4.0.0.13	CARBOHYD	150	150	0.088714	.	.
PGIP6	netOGlyc-4.0.0.13	CARBOHYD	159	159	0.0426208	.	.
PGIP6	netOGlyc-4.0.0.13	CARBOHYD	168	168	0.0828409	.	.
PGIP6	netOGlyc-4.0.0.13	CARBOHYD	176	176	0.214792	.	.
PGIP6	netOGlyc-4.0.0.13	CARBOHYD	183	183	0.133708	.	.
PGIP6	netOGlyc-4.0.0.13	CARBOHYD	185	185	0.0310627	.	.
PGIP6	netOGlyc-4.0.0.13	CARBOHYD	189	189	0.0489793	.	.
PGIP6	netOGlyc-4.0.0.13	CARBOHYD	192	192	0.0787931	.	.
PGIP6	netOGlyc-4.0.0.13	CARBOHYD	194	194	0.0450661	.	.
PGIP6	netOGlyc-4.0.0.13	CARBOHYD	196	196	0.0568902	.	.
PGIP6	netOGlyc-4.0.0.13	CARBOHYD	201	201	0.159497	.	.
PGIP6	netOGlyc-4.0.0.13	CARBOHYD	203	203	0.0816698	.	.
PGIP6	netOGlyc-4.0.0.13	CARBOHYD	208	208	0.0778229	.	.
PGIP6	netOGlyc-4.0.0.13	CARBOHYD	214	214	0.156287	.	.
PGIP6	netOGlyc-4.0.0.13	CARBOHYD	226	226	0.0201051	.	.
PGIP6	netOGlyc-4.0.0.13	CARBOHYD	235	235	0.0186465	.	.
PGIP6	netOGlyc-4.0.0.13	CARBOHYD	240	240	0.0558958	.	.
PGIP6	netOGlyc-4.0.0.13	CARBOHYD	244	244	0.0945654	.	.
PGIP6	netOGlyc-4.0.0.13	CARBOHYD	253	253	0.0482411	.	.
PGIP6	netOGlyc-4.0.0.13	CARBOHYD	264	264	0.162244	.	.
PGIP6	netOGlyc-4.0.0.13	CARBOHYD	266	266	0.147964	.	.
PGIP6	netOGlyc-4.0.0.13	CARBOHYD	273	273	0.154883	.	.
PGIP6	netOGlyc-4.0.0.13	CARBOHYD	286	286	0.0527557	.	.
PGIP6	netOGlyc-4.0.0.13	CARBOHYD	287	287	0.0490401	.	.
PGIP6	netOGlyc-4.0.0.13	CARBOHYD	293	293	0.0624566	.	.
PGIP6	netOGlyc-4.0.0.13	CARBOHYD	297	297	0.0442215	.	.
PGIP6	netOGlyc-4.0.0.13	CARBOHYD	317	317	0.310195	.	.
PGIP6	netOGlyc-4.0.0.13	CARBOHYD	318	318	0.194824	.	.
PGIP6	netOGlyc-4.0.0.13	CARBOHYD	320	320	0.250087	.	.
PGIP6	netOGlyc-4.0.0.13	CARBOHYD	328	328	0.174549	.	.
```

**(*Gm*PGIP7)**

```
##gff-version 2
##source-version NetOGlyc 4.0.0.13
##date 22-2-19
##Type Protein
```

#seqname^a^	source^b^	feature^c^	start^d^	end^e^	score^f^	strand^g^	frame^h^	comment^i^ PGIP7	netOGlyc-4.0.0.13	CARBOHYD	2	2	0.0680714	.	.

```
PGIP7	netOGlyc-4.0.0.13	CARBOHYD	5	5	0.0343643	.	.
PGIP7	netOGlyc-4.0.0.13	CARBOHYD	13	13	0.04591	.	.
PGIP7	netOGlyc-4.0.0.13	CARBOHYD	15	15	0.0597359	.	.
PGIP7	netOGlyc-4.0.0.13	CARBOHYD	16	16	0.0478599	.	.
PGIP7	netOGlyc-4.0.0.13	CARBOHYD	19	19	0.0333539	.	.
PGIP7	netOGlyc-4.0.0.13	CARBOHYD	41	41	0.319113	.	.
PGIP7	netOGlyc-4.0.0.13	CARBOHYD	42	42	0.4194	.	.
PGIP7	netOGlyc-4.0.0.13	CARBOHYD	44	44	0.306094	.	.
PGIP7	netOGlyc-4.0.0.13	CARBOHYD	45	45	0.483609	.	.
PGIP7	netOGlyc-4.0.0.13	CARBOHYD	49	49	0.202138	.	.
PGIP7	netOGlyc-4.0.0.13	CARBOHYD	50	50	0.272711	.	.
PGIP7	netOGlyc-4.0.0.13	CARBOHYD	56	56	0.119466	.	.
PGIP7	netOGlyc-4.0.0.13	CARBOHYD	61	61	0.371904	.	.
PGIP7	netOGlyc-4.0.0.13	CARBOHYD	64	64	0.0934587	.	.
PGIP7	netOGlyc-4.0.0.13	CARBOHYD	66	66	0.0220972	.	.
PGIP7	netOGlyc-4.0.0.13	CARBOHYD	68	68	0.0634477	.	.
PGIP7	netOGlyc-4.0.0.13	CARBOHYD	77	77	0.0147825	.	.
PGIP7	netOGlyc-4.0.0.13	CARBOHYD	86	86	0.191161	.	.
PGIP7	netOGlyc-4.0.0.13	CARBOHYD	90	90	0.122177	.	.
PGIP7	netOGlyc-4.0.0.13	CARBOHYD	101	101	0.278186	.	.
PGIP7	netOGlyc-4.0.0.13	CARBOHYD	103	103	0.26925	.	.
PGIP7	netOGlyc-4.0.0.13	CARBOHYD	105	105	0.108051	.	.
PGIP7	netOGlyc-4.0.0.13	CARBOHYD	107	107	0.294973	.	.
PGIP7	netOGlyc-4.0.0.13	CARBOHYD	114	114	0.5	.	.	#POSITIVE
PGIP7	netOGlyc-4.0.0.13	CARBOHYD	120	120	0.130757	.	.
PGIP7	netOGlyc-4.0.0.13	CARBOHYD	128	128	0.104984	.	.
PGIP7	netOGlyc-4.0.0.13	CARBOHYD	130	130	0.140033	.	.
PGIP7	netOGlyc-4.0.0.13	CARBOHYD	133	133	0.210539	.	.
PGIP7	netOGlyc-4.0.0.13	CARBOHYD	141	141	0.0988745	.	.
PGIP7	netOGlyc-4.0.0.13	CARBOHYD	145	145	0.164236	.	.
PGIP7	netOGlyc-4.0.0.13	CARBOHYD	148	148	0.318523	.	.
PGIP7	netOGlyc-4.0.0.13	CARBOHYD	152	152	0.105882	.	.
PGIP7	netOGlyc-4.0.0.13	CARBOHYD	155	155	0.320724	.	.
PGIP7	netOGlyc-4.0.0.13	CARBOHYD	157	157	0.0572443	.	.
PGIP7	netOGlyc-4.0.0.13	CARBOHYD	163	163	0.321575	.	.
PGIP7	netOGlyc-4.0.0.13	CARBOHYD	165	165	0.163596	.	.
PGIP7	netOGlyc-4.0.0.13	CARBOHYD	166	166	0.0724671	.	.
PGIP7	netOGlyc-4.0.0.13	CARBOHYD	174	174	0.0560396	.	.
PGIP7	netOGlyc-4.0.0.13	CARBOHYD	181	181	0.112826	.	.
PGIP7	netOGlyc-4.0.0.13	CARBOHYD	187	187	0.0629567	.	.
PGIP7	netOGlyc-4.0.0.13	CARBOHYD	190	190	0.0531142	.	.
PGIP7	netOGlyc-4.0.0.13	CARBOHYD	192	192	0.0392374	.	.
PGIP7	netOGlyc-4.0.0.13	CARBOHYD	196	196	0.126254	.	.
PGIP7	netOGlyc-4.0.0.13	CARBOHYD	197	197	0.30606	.	.
PGIP7	netOGlyc-4.0.0.13	CARBOHYD	199	199	0.323886	.	.
PGIP7	netOGlyc-4.0.0.13	CARBOHYD	201	201	0.193772	.	.
PGIP7	netOGlyc-4.0.0.13	CARBOHYD	206	206	0.204859	.	.
PGIP7	netOGlyc-4.0.0.13	CARBOHYD	212	212	0.124555	.	.
PGIP7	netOGlyc-4.0.0.13	CARBOHYD	224	224	0.028954	.	.
PGIP7	netOGlyc-4.0.0.13	CARBOHYD	233	233	0.0297207	.	.
PGIP7	netOGlyc-4.0.0.13	CARBOHYD	238	238	0.0506281	.	.
PGIP7	netOGlyc-4.0.0.13	CARBOHYD	242	242	0.0890644	.	.
PGIP7	netOGlyc-4.0.0.13	CARBOHYD	262	262	0.14361	.	.
PGIP7	netOGlyc-4.0.0.13	CARBOHYD	278	278	0.105499	.	.
PGIP7	netOGlyc-4.0.0.13	CARBOHYD	284	284	0.136852	.	.
PGIP7	netOGlyc-4.0.0.13	CARBOHYD	291	291	0.104451	.	.
PGIP7	netOGlyc-4.0.0.13	CARBOHYD	295	295	0.0233434	.	.
PGIP7	netOGlyc-4.0.0.13	CARBOHYD	315	315	0.1614	.	.
PGIP7	netOGlyc-4.0.0.13	CARBOHYD	316	316	0.150296	.	.
PGIP7	netOGlyc-4.0.0.13	CARBOHYD	326	326	0.100809	.	.
PGIP7	netOGlyc-4.0.0.13	CARBOHYD	332	332	0.0255544	.	.
```

**(*Gm*PGIP8)**

```
##gff-version 2
##source-version NetOGlyc 4.0.0.13
##date 22-2-19
##Type Protein
```

#seqname^a^	source^b^	feature^c^	start^d^	end^e^	score^f^	strand^g^	frame^h^	comment^i^ PGIP8	netOGlyc-4.0.0.13	CARBOHYD	12	12	0.161868	.	.

```
PGIP8	netOGlyc-4.0.0.13	CARBOHYD	17	17	0.0230483	.	.
PGIP8	netOGlyc-4.0.0.13	CARBOHYD	38	38	0.307065	.	.
PGIP8	netOGlyc-4.0.0.13	CARBOHYD	41	41	0.460063	.	.
PGIP8	netOGlyc-4.0.0.13	CARBOHYD	42	42	0.715066	.	.	#POSITIVE
PGIP8	netOGlyc-4.0.0.13	CARBOHYD	46	46	0.296107	.	.
PGIP8	netOGlyc-4.0.0.13	CARBOHYD	47	47	0.232091	.	.
PGIP8	netOGlyc-4.0.0.13	CARBOHYD	52	52	0.105953	.	.
PGIP8	netOGlyc-4.0.0.13	CARBOHYD	61	61	0.0902718	.	.
PGIP8	netOGlyc-4.0.0.13	CARBOHYD	63	63	0.0465949	.	.
PGIP8	netOGlyc-4.0.0.13	CARBOHYD	67	67	0.0448161	.	.
PGIP8	netOGlyc-4.0.0.13	CARBOHYD	76	76	0.0369092	.	.
PGIP8	netOGlyc-4.0.0.13	CARBOHYD	89	89	0.124194	.	.
PGIP8	netOGlyc-4.0.0.13	CARBOHYD	104	104	0.018307	.	.
PGIP8	netOGlyc-4.0.0.13	CARBOHYD	112	112	0.106086	.	.
PGIP8	netOGlyc-4.0.0.13	CARBOHYD	113	113	0.108359	.	.
PGIP8	netOGlyc-4.0.0.13	CARBOHYD	118	118	0.0387492	.	.
PGIP8	netOGlyc-4.0.0.13	CARBOHYD	126	126	0.0441527	.	.
PGIP8	netOGlyc-4.0.0.13	CARBOHYD	128	128	0.0820897	.	.
PGIP8	netOGlyc-4.0.0.13	CARBOHYD	131	131	0.112732	.	.
PGIP8	netOGlyc-4.0.0.13	CARBOHYD	133	133	0.0680043	.	.
PGIP8	netOGlyc-4.0.0.13	CARBOHYD	139	139	0.111967	.	.
PGIP8	netOGlyc-4.0.0.13	CARBOHYD	143	143	0.159215	.	.
PGIP8	netOGlyc-4.0.0.13	CARBOHYD	146	146	0.233628	.	.
PGIP8	netOGlyc-4.0.0.13	CARBOHYD	150	150	0.0407331	.	.
PGIP8	netOGlyc-4.0.0.13	CARBOHYD	155	155	0.0290343	.	.
PGIP8	netOGlyc-4.0.0.13	CARBOHYD	161	161	0.261302	.	.
PGIP8	netOGlyc-4.0.0.13	CARBOHYD	163	163	0.0555377	.	.
PGIP8	netOGlyc-4.0.0.13	CARBOHYD	164	164	0.0632985	.	.
PGIP8	netOGlyc-4.0.0.13	CARBOHYD	174	174	0.0636885	.	.
PGIP8	netOGlyc-4.0.0.13	CARBOHYD	179	179	0.232755	.	.
PGIP8	netOGlyc-4.0.0.13	CARBOHYD	185	185	0.025536	.	.
PGIP8	netOGlyc-4.0.0.13	CARBOHYD	188	188	0.0175824	.	.
PGIP8	netOGlyc-4.0.0.13	CARBOHYD	190	190	0.0201917	.	.
PGIP8	netOGlyc-4.0.0.13	CARBOHYD	199	199	0.0863769	.	.
PGIP8	netOGlyc-4.0.0.13	CARBOHYD	204	204	0.100916	.	.
PGIP8	netOGlyc-4.0.0.13	CARBOHYD	210	210	0.269563	.	.
PGIP8	netOGlyc-4.0.0.13	CARBOHYD	222	222	0.0411312	.	.
PGIP8	netOGlyc-4.0.0.13	CARBOHYD	231	231	0.0297619	.	.
PGIP8	netOGlyc-4.0.0.13	CARBOHYD	236	236	0.0572161	.	.
PGIP8	netOGlyc-4.0.0.13	CARBOHYD	240	240	0.0440343	.	.
PGIP8	netOGlyc-4.0.0.13	CARBOHYD	260	260	0.0867591	.	.
PGIP8	netOGlyc-4.0.0.13	CARBOHYD	276	276	0.136941	.	.
PGIP8	netOGlyc-4.0.0.13	CARBOHYD	282	282	0.199574	.	.
PGIP8	netOGlyc-4.0.0.13	CARBOHYD	288	288	0.0340179	.	.
PGIP8	netOGlyc-4.0.0.13	CARBOHYD	293	293	0.0420164	.	.
PGIP8	netOGlyc-4.0.0.13	CARBOHYD	314	314	0.216642	.	.
PGIP8	netOGlyc-4.0.0.13	CARBOHYD	324	324	0.11433	.	.
PGIP8	netOGlyc-4.0.0.13	CARBOHYD	330	330	0.0536843	.	.
```

**(*Gm*PGIP9)**

```
##gff-version 2
##source-version NetOGlyc 4.0.0.13
##date 22-2-19
##Type Protein
```

#seqname^a^	source^b^	feature^c^	start^d^	end^e^	score^f^	strand^g^	frame^h^	comment^i^

```
PGIP9	netOGlyc-4.0.0.13	CARBOHYD	6	6	0.814914	.	.	#POSITIVE
PGIP9	netOGlyc-4.0.0.13	CARBOHYD	18	18	0.522934	.	.	#POSITIVE
PGIP9	netOGlyc-4.0.0.13	CARBOHYD	28	28	0.393419	.	.
PGIP9	netOGlyc-4.0.0.13	CARBOHYD	33	33	0.0735839	.	.
PGIP9	netOGlyc-4.0.0.13	CARBOHYD	41	41	0.0867796	.	.
PGIP9	netOGlyc-4.0.0.13	CARBOHYD	46	46	0.105559	.	.
PGIP9	netOGlyc-4.0.0.13	CARBOHYD	53	53	0.0406352	.	.
PGIP9	netOGlyc-4.0.0.13	CARBOHYD	55	55	0.0647796	.	.
PGIP9	netOGlyc-4.0.0.13	CARBOHYD	58	58	0.0658055	.	.
PGIP9	netOGlyc-4.0.0.13	CARBOHYD	59	59	0.226437	.	.
PGIP9	netOGlyc-4.0.0.13	CARBOHYD	84	84	0.153186	.	.
PGIP9	netOGlyc-4.0.0.13	CARBOHYD	85	85	0.303214	.	.
PGIP9	netOGlyc-4.0.0.13	CARBOHYD	105	105	0.00691951	.	.
PGIP9	netOGlyc-4.0.0.13	CARBOHYD	110	110	0.0175882	.	.
PGIP9	netOGlyc-4.0.0.13	CARBOHYD	114	114	0.140895	.	.
PGIP9	netOGlyc-4.0.0.13	CARBOHYD	115	115	0.0968211	.	.
PGIP9	netOGlyc-4.0.0.13	CARBOHYD	116	116	0.106169	.	.
PGIP9	netOGlyc-4.0.0.13	CARBOHYD	123	123	0.0535263	.	.
PGIP9	netOGlyc-4.0.0.13	CARBOHYD	129	129	0.384881	.	.
PGIP9	netOGlyc-4.0.0.13	CARBOHYD	138	138	0.0850425	.	.
PGIP9	netOGlyc-4.0.0.13	CARBOHYD	140	140	0.0664437	.	.
PGIP9	netOGlyc-4.0.0.13	CARBOHYD	154	154	0.296489	.	.
PGIP9	netOGlyc-4.0.0.13	CARBOHYD	159	159	0.111578	.	.
PGIP9	netOGlyc-4.0.0.13	CARBOHYD	163	163	0.322269	.	.
PGIP9	netOGlyc-4.0.0.13	CARBOHYD	167	167	0.100531	.	.
PGIP9	netOGlyc-4.0.0.13	CARBOHYD	169	169	0.117935	.	.
PGIP9	netOGlyc-4.0.0.13	CARBOHYD	172	172	0.0976109	.	.
PGIP9	netOGlyc-4.0.0.13	CARBOHYD	191	191	0.017909	.	.
PGIP9	netOGlyc-4.0.0.13	CARBOHYD	196	196	0.0495182	.	.
PGIP9	netOGlyc-4.0.0.13	CARBOHYD	201	201	0.219603	.	.
PGIP9	netOGlyc-4.0.0.13	CARBOHYD	202	202	0.122675	.	.
PGIP9	netOGlyc-4.0.0.13	CARBOHYD	204	204	0.0789921	.	.
PGIP9	netOGlyc-4.0.0.13	CARBOHYD	211	211	0.198689	.	.
PGIP9	netOGlyc-4.0.0.13	CARBOHYD	220	220	0.140558	.	.
PGIP9	netOGlyc-4.0.0.13	CARBOHYD	225	225	0.183102	.	.
PGIP9	netOGlyc-4.0.0.13	CARBOHYD	226	226	0.240444	.	.
PGIP9	netOGlyc-4.0.0.13	CARBOHYD	240	240	0.0516758	.	.
PGIP9	netOGlyc-4.0.0.13	CARBOHYD	245	245	0.116495	.	.
PGIP9	netOGlyc-4.0.0.13	CARBOHYD	251	251	0.163959	.	.
PGIP9	netOGlyc-4.0.0.13	CARBOHYD	263	263	0.0450118	.	.
PGIP9	netOGlyc-4.0.0.13	CARBOHYD	272	272	0.0601296	.	.
PGIP9	netOGlyc-4.0.0.13	CARBOHYD	276	276	0.0785803	.	.
PGIP9	netOGlyc-4.0.0.13	CARBOHYD	277	277	0.141662	.	.
PGIP9	netOGlyc-4.0.0.13	CARBOHYD	280	280	0.286693	.	.
PGIP9	netOGlyc-4.0.0.13	CARBOHYD	281	281	0.0712191	.	.
PGIP9	netOGlyc-4.0.0.13	CARBOHYD	287	287	0.101333	.	.
PGIP9	netOGlyc-4.0.0.13	CARBOHYD	296	296	0.0449499	.	.
PGIP9	netOGlyc-4.0.0.13	CARBOHYD	299	299	0.0526961	.	.
PGIP9	netOGlyc-4.0.0.13	CARBOHYD	304	304	0.236609	.	.
PGIP9	netOGlyc-4.0.0.13	CARBOHYD	324	324	0.256211	.	.
PGIP9	netOGlyc-4.0.0.13	CARBOHYD	325	325	0.0222227	.	.
PGIP9	netOGlyc-4.0.0.13	CARBOHYD	335	335	0.308968	.	.
PGIP9	netOGlyc-4.0.0.13	CARBOHYD	341	341	0.0890681	.	.
PGIP9	netOGlyc-4.0.0.13	CARBOHYD	361	361	0.0712182	.	.
PGIP9	netOGlyc-4.0.0.13	CARBOHYD	366	366	0.117992	.	.
```

**(*Gm*PGIP10)**

```
##gff-version 2
##source-version NetOGlyc 4.0.0.13
##date 22-2-19
##Type Protein
```

#seqname^a^	source^b^	feature^c^	start^d^	end^e^	score^f^	strand^g^	frame^h^	comment^i^

```
PGIP10	netOGlyc-4.0.0.13	CARBOHYD	13	13	0.0898442	.	.
PGIP10	netOGlyc-4.0.0.13	CARBOHYD	20	20	0.105951	.	.
PGIP10	netOGlyc-4.0.0.13	CARBOHYD	46	46	0.355347	.	.
PGIP10	netOGlyc-4.0.0.13	CARBOHYD	66	66	0.0114181	.	.
PGIP10	netOGlyc-4.0.0.13	CARBOHYD	71	71	0.0526356	.	.
PGIP10	netOGlyc-4.0.0.13	CARBOHYD	75	75	0.190997	.	.
PGIP10	netOGlyc-4.0.0.13	CARBOHYD	76	76	0.0853981	.	.
PGIP10	netOGlyc-4.0.0.13	CARBOHYD	81	81	0.0283776	.	.
PGIP10	netOGlyc-4.0.0.13	CARBOHYD	84	84	0.0570623	.	.
PGIP10	netOGlyc-4.0.0.13	CARBOHYD	90	90	0.322435	.	.
PGIP10	netOGlyc-4.0.0.13	CARBOHYD	99	99	0.129973	.	.
PGIP10	netOGlyc-4.0.0.13	CARBOHYD	120	120	0.0751984	.	.
PGIP10	netOGlyc-4.0.0.13	CARBOHYD	128	128	0.0378474	.	.
PGIP10	netOGlyc-4.0.0.13	CARBOHYD	133	133	0.0284886	.	.
PGIP10	netOGlyc-4.0.0.13	CARBOHYD	152	152	0.0118833	.	.
PGIP10	netOGlyc-4.0.0.13	CARBOHYD	157	157	0.0481331	.	.
PGIP10	netOGlyc-4.0.0.13	CARBOHYD	162	162	0.17747	.	.
PGIP10	netOGlyc-4.0.0.13	CARBOHYD	163	163	0.149016	.	.
PGIP10	netOGlyc-4.0.0.13	CARBOHYD	176	176	0.0597594	.	.
PGIP10	netOGlyc-4.0.0.13	CARBOHYD	181	181	0.167594	.	.
PGIP10	netOGlyc-4.0.0.13	CARBOHYD	183	183	0.0940589	.	.
PGIP10	netOGlyc-4.0.0.13	CARBOHYD	187	187	0.243155	.	.
PGIP10	netOGlyc-4.0.0.13	CARBOHYD	190	190	0.0768798	.	.
PGIP10	netOGlyc-4.0.0.13	CARBOHYD	201	201	0.0461642	.	.
PGIP10	netOGlyc-4.0.0.13	CARBOHYD	206	206	0.0797479	.	.
PGIP10	netOGlyc-4.0.0.13	CARBOHYD	212	212	0.234128	.	.
PGIP10	netOGlyc-4.0.0.13	CARBOHYD	218	218	0.0404947	.	.
PGIP10	netOGlyc-4.0.0.13	CARBOHYD	224	224	0.0556408	.	.
PGIP10	netOGlyc-4.0.0.13	CARBOHYD	233	233	0.0329317	.	.
PGIP10	netOGlyc-4.0.0.13	CARBOHYD	238	238	0.0918819	.	.
PGIP10	netOGlyc-4.0.0.13	CARBOHYD	242	242	0.0995071	.	.
PGIP10	netOGlyc-4.0.0.13	CARBOHYD	244	244	0.117339	.	.
PGIP10	netOGlyc-4.0.0.13	CARBOHYD	248	248	0.0686847	.	.
PGIP10	netOGlyc-4.0.0.13	CARBOHYD	257	257	0.0683471	.	.
PGIP10	netOGlyc-4.0.0.13	CARBOHYD	265	265	0.510159	.	.	#POSITIVE
PGIP10	netOGlyc-4.0.0.13	CARBOHYD	279	279	0.166788	.	.
PGIP10	netOGlyc-4.0.0.13	CARBOHYD	285	285	0.0543536	.	.
PGIP10	netOGlyc-4.0.0.13	CARBOHYD	296	296	0.036813	.	.
PGIP10	netOGlyc-4.0.0.13	CARBOHYD	316	316	0.11373	.	.
PGIP10	netOGlyc-4.0.0.13	CARBOHYD	319	319	0.0614034	.	.
PGIP10	netOGlyc-4.0.0.13	CARBOHYD	327	327	0.14565	.	.
PGIP10	netOGlyc-4.0.0.13	CARBOHYD	330	330	0.0396197	.	.
PGIP10	netOGlyc-4.0.0.13	CARBOHYD	338	338	0.0555318	.	.
```

**(*Gm*PGIP11)**

```
##gff-version 2
##source-version NetOGlyc 4.0.0.13
##date 22-2-18
##Type Protein
```

#seqname^a^	source^b^	feature^c^	start^d^	end^e^	score^f^	strand^g^	frame^h^	comment^i^

```
PGIP11	netOGlyc-4.0.0.13	CARBOHYD	5	5	0.0830737	.	.
PGIP11	netOGlyc-4.0.0.13	CARBOHYD	13	13	0.0163051	.	.
PGIP11	netOGlyc-4.0.0.13	CARBOHYD	15	15	0.0296026	.	.
PGIP11	netOGlyc-4.0.0.13	CARBOHYD	17	17	0.0344033	.	.
PGIP11	netOGlyc-4.0.0.13	CARBOHYD	21	21	0.126429	.	.
PGIP11	netOGlyc-4.0.0.13	CARBOHYD	30	30	0.00580307	.	.
PGIP11	netOGlyc-4.0.0.13	CARBOHYD	35	35	0.0185906	.	.
PGIP11	netOGlyc-4.0.0.13	CARBOHYD	40	40	0.11448	.	.
PGIP11	netOGlyc-4.0.0.13	CARBOHYD	66	66	0.352246	.	.
PGIP11	netOGlyc-4.0.0.13	CARBOHYD	71	71	0.0427301	.	.
PGIP11	netOGlyc-4.0.0.13	CARBOHYD	85	85	0.0358737	.	.
PGIP11	netOGlyc-4.0.0.13	CARBOHYD	90	90	0.0226766	.	.
PGIP11	netOGlyc-4.0.0.13	CARBOHYD	95	95	0.138977	.	.
PGIP11	netOGlyc-4.0.0.13	CARBOHYD	96	96	0.17709	.	.
PGIP11	netOGlyc-4.0.0.13	CARBOHYD	100	100	0.0908665	.	.
PGIP11	netOGlyc-4.0.0.13	CARBOHYD	103	103	0.0885722	.	.
PGIP11	netOGlyc-4.0.0.13	CARBOHYD	109	109	0.204119	.	.
PGIP11	netOGlyc-4.0.0.13	CARBOHYD	118	118	0.203041	.	.
PGIP11	netOGlyc-4.0.0.13	CARBOHYD	128	128	0.173504	.	.
PGIP11	netOGlyc-4.0.0.13	CARBOHYD	134	134	0.472943	.	.
PGIP11	netOGlyc-4.0.0.13	CARBOHYD	139	139	0.0738467	.	.
PGIP11	netOGlyc-4.0.0.13	CARBOHYD	147	147	0.0524109	.	.
PGIP11	netOGlyc-4.0.0.13	CARBOHYD	149	149	0.0726275	.	.
PGIP11	netOGlyc-4.0.0.13	CARBOHYD	152	152	0.132728	.	.
PGIP11	netOGlyc-4.0.0.13	CARBOHYD	163	163	0.0380571	.	.
PGIP11	netOGlyc-4.0.0.13	CARBOHYD	171	171	0.0266339	.	.
PGIP11	netOGlyc-4.0.0.13	CARBOHYD	176	176	0.0462855	.	.
PGIP11	netOGlyc-4.0.0.13	CARBOHYD	181	181	0.0999033	.	.
PGIP11	netOGlyc-4.0.0.13	CARBOHYD	182	182	0.0868784	.	.
PGIP11	netOGlyc-4.0.0.13	CARBOHYD	184	184	0.0426701	.	.
PGIP11	netOGlyc-4.0.0.13	CARBOHYD	187	187	0.0665464	.	.
PGIP11	netOGlyc-4.0.0.13	CARBOHYD	191	191	0.421033	.	.
PGIP11	netOGlyc-4.0.0.13	CARBOHYD	200	200	0.164029	.	.
PGIP11	netOGlyc-4.0.0.13	CARBOHYD	206	206	0.164803	.	.
PGIP11	netOGlyc-4.0.0.13	CARBOHYD	209	209	0.0479004	.	.
PGIP11	netOGlyc-4.0.0.13	CARBOHYD	216	216	0.1674	.	.
PGIP11	netOGlyc-4.0.0.13	CARBOHYD	220	220	0.063071	.	.
PGIP11	netOGlyc-4.0.0.13	CARBOHYD	225	225	0.0996527	.	.
PGIP11	netOGlyc-4.0.0.13	CARBOHYD	231	231	0.206716	.	.
PGIP11	netOGlyc-4.0.0.13	CARBOHYD	243	243	0.056921	.	.
PGIP11	netOGlyc-4.0.0.13	CARBOHYD	252	252	0.0485697	.	.
PGIP11	netOGlyc-4.0.0.13	CARBOHYD	260	260	0.153564	.	.
PGIP11	netOGlyc-4.0.0.13	CARBOHYD	261	261	0.120446	.	.
PGIP11	netOGlyc-4.0.0.13	CARBOHYD	267	267	0.265011	.	.
PGIP11	netOGlyc-4.0.0.13	CARBOHYD	270	270	0.288311	.	.
PGIP11	netOGlyc-4.0.0.13	CARBOHYD	276	276	0.0311333	.	.
PGIP11	netOGlyc-4.0.0.13	CARBOHYD	282	282	0.178516	.	.
PGIP11	netOGlyc-4.0.0.13	CARBOHYD	283	283	0.125604	.	.
PGIP11	netOGlyc-4.0.0.13	CARBOHYD	286	286	0.226361	.	.
PGIP11	netOGlyc-4.0.0.13	CARBOHYD	295	295	0.244758	.	.
PGIP11	netOGlyc-4.0.0.13	CARBOHYD	297	297	0.083817	.	.
PGIP11	netOGlyc-4.0.0.13	CARBOHYD	303	303	0.194423	.	.
PGIP11	netOGlyc-4.0.0.13	CARBOHYD	314	314	0.040665	.	.
PGIP11	netOGlyc-4.0.0.13	CARBOHYD	335	335	0.0508127	.	.
PGIP11	netOGlyc-4.0.0.13	CARBOHYD	345	345	0.0430297	.	.
PGIP11	netOGlyc-4.0.0.13	CARBOHYD	348	348	0.00688606	.	.
PGIP11    netOGlyc-4.0.0.13  CARBOHYD	 349	   349	     0.0496855        	.	._____________


Legend
```

^a^ #seqname, The *G. max* PGIP protein under study

^b^ source, The program used to determine *O*-glycosylation

^c^ feature, The studied parameter (*O*-glycosylation)

^d^ start, The starting amino acid position along the PGIP polypeptide chain.

^e^ end, The ending amino acid position along the PGIP polypeptide chain.

^f^ score, The probability score that O-glycosylation occurs at the site.

^g^ strand, DNA strand (not applicable in the analyses presented here)

^h^ frame, The codon frame (not applicable in the analyses presented here)

^i^ comment Whether O-glycosylation is predicted to occur (POSITIVE, if so)
